# The SARS-CoV-2 Spike protein induces long-term transcriptional perturbations of mitochondrial metabolic genes, causes cardiac fibrosis, and reduces myocardial contractile in obese mice

**DOI:** 10.1101/2023.01.05.522853

**Authors:** Xiaoling Cao, Vi Nguyen, Joseph Tsai, Chao Gao, Yan Tian, Yuping Zhang, Wayne Carver, Hippokratis Kiaris, Taixing Cui, Wenbin Tan

**Affiliations:** Department of Cell Biology and Anatomy, School of Medicine, University of South Carolina, Columbia, South Carolina, 29209, USA; Department of Surgery, Division of Otolaryngology-Head and Neck Surgery, UC San Diego School of Medicine, San Diego, CA, 92093, USA; Department of Obstetrics and Gynecology, Xiangya Hospital, Central South University, Changsha, Hunan, 410008, China; Department of General Surgery, The 3rd Xiangya Hospital of Central South University, Changsha, Hunan, 410013, China; Department of Biomedical Engineering, College of Engineering and Computing, University of South Carolina, Columbia, South Carolina, 29208, USA; Drug Discovery & Biomedical Sciences, College of Pharmacy, University of South Carolina, Columbia, South Carolina, 29208, USA

**Keywords:** COVID-19, cardiomyopathy, post-acute sequelae of CoV-2, obesity, mitochondria, respiratory chain complex, ATP synthases, NDUF

## Abstract

**Background:** As the pandemic evolves, post-acute sequelae of CoV-2 (PACS) including cardiovascular manifestations have emerged as a new health threat. This study aims to study whether the Spike protein plus obesity can exacerbate PACS-related cardiomyopathy.

**Methods:** A Spike protein-pseudotyped (Spp) virus with the proper surface tropism of SARS-CoV-2 was developed for viral entry assay *in vitro* and administration into high fat diet (HFD)-fed mice. The systemic viral loads and cardiac transcriptomes were analyzed at 2 and 24 hrs, 3, 6, and 24 weeks post introducing (wpi) Spp using RNA-seq or real time RT-PCR. Echocardiography was used to monitor cardiac functions.

**Results:** Low-density lipoprotein cholesterol enhanced viral uptake in endothelial cells, macrophages, and cardiomyocyte-like H9C2 cells. Selective cardiac and adipose viral depositions were observed in HFD mice but not in normal-chow-fed mice. The cardiac transcriptional signatures in HFD mice at 3, 6, and 24 wpi showed systemic suppression of mitochondria respiratory chain genes including ATP synthases and nicotinamide adenine dinucleotide:ubiquinone oxidoreductase gene members, upregulation of stress pathway-related crucial factors such as nuclear factor-erythroid 2-related factor 1 and signal transducer and activator of transcription 5A, and increases in expression of glucose metabolism-associated genes. As compared with the age-matched HFD control mice, cardiac ejection fraction and fractional shortening were significantly decreased, while left ventricular end-systolic diameter and volume were significantly elevated, and cardiac fibrosis was increased in HFD mice at 24 wpi.

**Conclusion:** Our data demonstrated that the Spike protein could induce long-term transcriptional suppression of mitochondria metabolic genes and cause cardiac fibrosis and myocardial contractile impairment, providing mechanistic insights to PACS-related cardiomyopathy.

## 1. Introduction

Coronavirus disease 19 (COVID-19) is caused by the severe acute respiratory syndrome coronavirus 2 (SARS-CoV-2), a positive-sense single-stranded RNA virus [1]. Viral entry is mediated through the binding of angiotensin converting enzyme 2 (ACE2) with the Spike protein [2]. In addition, neuropillin (NRP) 1 has been identified as an additional host factor for viral entry, which is thought to be ACE2-independent [3, 4]. The Spike protein alone can induce multiple intracellular pathologies. It can increase levels of the hemeoxygenase-1 in kidney cell lines [5], impair endothelial mitochondria functions through downregulation of ACE2 [6], result in natural killer cell-reduced degranulation in lung epithelial cells [7], and disrupt human cardiac pericytes function through CD147-receptor-mediated signaling [8]. We recently have found that the Spike protein impairs lipid metabolism and increases susceptibility to lipotoxicity [9].

Metabolic-associated preconditions such as diabetes mellitus (DM), cardiovascular disorders (CVD), hypertension, and obesity are risk factors for patients with COVID-19 to develop severe symptoms [10]. Our studies have shown decreased levels of total cholesterol (TC), low density lipoprotein cholesterol (LDL-c) and high density lipoprotein cholesterol (HDL-c) in patients with COVID-19, which are associated with disease severity and mortality [11x2013;14]. Mechanistically, lipids have been shown to be a critical contributor to transmission, replication, and transportation for coronaviruses. For example, lipid rafts have been reported to be necessary for SARS virus replication [15]. SARS-CoV-2 Spike protein confers a hydrophobic binding pocket of free fatty acid (FFA), linoleic acid (LA) and cholesterols [16, 17]. The Spike protein alone can impair lipid metabolism in host cells [9]. These lines of evidence have demonstrated that lipids and obesity are important mechanistic factors for COVID-19 severity.

Incomplete recovery has been reported in many COVID-19 patients who have persistent symptoms months beyond the acute stage. The post-acute sequelae of CoV-2 (PACS) or long-COVID is defined as persistence of symptoms and/or long-term complications beyond 4 weeks of SARS-CoV-2 infection or onset of symptoms [18, 19]. Symptoms of PACS have a diverse range with multiorgan involvement [18, 19]. Particularly, CVD sequelae are among major manifestations of PACS which may include dyspnea, palpitation, chest pain, myocardial fibrosis, arrhythmias, and increased cardiometabolic demand [18, 19]. In addition, COVID-19 survivors have an increased risk of incident CVD spanning multiple categories within one year [20]. Obesity is a major risk factor for PASC [21]. Particularly, microvascular injuries caused by SARS-CoV-2 infection are thought to be among the contributive factors to cardiac sequelae of PACS. However, the underlying mechanisms have not been determined.

## 2. Methods

### 2.1 Materials

Human dermal microvascular endothelial cells (hDMVECs) and endothelial cell (EC) culture medium were purchased from ScienCell (San Diego, CA, USA). Phoenix cells and Dulbecco’s Modified Eagle Medium (DMEM) were purchased from ATCC (Manassas, VA, USA). Lentiviral vector pLV-mCherry were obtained from Addgene (Watertown, MA, USA). Coding sequence of SARS-CoV-2 Spike gene (Wuhan variant, GenBank: QHU36824.1) and the expression vector were described previously [22]. LDL-c, HDL-c and lipoprotein depleted fetal bovine serum (LD-FBS) were obtained from Kalen Biomedical, LLC (Germantown, MD, USA). Sulfo-NHS-LC-biotin and desalting spin column were obtained from ThermoFisher (Waltham, MA, USA). The sources of antibodies were listed in the supplementary table 1. Primers were synthesized by IDT (Coralville, IA, USA) and listed in supplementary table 2. The animal protocol was approved by the IACUC committee at the University of South Carolina, Columbia. C57BL/6J wild type and LDL receptor (LDLR) KO (B6.129S7-Ldlr^tm1her^/J) mice were purchased from the Jackson Laboratory (Bar Harbor, ME, USA). The generation of Spp virus was previously reported [22].

### 2.2 In vitro LDL-c and Spp lentivirus binding assay

Human LDL-c and HDL-c were biotinylated using Sulfo-NHS-LC-biotin and purified using a 10Kd-cut off desalting column (ThermoFisher, Waltham, MA, USA). The biotin-LDL-c (2 μg) or biotin-HDL-c (2 μg) was incubated with different quantities of Spp virus (1 to 32 million particles) for 1 hour at room temperature in PBS. The biotin-LDL-c/lentivirus or biotin-HDL-c/lentivirus complex were pulled down and washed three times. The virus bound with biotin-LDL-c or HDL-c was quantified using real time RT-PCR to determine the copy of mCherry gene.

### 2.3 LDL-c-mediated Spp viral cell entry assay

C57BL/6J wild type and LDLR KO (B6.129S7-Ldlr^tm1her^/J) mice were intraperitoneally injected by 3% brewer thioglycollate medium (Millipore-Sigma, St. Louis, MO, USA). Mouse peritoneal cavity elicited macrophages (MØ) were collected, isolated, and cultured in serum-free DMEM/F12 medium for overnight prior to Spp viral entry assay. Primary hDMVECs were cultured in EC medium with 5% FBS. Cardiomyocytes-like H9C2 cells (ATCC, Manassas, VA, USA) were cultured in DMEM (10% FBS) medium. Both hDMVECs and H9C2 cells are plated to 60% confluence and were incubated with 2% LD-FBS (cholesterol levels <0.8 μg/mL) in the medium for overnight prior to assays. Spp virus (4.8×10^7^ particles) was incubated with human LDL-c (12.5 μg) for one hour at room temperature. The Spp lentivirus alone or Spp/LDL-c mixture containing the same quantity of viral particles were added into cells and incubate through 30 minutes to 16 hours. The cells were washed by PBS three time and RNAs were extracted using a Zymo RNA extraction kit. The uptake of Spp virus by cells was determined using a real time RT-PCR. Beta-actin was used as an internal amplification control for normalization. For inhibitory experiments, a scavenger receptor class B type 1 (SRB1) antagonist, block lipid transport-1(BLT-1), was added into the cells to a final concentration of 10 μM for 2 hrs prior to addition of the Spp virus.

### 2.4 Viral administration in vivo

C57BL/6J wild type male mice (8 weeks old) from the same cohort were obtained from the Charles River Labs (Wilmington, MA, USA) and fed normal chows or high fat diet (HFD) (40% fat, TD.95217, Envigo, Indianapolis, IN, USA) for 5 months. In order to achieve 80% power to detect a standardized effect size (means/standard deviation) of 0.80 at alpha = 0.05 level, 6 animals per group were randomly assigned for normal-chow-fed (NCF) mice using the simple randomization method. For HFD mouse groups, 10-12 animals per group were randomly assigned. About 1/3 of mice were unsuccessful to gain weights under HFD thus were removed from the study. We used male mice in this study to minimize the potential hormonal effects on inflammatory responses induced by Spp virus. The Spp virus (8×10^8^ particles) was intravenously administered. The animals were continuously fed normal chows or HFD until the endpoints of the experiments. The animals were sacrificed at 2 or 24 hours post-introducing (hpi) Spp, 3 or 4 days post introducing (dpi) Spp, 3, 6, or 24 weeks post introducing (wpi) Spp and perfused by 50 ml PBS per mouse. The tissues including lungs, heart, liver, kidney, aorta, adipose tissue, and spleen were collected. In general, one part of tissue was used for RNA extraction followed by a real time RT-PCR to determine the number of viral particles in each tissue. The other part was fixed, embedded, and used for histology and immunohistochemistry. Male mice were used in this study to minimize the potential hormonal effects on inflammatory responses induced by Spp virus.

### 2.5 RNA-seq

The cardiac RNA-seq was carried out in the following groups of both normal chow and HFD fed mice: control (prior to viral administration), 24 hpi and 3 wpi. The total cardiac RNAs (n=3 animals per group) were extracted using a Qiagen RNA isolation kit. The ribosomal RNAs were depleted using a Qiagen rRNA HMR kit. RNA-seq libraries were constructed using a Qiagen Standard RNA Library kit followed by the manufacture manual. The insertion size of the library was determined using an Agilent 2100 Bioanalyzer system (Santa Clara, CA, USA). Library quantification was carried out using a NGS library quantification kit from Takara Bio. NGS was performed on a NovaSeq 6000 system (Illumina, San Diego, CA, USA) for 25 M reads per library. The raw reads were aligned and mapped to mouse reference genome using STAR [23]. Validation and quantification of RNA transcripts were performed using FeatureCounts [24]. Biostatistical analysis for gene differential expression (DE) among groups and data graphic generation were carried out in the Rstudio package (https://www.rstudio.com/). False discovery rate (FDR) < 0.05 was considered significant.

### 2.6 Echocardiography

The HFD mice at 24 wpi and age-matched HFD control mice (13 months old) were anesthetized and echocardiography was performed using the VisualSonics Vevo 2100 system (VisualSonics, Inc.) with a previously reported procedure [25].

### 2.7 Statistical analyses

All statistical analysis (except RNA-seq) were performed in Origin 2019 (Northampton, MA, USA). The student two sample *t* test was used for two groups comparisons test. The data was presented as “mean ± s.d.” and *p* < 0.05 was considered as significant.

## 3. Results

### 3.1 LDL-c binds to Spp and facilitates viral entry via SR-B1

In order to test whether Spike protein could be directly associated with lipoprotein cholesterols, we performed a binding assay between LDL-c or HDL-c and Spp virus. The Spp virus bound with biotin-LDL-c or biotin-HDL-c was quantified. Spp virus showed two times maximal binding affinity with LDL-c than HDL-c (Fig 1A). LDL-c showed ten times higher binding capacity with Spp than a regular VSV-G lentivirus (Fig 1A). HDL-c showed similar binding capacities with Spp to VSV-G lentivirus (Fig.1A), suggesting that HDL-c is non-selective to both types of surface glycoproteins on viruses.

**Fig 1.**
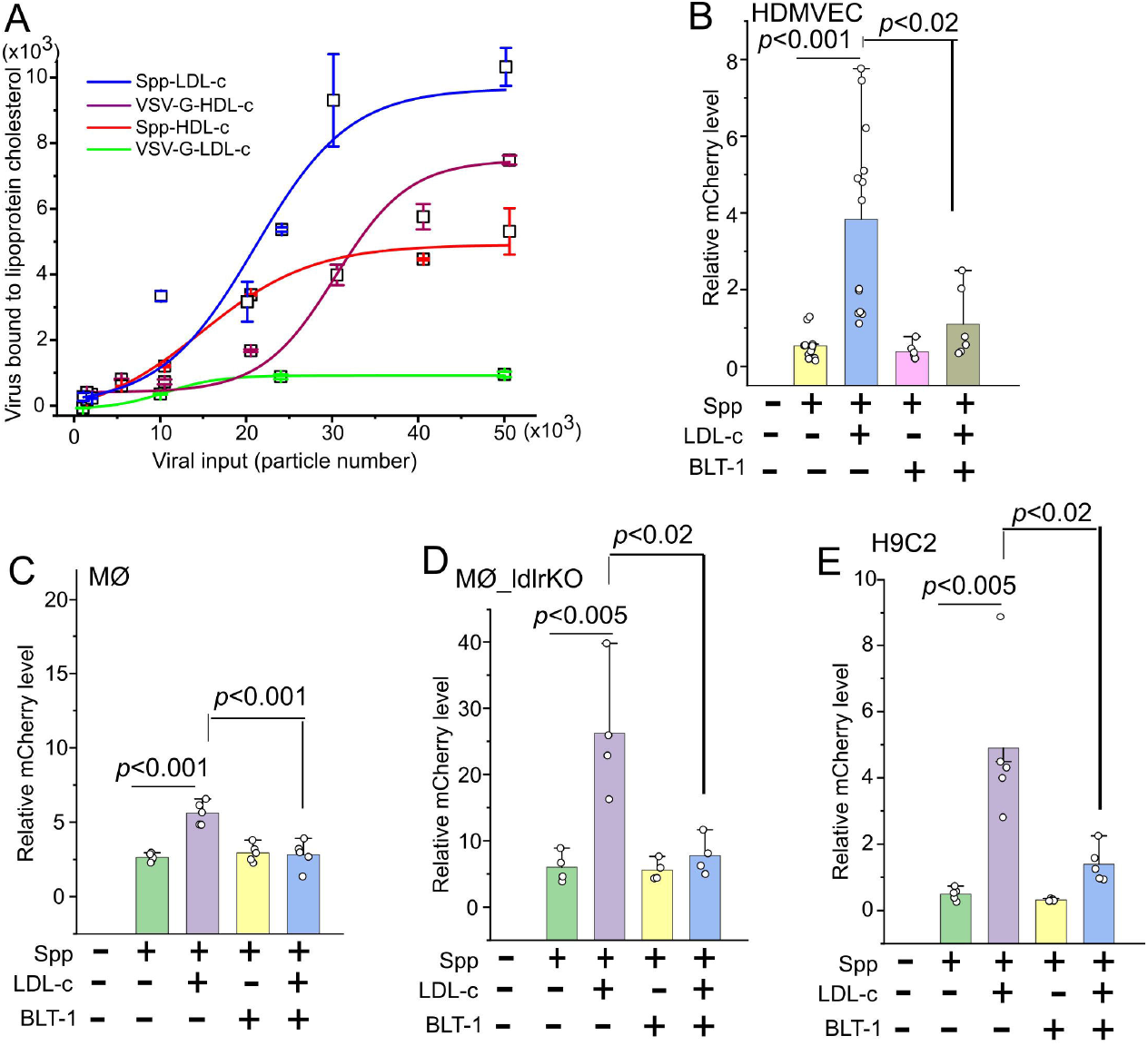
LDL-c augments Spp viral entry which can be blocked by a SR-B1 inhibitor BLT-1. (A) Spp virus shows a higher maximal binding affinity to LDL-c than HDL-c. The biotinylated LDL-c and HDL-c were incubated with various quantities of Spp virus. The lipoprotein cholesterol bound virus was determined by mCherry RNA copies. (B-E) LDL-c significantly increases cellular uptake of Spp virus in HDMVEC (B), MØ (C), LDLr^-/-^ MØ (D), and H9C2 at 2 hpi (E) (*p*<0.001 or 0.005). The addition of BLT-1 significantly blocks the LDL-c augmented Spp viral uptake in HDMVEC (B), MØ (C), LDLr^-/-^ MØ (D), and H9C2 (E) at 2 hpi (*p*<0.02 or 0.001).

The binding preference of Spp to LDL-c let us focus on the potential role of LDL-c in facilitation of Spp cell entry in cardiovascular system. The HDMVECs, mouse peritoneal cavity elicited MØ, and cardiomyocyte-like H9C2 cells were used in this assay. The cellular uptake of Spp virus was significantly increased in the presence of LDL-c in 0.5 to 2 hrs in HDMVECs, 2 to 6 hrs in MØ, and 2 hrs in H9C2 cells as compared with Spp alone (Fig 1B-E, Supplementary Fig 1). A recent report has shown that SARS-2 can bind to HDL-c and enter into cells via SR-B1 [26]. We next investigated whether SR-B1 mediated LDL-c-enhanced Spp uptake in cells. We found that the SR-B1 inhibitor BLT-1 could significantly block the LDL-c-augmented Spp entry in HDMVEC, MØ, and H9C2 (Fig 1B, C, and E). Interestingly, uptake of Spp in LDLR^-/-^ MØ showed a substantial increase in the presence of LDL-c, which could be fully blocked by BLT-1 (Fig 1D). In addition, an anti-LDLR antibody failed to block the LDL-c-augmented Spp entry in HDMVEC (data not shown). This data demonstrates that SR-B1, but not LDLR, accounts for LDL-c-enhanced Spp cellular uptake.

### 3.2 Selective cardiac viral accumulation in obese mice

The timeline of the animal study protocol was shown in Fig 2A. The serum levels of total cholesterol, LDL-c/VLDL-c, and HDL-c were significantly elevated in HFD mice as compared with the mice fed by NCF (Fig 2B). The protein levels of SR-B1 were significantly increased in heart, adipose tissue and kidney, but not in liver, lung and spleen, in HFD mice as compared with NCF mice (Fig 2C, Supplementary Fig 2). In our previous report, we have shown that Spp entered various tissues including adipose tissues, the heart, lungs, the liver, kidneys and the spleen in normal chow-fed mice with the highest level in the lungs at 2 hpi [22]. At 24 hpi, the Spp was effectively removed from these tissues except adipose tissues in which it was instead further increased (*, *p*<0.05, Fig 2B). Intriguingly, the pattern of Spp viral trafficking was shifted in HFD-fed mice, which was characterized by a selective increase in the heart, kidney, aorta and adipose tissues at 2 hpi and remained accumulation in those tissues except kidney at 24 hpi (&, *p*<0.05, Fig 2D). Since the Spp virus was replication incompetent, the viral particles were rapidly cleared in mice and undetectable 3~4 dpi in both NCF and HFD-fed mice using real time RT-PCR assay (data not shown), which could be equivalent to the viral-free stage in discharged COVID-19 patients. The timelines of 3 and 6 wpi could be used as animal models to mimic cardiac sequelae in PASC as we studied their cardiac transcripts below. At 2 and 24 hpi in HFD mice, the cellular locations of Spp virus could be detected in cardiac capillaries and big caliber vessels using co-immunostaining of anti-Spike S1 subunit and anti-CD31 antibodies (Fig 2E). A small number of scattered cardiac MRC1^+^ MØ was observed to have an uptake of Spp virus (Fig 2E). Due to the sensitivity of immunoassay, we could hardly detect any cellular location of Spike protein in hearts in NCF mice at 2 and 24 hpi (data not shown), which was consistent with the presence of very low viral RNA levels in hearts in those animals (Fig 2D).

**Fig 2.**
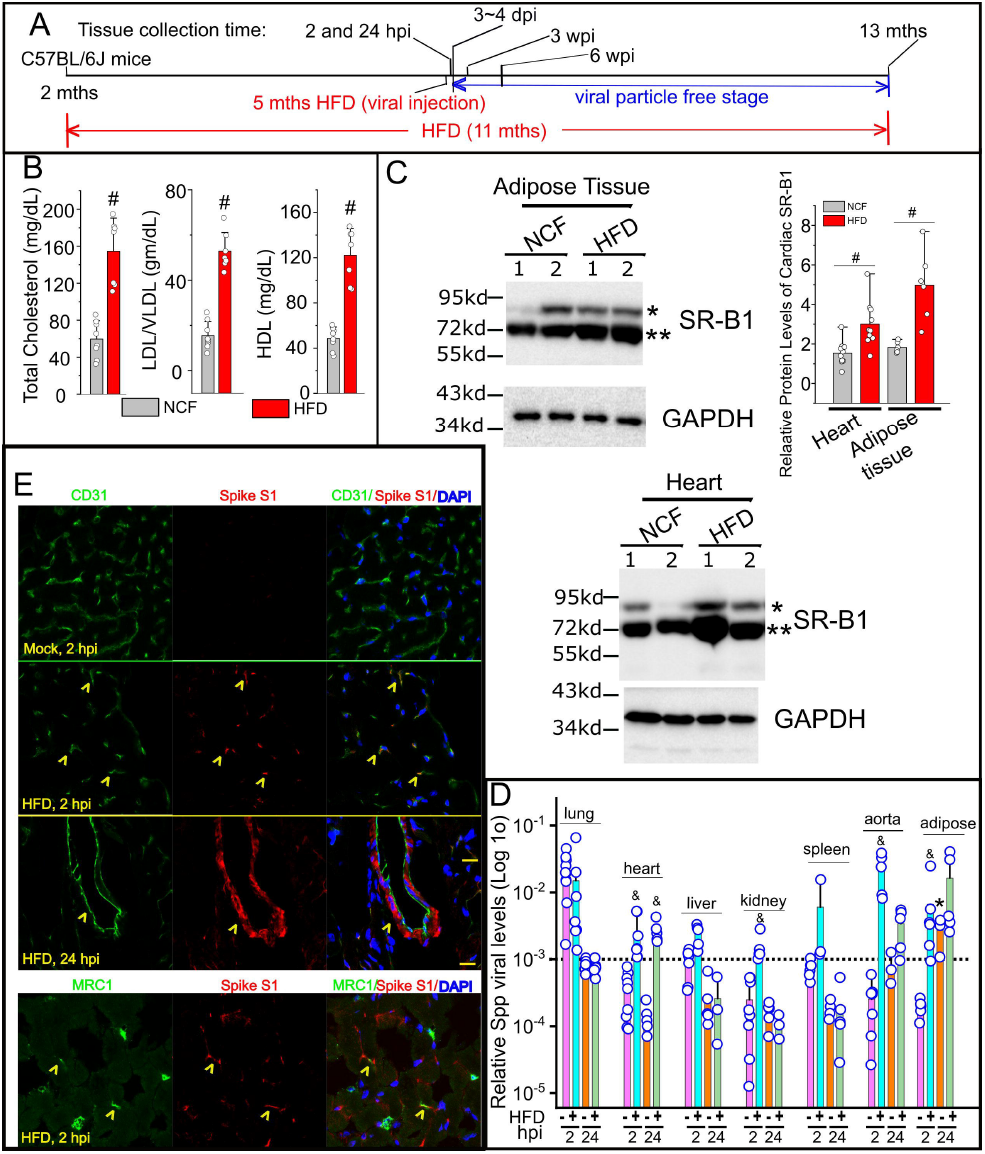
Cardiac Spp viral accumulation in obese mice. (A) Outline of the animal treatment protocol. The C57BL/6J mice (2 months old) were fed normal chow or HFD. The viral administration was given at 3 months later (5 months old). The animals were remained on the same types of diets until being sacrificed at 2 hpi, 24 hpi, 3 wpi, 6 wpi, and 25 wpi. (B) Quantification of serum levels of total cholesterol, LDL/VLDL, and HDL in HFD mice and the NCF group. # indicates *p* < 0.01 for intergroup comparison. (C) The relative protein levels of SR-B1 in hearts and adipose tissues in HFD mice and the NCF group by Western blot. Blots from two animals per group were shown. # indicates *p* < 0.01 in HFD vs NCF group for SR-B1 with the 75Kd MW band (**). (D) Systemic dissemination of Spp virus in HFD mice. Viral load was determined by ratios of viral mCherry levels to housekeeping gene RPS18 levels in each tissue using a real time RT-PCR (Log 10 scale in Y-axis). & indicates statistical significance in HFD group as compared with normal mice group 2 hpi or 24 hpi. Dashed line: the average Spp level in the lungs at 24 hpi. *, indicates statistical significance at 24 hpi as compared with 2 hpi in adipose tissues from normal chow-fed mice. The datasets of various tissues (except adipose tissues) at 2 and 24 hpi from normal-chow-fed mice were adapted from our previous report [22] for a direct comparison to the datasets from HFD mice. (E) Cellular colocalizations of Spike protein of Spp virus in the HFD hearts at 2 and 24 hpi. An anti-Spike S1 subunit antibody (red) is used to recognize Spike protein in Spp. An anti-CD31 or anti-MRC1 antibody (green) is used to stain blood vessels or MØ in heart, respectively. Yellow arrowhead indicates cells positive for both markers. Blue: DAPI staining for nuclei. Scale bar: 20 μm.

### 3.3 Acute cardiac transcriptional responses in obese mice post Spp administration

There were 30 DE genes (FDR<0.05) including 19 upregulated and 11 downregulated transcripts in the NCF mice after 24 hpi as compared with the NCF mice without viral administration (Supplementary Fig 3A & 4A, Supplementary Table 3). This result was consistent with our previous data showing that cardiac viral loads were almost eliminated in NCF mice post 24 hpi [22], which reasonably resulted in very mild transcriptional changes. After 24 hpi, there were total 434 DE genes (FDR<0.05) in the HFD mice as compared with the NCF mice (Supplementary Fig 3B & 4B, Supplementary Table 4). There were total 548 DE genes (FDR<0.05) in the HFD mice after 24 hpi as compared with the HFD control mice without viral administration (Supplementary Fig 3C and D, Supplementary Table 5). The representative transcriptional signatures of downregulation include gene families from ATP synthases (Complex V), reduced nicotinamide adenine dinucleotide (NADH):ubiquinone oxidoreductase family (NDUF) (Complex I) and cytochrome c oxidases (COX) (Complex IV); the representative transcriptional signatures of upregulation include master transcriptional factor of stress signaling, nuclear factor-erythroid 2-related factor 1 (NFE2L1) and signal transducer and activator of transcription 5A (STAT5A), and glucose metabolic gene clusters.

### 3.4 Obesity exacerbates Spp-induced long-term aberrances of cardiac transcriptional signatures

We next focused on the Spp-induced long-term changes in cardiac transcriptomes in obese mice. There were no DE cardiac genes found in NCF mice 3 wpi as compared with the NCF control mice at the level of FDR <0.05 (Supplementary Fig 4C), suggesting that long-term effects on cardiac transcriptome in normal-chow-fed mice were minimal post Spp administration. There were total 209 DE genes (FDR<0.05) including 69 upregulated and 140 downregulated transcripts in the HFD mice 3 wpi as compared with the HFD control mice without Spp administration (Fig 3A, B, Supplementary Table 6). The Gene Ontology (GO) analysis showed the enriched downregulated functional annotations were electron transfer activity and proton transmembrane transporter activity; the enriched upregulated functional annotations were GTPase activity and GTP binding. In particular, three clusters of gene families involving mitochondria respiratory chains (MRC) were significantly downregulated including ATP synthases, NDUFs, and COXs. Transcripts from a total of 33 ATP synthases gene members (including 17 pseudogene members) were detected in HFD mouse hearts; Eleven of them (including 7 pseudogene members) showed significant decrease at 3 wpi (FDR < 0.05, Fig 3C). For NDUF gene family, 9 downregulated DEs out of 35 detected genes were identified; No upregulated DEs were found (FDR < 0.05, Fig 3C). For COX gene family, 3 downregulated DEs out of 23 detected genes were identified (FDR < 0.05).

**Fig 3.**
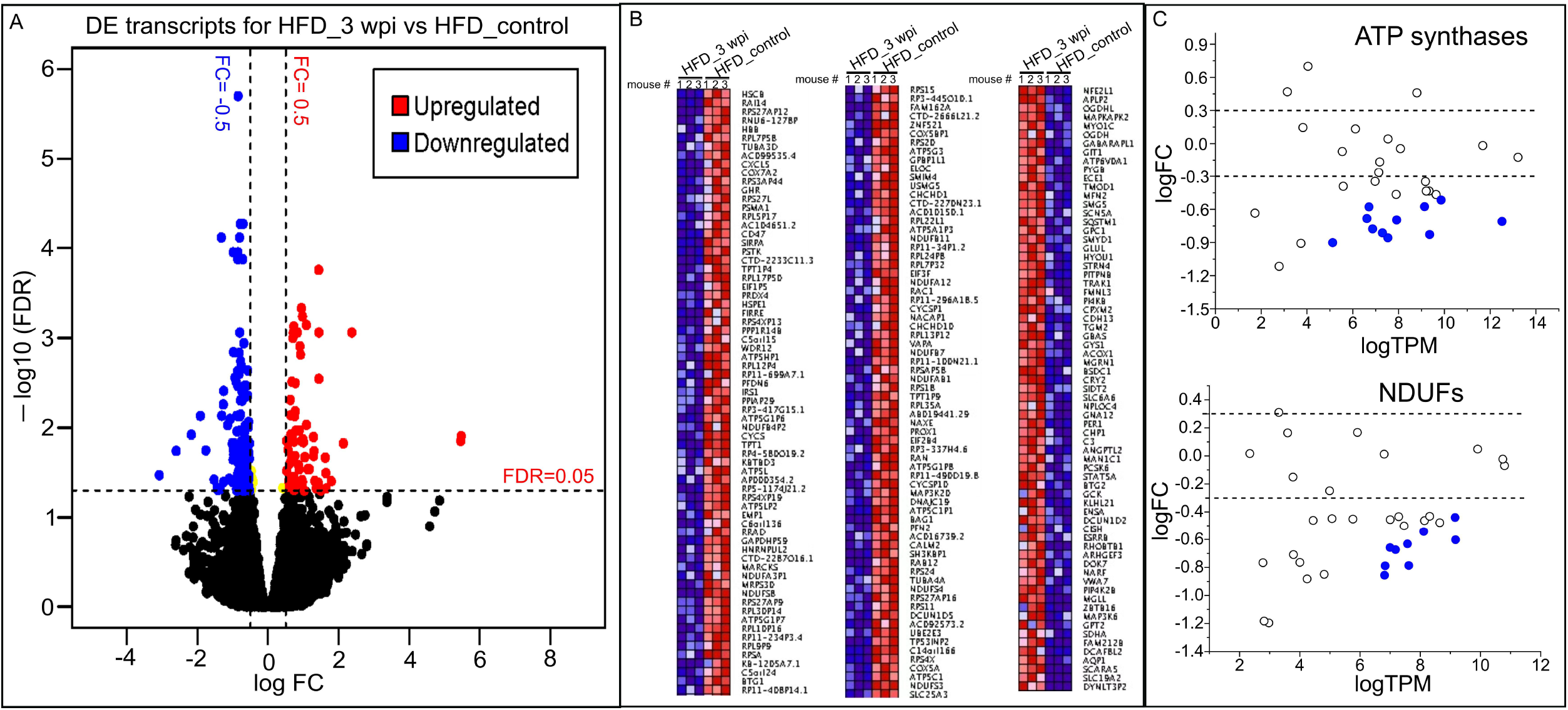
Transient administration of Spp virus induces long-term changes in cardiac transcriptome in obese mice. (A) Volcano plot showing the cardiac DEs in HFD mice 3 wpi as compared with control HFD mice. FDR < 0.05 (horizontal dashed line) is used for cut-off threshold. Right vertical dashed line: fold changes (FC) > 0.5 (increase); left vertical dashed line: FC < − 0.5 (decrease). Blue dots represent downregulated DEs (n=140) and red dots represent upregulated DEs (n=69). (B) Ranked heatmap showing the individual downregulated (left two panels) and upregulated DEs (right panel), which is ranked by FDR. (C) Scatter plot showing significant downregulations of gene families in ATP synthases and NDUFs (FDR < 0.05). Dashed lines in both panels represent FC = ± 0.5 levels. Blue dots represent downregulated DEs in ATP synthase (n = 10 out of 32) or NDUF (n = 9 out of 35) family.

We next compared the cardiac transcripts per million (TPM) of these DE gene members from 24 hpi to 3 wpi in both control and obese mice (Fig 4). In the ATP synthase family, ATP5MK (USMG5), ATP5G3 and ATP5C1P1 in HFD mice at 24 hpi to 3 wpi showed significant decreases as compared with the NCF or HFD control mice (FDR<0.05, Fig 4A). ATP5L, ATP5C1, ATP5G1P6, ATP5G1P7, ATP5LP5, and ATP5G1P8 in HFD mice at 3 wpi showed significant decreases as compared with the NCF or HFD control mice (FDR<0.05, Fig 4A). ATP5A1P3 and ATP5HP1 in HFD mice at 3 wpi showed significant decreases as compared with HFD control mice (FDR<0.05, Fig 4A).

**Fig 4.**
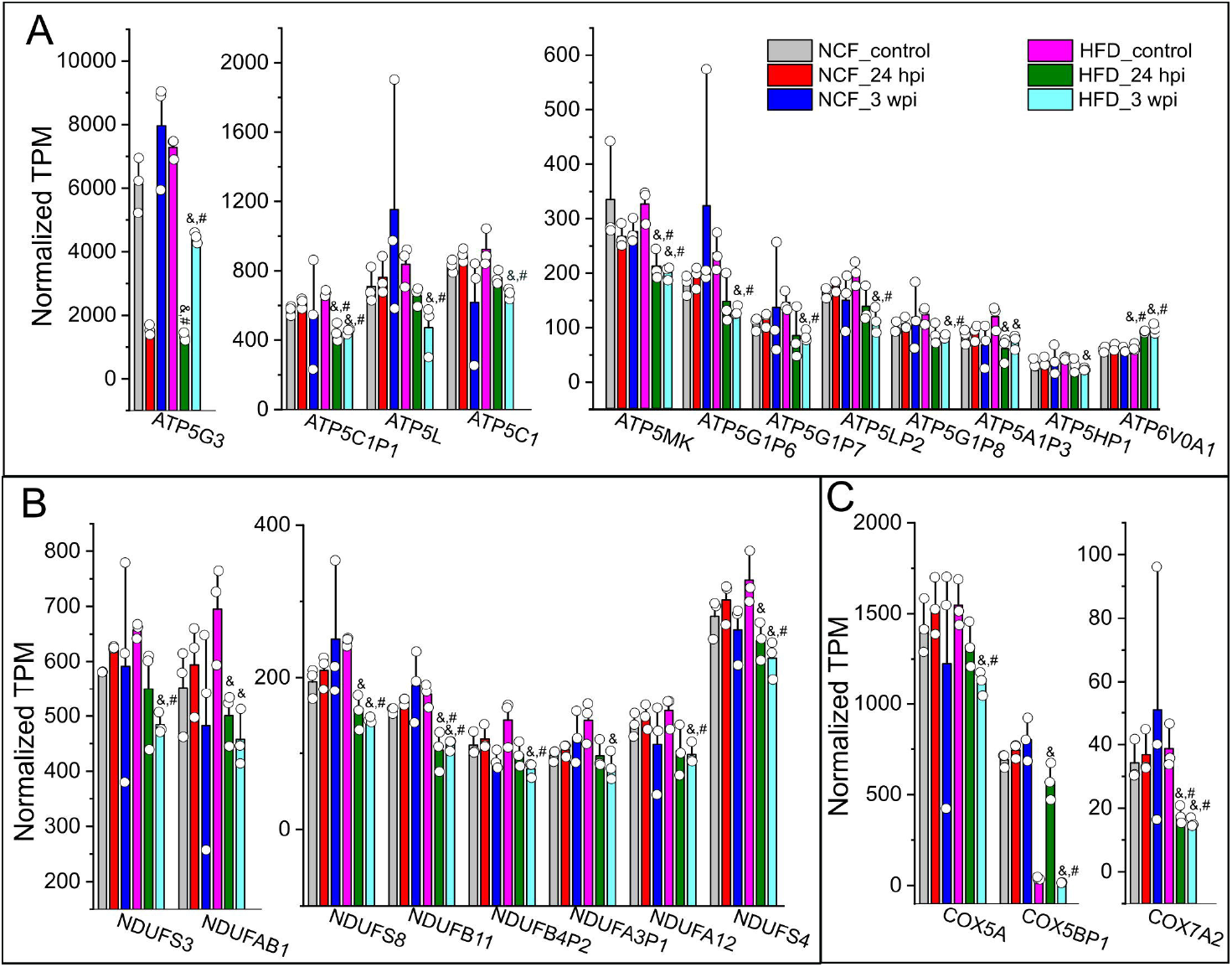
Cardiac DE profiles in mitochondria respiratory chain complex in response to viral administration in obese mice. Normalized TPMs for individual DEs in ATP synthase (A), NDUF (B), and COX (C) families were plotted crossing six sample groups, e.g., NCF mice control or 24 hpi or 3 wpi, HFD mice control or 24 hpi or 3 wpi. The TPMs were extracted from RNA-seq datasets which was analyzed using a Rstudio package. As compared with HFD mouse controls, these genes typically showed significant decreases in HFD mice 3 wpi but not in other groups. &, FDR < 0.05 as compared with HFD mouse control group; #, FDR < 0.05 as compared with NCF control mouse group.

In the NDUF and COX families, NDUFB11 and COX7A2 in HFD mice at 24 hpi or 3 wpi showed significant decreases as compared with the NFC or HFD control mice (FDR<0.05, Fig 5B&C). NDUFS3, NDUFS8, NDUFB4P2, NDUFA12, NDUFS4, COX5A, and COX5BP1 in HFD mice at 3 wpi showed significant decreases as compared with NCF or HFD control mice (FDR<0.05, Fig 4B&C). NDUFAB1 and NDUFA3P1 in HFD mice at 3 wpi showed significant decreases as compared with the HFD control mice (FDR<0.05, Fig 4B&C).

**Fig 5.**
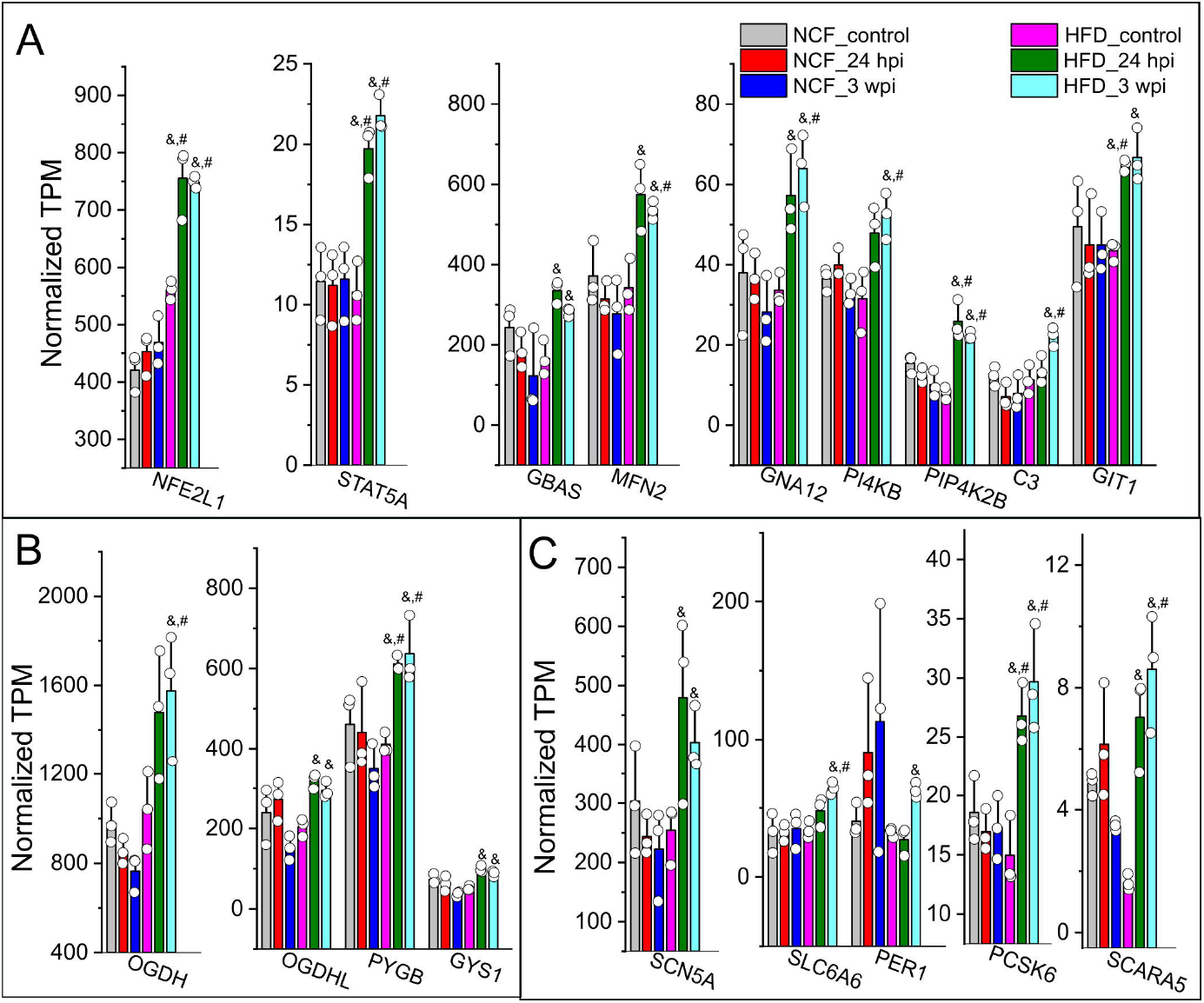
Upregulated cardiac DE profiles in response to viral administration in obese mice. Normalized TPMs for individual DEs in categories of stress pathway / kinase activity (A), glucose metabolism (B), and cardiovascular functions (C) were plotted crossing six sample groups, e.g., NCF mice control or 24 hpi or 3 wpi, HFD mice control or 24 hpi or 3 wpi. The TPMs were extracted from RNA-seq datasets which was analyzed using a Rstudio package. As compared with HFD mouse controls, these genes typically showed significant increases in HFD mice 3 wpi but not in other groups. &, FDR < 0.05 as compared with HFD mouse control group; #, FDR < 0.05 as compared with NCF control mouse group.

There were several functional gene clusters among those 69 upregulated DE transcripts in the HFD mice at 3 wpi as compared with the HFD control mice. The first gene cluster involves stress pathway and kinase activity. NFE2L1, STAT5A and phosphatidylinositol 5-phosphate 4-kinase type-2 beta (PIP4K2B) in HFD mice at 24 hpi or 3 wpi showed significant increases as compared with the NFC or HFD control mice (FDR<0.05, Fig 5A). Mitofusin 2 (MFN2), guanine nucleotide-binding protein subunit alpha-12 (GNA12), phosphatidylinositol 4-kinase beta (PI4KB), and complement component 3 (C3) showed in HFD mice at 3 wpi showed significant increases as compared with the NCF or HFD control mice (FDR<0.05, Fig 5A). Glioblastoma amplified sequence (GBAS) and G protein-coupled receptor kinase-interacting protein 1 (GIT1) in HFD mice at 3 wpi showed significant increases as compared with the HFD mice (FDR<0.05, Fig 5A).

In the gene functional clusters involve in the mitochondria Krebs cycle and cardiovascular functions, glycogen phosphorylase B (PYGB) and proprotein convertase subtilisin/kexin type 6 (PCSK6) in HFD mice at 24 hpi or 3 wpi showed significant increases as compared with the NCF or HFD control mice (FDR<0.05, Fig 5B&C). Oxoglutarate dehydrogenase (OGDH), solute carrier family 6 member 6 (SLC6A6), and scavenger receptor class A member 5 (SCARA5) in HFD mice at 3 wpi showed significant increases as compared with the NCF or HFD control groups (FDR<0.05, Fig 5B&C). Oxoglutarate dehydrogenase L (OGDHL), glycogen synthase 1 (GYS1), sodium voltage-gated channel alpha subunit 5 (SCN5A) and period circadian regulator 1 (PER1) in HFD mice at 3 wpi showed significant increases as compared with the HFD control mice (FDR<0.05, Fig 5B&C).

We next examined the long-term changes of some of those DEs with high abundances (e.g., ATP5L, ATP5G3, NDUFAB1, NDUFA2, NFE2L1, OGDH, PYGB, SCN5A and PER1) in HFD mice using quantitative real time RT-PCR (primers are listed in Supplementary Table 2). The persistent downregulation (ATP5L, ATP5G3, NDUFAB1, and NDUFA2) and upregulation (NFE2L1, OGDH, PYGB, SCN5A and PER1) of those DEs were observed in the hearts of HFD mice at 6 or 24 wpi as compared with the age-matched HFD controls (Fig 6A&B, Supplementary Fig 5). We next examined the cardiac protein levels of some representative DE genes. Consistently with their transcriptional profiles, ATP5C1, NDUFS3, and NDUFA2 showed downregulation while the STAT5A, OGDH, and PYGB exhibited upregulation of their cardiac protein levels in HFD mice at 6 wpi as compared with the NCF or HFD control groups (Fig 6C). Next, we re-examined the existing STAT5A-Cistromes database deposited in the Signaling Pathways Project (http://www.signalingpathways.org/index.jsf) [27–30]. Through these available experimental ChIP-seq data, STAT5A has been found to the common transcriptional factor that binds in the promoters of most of these DE genes (Supplementary table 7), suggesting that STAT5A may act as a master transcriptional factor involving regulation of these DE genes.

**Fig 6.**
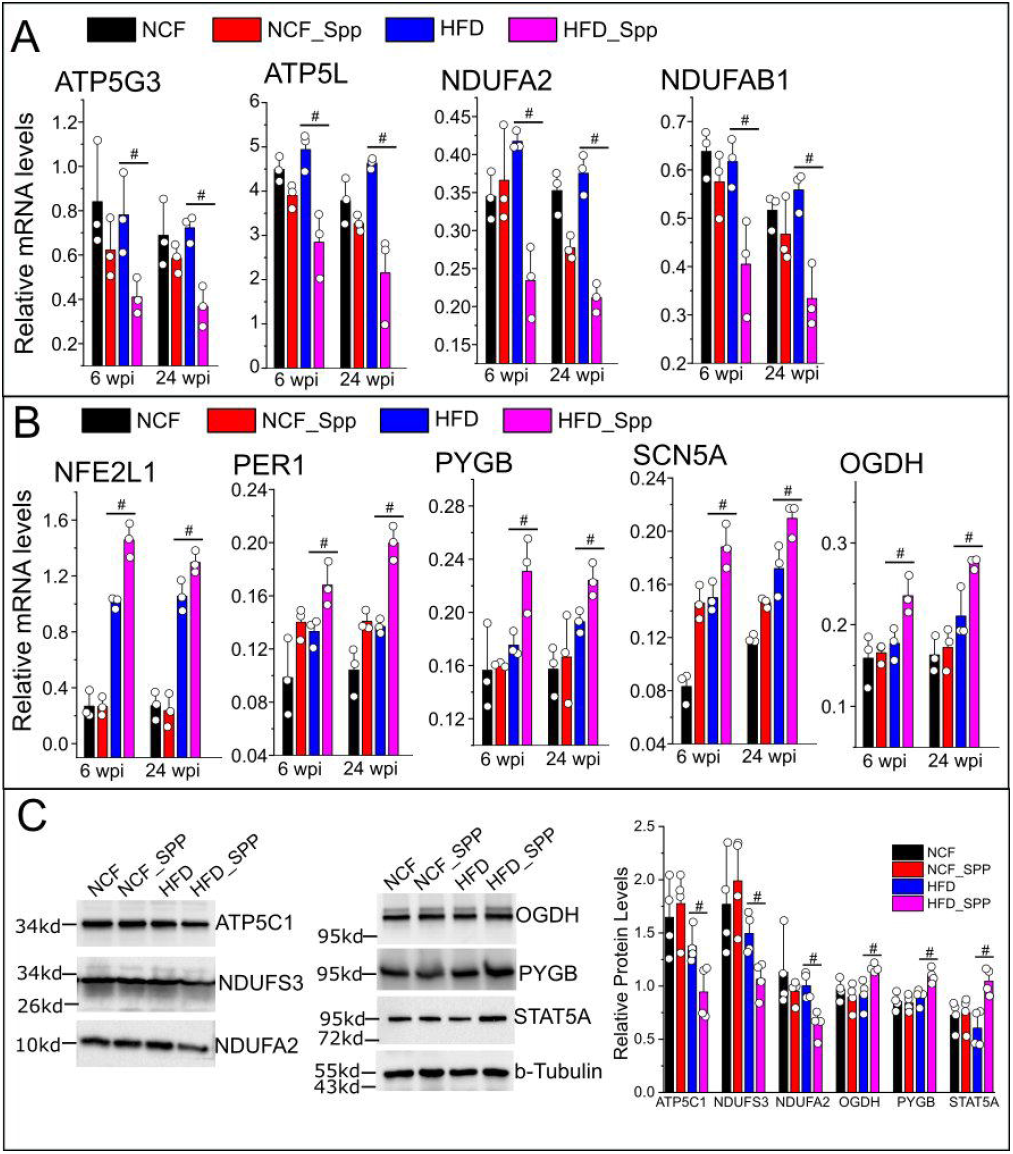
Long-term changes in cardiac DE transcripts in obese mice in response to viral administration. Real time RT-PCR assay was performed to evaluate the relative mRNA levels of representative downregulated (A) and upregulated (B) DE transcripts in the following groups, NCF or HFD mice post Spp administration for 6 or 24 wpi, age-matched NCF or HFD control mouse groups (8.5 or 13 months old). (C) Relative cardiac protein levels of some representative DEs such as ATP5C1, NDUFS3, NDUFA2, OGDH, PYGB, and STAT5A at 6 wpi using Western blot assay. #, *p* < 0.05 as compared with the age-matched control groups with matching ages using two sample *t* test.

Finally, we examined the cardiac fibrosis and *in vivo* cardiac functions. Cross sections of hearts in each group at 6 wpi were examined using Masson’s trichrome staining assay. Except basal level of perivascular fibrosis, there was no obvious fibrosis observed in both NCF and NCF_SPP control groups at 6 wpi (Fig 7A-D, Supplementary Fig 6). Interstitial and perivascular fibrosis were moderately and significantly increased in the HFD group at 6 wpi as compared with the basal levels in NCF and NCF_SPP groups (Fig 7D, Supplementary Fig 6). In the HFD_SPP group at 6 wpi, the development of focal fibrosis was evident, exhibiting significantly higher percentages of cardiac fibrotic areas as compared with the other three groups (Fig 7A-D, Supplementary Fig 6). In order to explore the potential inflammatory mediators for the cardiac fibrosis, we examined the DEs profiles and found consistent upregulations of some inflammatory mediators such as Angiopoietin Like 2 (ANGPTL2), MAPK Activated Protein Kinase 2 (MAPKAPK2) and STAT5A in viral-administered HFD mice versus age-matched HFD controls at both 24 hpi and 3 wpi (Supplementary Tables 5&6). ANGPTL2 is a proinflammatory protein associated with various chronic inflammatory diseases and MAPKs are among downstream signaling targets [31]. Persistent STAT5 activation is known to promote chronic inflammation [32]. We then performed an IHC to investigate the expression patterns of phosphorylated-STAT5A (p-STAT5A) and p-ERK in the cardiac sections. There were many p-STAT5A and p-ERK positive cells in the focal fibrotic areas at 3 wpi in HFD hearts (Supplementary Fig 7). There was no evident p-STAT5A positive cells in the heart sections from the groups of NCF, NCF+Spp, and HFD (Supplementary Fig 7). There was a basal level of p-ERK signaling in scattered cells in the heart sections from the groups of NCF, NCF+Spp, and HFD, but less intensity than in the cardiac fibrotic areas from the HFD+Spp group (Supplementary Fig 7).

**Fig 7.**
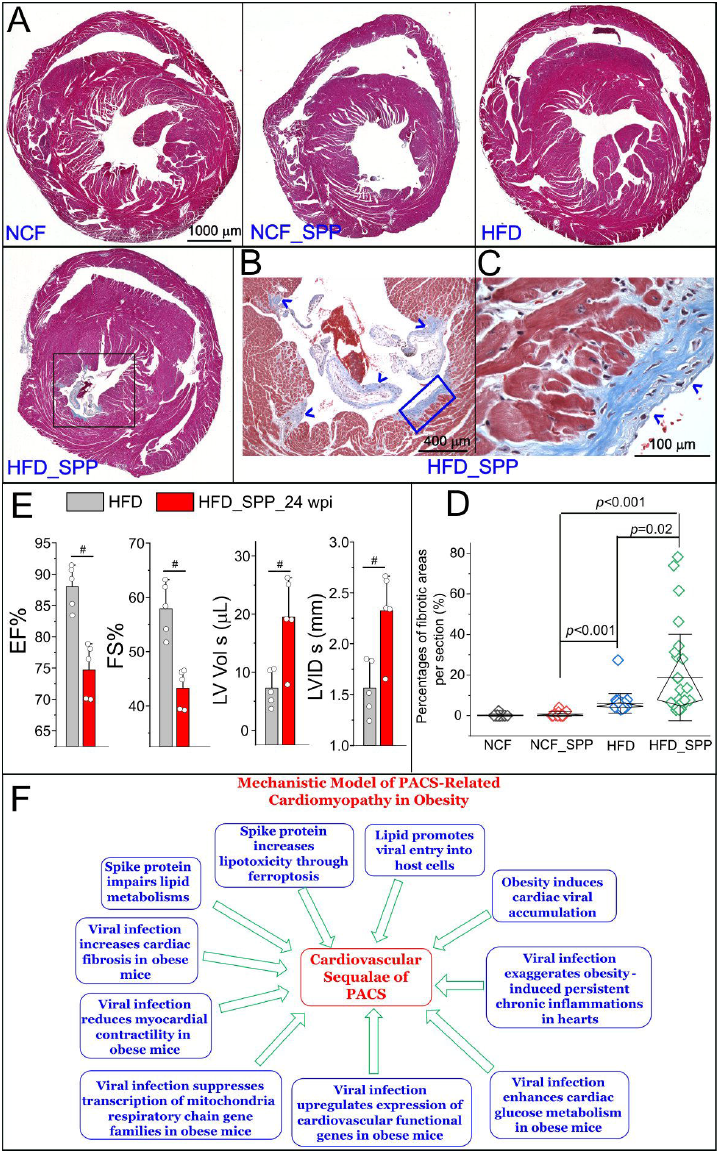
Development of cardiac fibrosis and cardiac functional changes in obese mice in response to viral administration. (A) Masson’s trichrome staining on heart cross sections at 6 wpi. The development of focal fibrosis is evident in the HFD_SPP group (black boxed area). (B) a high magnification of the boxed area in the HFD_SPP group in (A). (C) a high magnification of the boxed area in (B). Blue arrow heads indicate the focal fibrotic areas. (D) Quantitative analysis of the percentage of fibrotic areas in each group. Mann-Whitney test was used to compare the differences among two groups. Whiskers are presented as mean ± S.D. with the diamond boxed showing interquartile ranges of the data. (E) Comparison of echo parameters such as EF (%), FS (%), LV Vol s (μL), and LVID s (mm) between the HFD mice 24 wpi and the age-matched HFD controls (13 months old). #, *p* < 0.05. (F) A proposed mechanistic model for cardiomyopathy in PACS under obese conditions which are likely resulted from multifactorial contributions: spike protein impairs lipid metabolisms and increases lipotoxicity-associated ferroptosis; lipids promote viral administration and facilitate cardiac viral accumulation; and viral administration exaggerates obesity-induced persistent chronic inflammation, enhances glucose metabolism, compensates cardiovascular functions, systemically suppresses expression of MRC gene families, decreases cardiac myocardial contractility, and increases cardiac fibrosis.

Echocardiogram showed that there was a significant decrease in cardiac ejection fraction (EF) (%) and fractional shortening (FS) (%) in HFD mice at 24 wpi as compared with the age-matched HFD controls (Fig 7E). In addition, the left ventricular end-systolic diameter (LVID s) and volume (LV Vol s) in HFD mice at 24 wpi were significantly increased as compared with the age-matched HFD controls (Fig 7E). There were no significant in other parameters between these two groups, including left ventricular end-diastolic diameter (LVID d), left ventricular end-diastolic volume (LV Vol d), left ventricular end-diastolic posterior wall thickness (LVPW d), left ventricular end-systolic posterior wall thickness (LVPW s), left ventricular end-diastolic anterior wall thickness (LVAW d), left ventricular end-systolic anterior wall thickness (LVAW s), and stroke volume (SV), LV mass, cardiac output, body weight, and heart weight-to-tibia length (Supplementary Fig 8). This data together demonstrates a reduction of myocardial contractility, suggesting a development of myocardial damages.

## 4. Discussion

This study demonstrates that the Spike protein can induce a long-term transcriptional suppression of gene families related to mitochondria metabolic pathways as well as facilitate cardiac fibrotic development and myocardial contractility reduction in obese mice. It also shows that SR-B1 is a major contributor to the cholesterol-enhanced viral entry into host cells. This data reveals the cardiac pathological features exacerbated by the Spike protein with obesity, providing novel insights into CVD sequelae of PACS.

SR-B1 is a major receptor for trafficking of HDL-c, ox-LDL-c, and LDL-c and LDLr is the receptor for LDL-c uptake. Both are co-receptors for many pathogens entry of host cells with an aid from cholesterol [33–37], make them the leading candidates of co-factors in host cells such as EC, MØ and cardiomyocytes to mediate viral entry in obesity. SARS-2 S1 subunit can bind to HDL-c and facilitate entry into cells via SR-B1 [26]. In this study, we have shown that SR-B1 is the major receptor for LDL-c or HDL-c-mediated enhancement of viral entry. As the fact that obese patients generally have elevated LDL-c levels and decreased HDL-c levels in the blood [38], we posit that LDL-c is preferably exploited by SARS-CoV-2 for viral entry in obesity. We have shown that cardiac and adipocytic but not pulmonary SR-B1 are increased in HFD-fed mice. This may explain the preferable viral uptake in heart and blood vessels but not in lungs in obese mice, providing insightful information regarding the mechanism that patients with preconditions such as CVD, hypertension, and obesity have high risks to be infected and develop severe symptoms.

The potential roles of cardiac mitochondria perpetuations in CVD sequelae of PACS are yet to be determined. HFD has no effect on membrane potentials, redox profiles, and ATP synthases, but induces increased rates of electron leak, on cardiac mitochondria [39]. Therefore, the longterm transcriptomic changes in MRC genes may be a result from a viral administration and HFD. In addition, STAT5A has been revealed to be dysregulated in response to a viral administration and HFD in this study. Many of the DEs transcripts of MRC genes have a binding motif of STAT5A, suggesting that STAT5A may serve as the master transcriptional factor in controlling their expressions. Our data has been supported by and consistent with various reports in actual infection system or patients. For example, evidence has shown mitochondrial dysfunction, metabolic alterations with an increase in glycolysis, and high levels of mitokine in PBMCs from patients with COVID-19 [40]. Lower levels of mitochondrial membrane potential were found in the elderly who had symptoms suggestive of COVID-19 or with a confirmed diagnosis of COVID-19 [41]. SARS-CoV-2 can cause transcriptional downregulation of mitochondria-related processes, respiratory electron transport chain, and ATP synthesis coupled electron transport in the lung epithelial cell line A549 [42]. Another study has shown that SARS-CoV-2 can downregulate electron transport chain complex I and ATP synthase genes in human airway epithelial cells [43]. Depolarized mitochondria and abnormal mitochondrial ultrastructure have been found in monocytes in patients with COVID-19 [44]. Dysfunctional mitochondria-dependent lipid catabolism is found in the plasma of patients with PASC [45]. This accumulative evidence and our data together have shown that SARS-CoV-2 infection can induce mitochondriopathy in COVID-19. The mitochondria have been identified as an important drug target for the next wave of cardiac drugs since their dysfunction is closely associated with cardiac function declines and heart failure [46]. Therefore, targeting the mitochondrial metabolic pathways may provide a potential new revenue for treatment of cardiovascular-associated symptoms in PASC.

The molecular mechanisms underlying PACS remain elusive. Recent evidence suggests that active viral reservoirs or/and persistent circulating Spike may be associated with PASC symptoms. Persistent circulating Spike protein and/or viral RNA have been found in patients with PASC [47–49]; In addition, the Spike protein is efficiently incorporated into cell membranes [9]and has been found in viral RNA-absent extracellular vesicles in patient blood [48]. This may evade from a recognition by neutralized antibodies, resulting in a persistent and systemic distribution and prolonged lifetime of the Spike protein. Multiple studies have shown that the Spike protein can induce non-infective cardiovascular stress, including disruption of human cardiac pericytes function, vascular endothelial cells, and the blood-brain barrier [6, 8, 50]. We further evidence of this showing that the Spike protein can cause a long-term transcriptional suppression in MRC gene families. This accumulative data together provides an insight into the potential role of the Spike protein in the pathogenesis of PACS.

There are several limitations for this study. (1) The Spp lentivirus cannot completely replicate the pathological process of SARS-CoV-2 in human. However, it can be handled in a BLS2 facility and has been shown to be a very useful system to investigate the Spike protein-mediated cell type susceptibility, host tropism for infection and pathogenicity [22, 51]. Thus, it is a valuable tool to study the potential role of Spike protein in PACS. (2) This study does not simulate the nature of the entry pathway of SARS-CoV-2 through upper airway, which is a limitation. The PACSs are associated with the persistent and chronic presence of extrapulmonary residual viral components through circulatory system [47–49]. For example, an dissemination of SARS-CoV-2 RNA has been found in multiple organs even over 7 months in some patients; particularly, residue viral RNA can be detectable in the cardiovascular system including myocardium, pericardium, endothelium, aorta, and vena cava in 80% patients [52]. Another study has corroborated that the Spike and nucleocapsid proteins of SARS-CoV-2 are present in GI, hepatic tissues, and lymph nodes in recovered patients with COVID-19 over 6 months [53]. This evidence suggests the role of extrapulmonary viral burdens in PACS, which is distinct from the primary nasal infection pathway. Our pseudoviral model only mimics this secondary infection process and the Spike protein associated prolonged pathologies. (3) Our results can only be applicable to the Wuhan variant of Spike protein which is also the first step to enlighten the molecular mechanisms underlying CVD PACS. Clinical evidence has shown CVD PACS symptoms in patients infected by the Wuhan original strain. For example, Palpitation and tachycardia were reported in patients infected with the Wuhan variant in the cohorts from Spain and Italy [54, 55]. In U.S., long-term CVD sequelae was early reported from a cohort in Michigan (6 March to 1 July 2020) [56] and a cohort of veterans (1 March 2020 to 15 January 2021) [20]. Those CVD PACS were posited to be mainly caused by the infection of the Wuhan variant since the alpha variant first appeared late November 2020 [57] and was consisted of 1.0% of total cases by the 2^nd^ week of January 2021 in the U.S. [58]. The evolutionary mutating of the Spike protein has led to several variants of concern (VOC) including Alpha, Beta, Gamma, Delta, and the current Omicron, resulting in less pathogenicity but higher transmissibility [59, 60]. Our data may not reflect the mechanism of CVD PACS caused by the current dominant Omicron strain, which is a limitation for this study. However, the significance of mechanistic studies of CVD PACS caused by the original variant of Spike protein shouldn’t be undervalued as this variant presents a higher pathogenicity than other VOCs. Our data will also strengthen future studies that include the Omicron and other variants to elucidate this mechanism in a systemic spectrum. (4) The function of mitochondria has not been elicited in this model. The detailed role of cardiac mitochondrial perturbations in PACS remains to be determined.

In summary, we have provided an insight to the mechanisms underlying cardiac sequelae of PACS (Fig 7F). First, cholesterols such as LDL-c can enhance viral entry into host cells *in vitro* and obesity can cause selective viral accumulation in heart, aorta, and adipose tissues, demonstrating that lipid and obesity can particularly facilitate the viral burden in cardiovascular system. Second, the Spike protein alone can impair lipid metabolic and autophagic pathways thus augment the lipid overload-induced ferroptosis [9]. Third, consistent chronic inflammations are exacerbated in HFD hearts with addition of viral infection. Forth, the downregulation of transcriptional profiles of MRC gene families indicates a perturbance of mitochondria energy production as a long-term cardiac sequala upon Spp viral burden under obese conditions. Forth, upregulation of glucose metabolic-related genes implies imbalance of cardiac fatty acid oxidation (FAO)-produced ATPs. Fifth, increases in cardiovascular function-related transcripts suggest cardiac functional compensation and restoration in response to chronic inflammation-induced metabolic aberrances in hearts. Sixth, the increased cardiac fibrosis is evident. Seventh, the decreased cardiac EF and FS and increased LVID s and LV Vol s demonstrate reductions of myocardial contractility. This data suggests a development of cardiac damages towards evident functional impairments. Collectively, our data have shown Spike protein induces transcriptional suppression of cardiac mitochondrial metabolic genes, cardiac fibrosis, and reduction of myocardial contractility in obese mice, providing a mechanistic insight to cardiomyopathy sequelae of PACS.

## Abbreviations

ACE2: angiotensin converting enzyme 2
COVID-19: coronavirus disease 2019
CVD: cardiovascular disorders
EF: ejection fraction
FS: fractional shortening
HFD: high fat diet
LDL-c: low-density lipoprotein cholesterol
MRC: mitochondria respiratory chain
NCF: normal-chow-fed
NDUF: nicotinamide adenine dinucleotide:ubiquinone oxidoreductase family
NFE2L1: nuclear factor-erythroid 2-related factor 1
PASC: post-acute sequelae of CoV-2
SARS-CoV-2: severe acute respiratory syndrome coronavirus 2
Spp: spike protein-pseudotyped lentivirus
SRB1: scavenger receptor class B type 1
STAT5A: signal transducer and activator of transcription 5A
TMP: transcripts per million

## CRediT authorship contribution statement

**Xiaoling Cao**,Conceptualization, Animal study Conduct, Data Analysis, Methodology, Project administration. **Vi Nguyen**, Conceptualization, In vitro study Conduct, echocardiography, Data analysis, Methodology. **Chao Gao**, Study Conduct including serum lipid determination and mitochondria staining. **Joseph Tsai**, NGS data analysis; **Yan Tian**, Study Conduct, Data Analysis. **Yuping Zhang**, Study Conduct, Data Analysis. **Wayne Carver**, Writing – review and editing. **Hippokratis Kiaris**, Conceptualization, Writing – review and editing, funding acquisition. **Taixing Cui**, Conceptualization, Writing – review and editing, funding acquisition. **Wenbin Tan**, Conceptualization, Writing – original drafting, review and editing funding acquisition.

## Acknowledgements

We are very thankful to the support and assistance from Instrumentation Resource Facility at University of South Carolina School of Medicine. We are very grateful to the support from Dr. Igor Roninson and the COBRE Center for Targeted Therapeutics at University of South Carolina.

## Database deposition

Gene expression and transcriptome obtained from NGS are deposited into NIH GEO (Gene Expression Omnibus) (https://www.ncbi.nlm.nih.gov/geo/). The raw data (reads) of NGS are deposited into SRA (Sequence Read Archive) (https://www.ncbi.nlm.nih.gov/sra)

