## Supplemental materials for "The SARS-CoV-2 Spike protein induces long-term transcriptional perturbations of mitochondrial metabolic genes, causes cardiac fibrosis, and reduces myocardial contractile in obese mice"

#### Supplementary Methods and Materials:

*1.1 Spp virus generation*

Spp virus was generated in Phoenix cells after being co-transfected by pLV-mCherry and pcDNA-Spike using a calcium phosphate kit (ThermoFisher, Waltham, MA, USA), which was previously reported [1]. The supernatant containing Spp virus was harvested 48 to 72-hours post transfection, clarified by centrifuging at 5,000 g for 15 min, followed by passing through 0.45 μm filter disk, and precipitated by ultracentrifuging at 24,000 rpm for 2 hours. The viral pellets were resuspended in cold PBS buffer, aliquoted, and stored at -80 °C before use. The viral particle number was determined using real time RT-PCR assay. Co-transfection of pLV-mCherry and pMD2.G vector carrying vesicular stomatitis virus glycoprotein (VSV-G) expression cassette were performed to obtain VSV-G lentivirus [1].

*1.2 Real time RT-PCR, Western blot, and Immunostaining*

RNA extraction kit was obtained from Zymo Research (Irvine, CA, USA) and Qiagen (Hildern, Germany). The RT kit was obtained from Takara Bio USA (Mountain view, CA, USA). The SYBR green master mix was from Biorad (Hercules, CA, USA). For cDNA synthesis, RT reaction was carried out at 42°C for 2 hrs containing 1x RT buffer, 1.0~5 μg of total RNA, 0.5 mM dNTPs, 0.5 μg of oligo (dT) 15-mer primer, 20 units of RNasin, and 5 units of SMART Moloney murine leukemia virus reverse transcriptase. A combined index of three house-keeping genes beta-actin, Rps18 and Nono were used as controls to normalize the amplification data in real time PCR. The real time PCRs were performed as we previously reported [1]. For Western Blot assay, cellular proteins were extracted from hearts and other organs using RIPA lysis buffer (Santa Cruz Biotech., Inc., Dallas, TX, USA), separated by SDS-PAGE, followed by a transfer onto PVDF membranes. Primary antibodies were added and followed by HRP-labeled secondary antibodies. Images were acquired using a Bio-Rad Gel Imaging System (Hercules, CA, USA). GAPDH, beta-actin, and beta-tubulin were used as internal controls.

For immunostaining, hearts from mice were under snap freezing process and approximately 6 μm thick fresh sections were cut and collected. The sections were then fixed by 10% buffered formalin (Fisher Scientific, Pittsburgh, PA, USA), blocked with 5% donkey serum, and incubated in a humidified chamber overnight at 4$^{o}$C with primary antibodies, followed by fluorescent conjugated secondary antibodies for 2 hours at room temperature. Images were acquired using ImageXpress Pico System (Molecular Device, San Jose, CA, USA) or confocal microscopy system (Carl Zeiss AG, Oberkochen, Germany).

*1.3 Cardiac fibrosis staining*

The fibrosis in mouse hearts was evaluated using Masson's trichrome staining (#HT15-1KT, Millipore Sigma). Briefly, the heart cross sections were fixed by Formalin and stained in Weigert's iron hematoxylin working solution, followed by Biebrich scarlet-acid fuchsin solution, phosphomolybdic-phosphotungstic acid solution, and Acetic acid. The sections were dehydrated and mounted. The images were acquired using ImageXpress Pico System (Molecular Device, San Jose, CA, USA). Images from 5-8 sections (60x objective) per heart were analyzed using ImageJ software plug in Color deconvolution 2 and the percentage of fibrotic areas per section was estimated (n=3 hearts per group). Mann-Whitney test was used to compare the difference among two groups.

**Supplementary Discussions**:

Statins can upregulate the expression of SR-B1 [2], but such an effect seems unlikely to increase risk of SARS-CoV-2 infection since viral entry can be mitigated by lowering cholesterol levels and altering membrane lipid rafts by statins. Indeed, the use of statins does not alter the SARS-CoV-2 infection in patients [3]. Furthermore, many retrospective observational studies have shown that statin therapy can reduce the risk of severity and mortality and improve prognosis in hospitalized patients with COVID-19 [4, 5]. Mechanistically, statins can attenuate SARS-CoV-2 replication *via* main protease and RNA-dependent RNA polymerase, reduce inflammation, improve endothelial function, decrease risk of thrombosis, and alleviate pulmonary fibrosis in patients with COVID-19 [6, 7]. However, the statin-related side effects such as increase in risk of diabetes may worsen the outcome in inpatients with type 2 diabetes, e.g., elevating COVID-19 related mortality [8].

Pathophysiological mechanisms perpetuating CVD sequelae of PACS are yet to be determined. Both acute and chronic cardiac injuries secondary to SARS-CoV-2 infection are thought to be involved [9]. SARS-CoV-2 virus has been detected in the heart tissue in autopsy studies [10]. Acute infection of SARS-CoV-2 can cause microvascular and myocardial injuries, including endothelialitis, microthrombi, and altered renin-angiotensin homeostasis [11, 12]. In an animal model, SARS-CoV-2 infection results in cardia pericyte loss, fibrosis, cardiomyocyte hypertrophy, and diastolic dysfunction in hamster [13]. The Spike protein alone can damage cardiac pericytes function [14]. In this study, long-term transcriptomic suppression of mitochondria related respiratory genes are revealed after a transient (one time) exposure of the replication-incompetent Spp virus which viral particles are cleared in 3~4 days after administration in obese mice. This data suggests that potential mitochondria malfunction, probably caused by obese-induced inflammatory responses and subsequently exacerbated by viral infections, is a contributor to the long-term cardiac sequelae in COVID-19 patients. The transition from acute cardiac illnesses to chronic phase in PACS may be possibly explained by several putative mechanisms. First is the chronic inflammatory response evoked by SARS-CoV-2 infection or integration of the SARS-CoV-2 genome into host DNA, which can induce a continuous activation of immune-inflammatory cascades and results in persistent endothelial dysfunction and cardiomyopathy [15]. Second, autoimmune antibodies against cardiac antigens are elicited through molecular mimicry in COVID-19, which may contribute to the delayed damages [16].

It has been well established that obesity with impaired storage of fat in adipose tissue forces non-adipose tissues, such as the heart, to keep the excess fat, a phenomenon referred to as ectopic fat deposition, which is usually toxic to the non-adipose tissue, i.e. lipotoxicity, leading to obesity-related complications, such as vasculopathy and cardiomyopathy [17, 18]. It is reasonable to posit that SARS-CoV-2 infection interplays thus exacerbates obesity-related cardiac comorbidities. In this study, we have identified upregulations of clusters of genes involving stress signaling, glucose metabolism, and cardiovascular functions. (1) NFE2L1 is a master transcription factor that regulate expression of all the proteasome subunit genes during stress response [19].

Other upregulated DEs with functional characteristics of inflammatory and kinase activity included GBAS, MFN2, GNA12, PI4KB, PIP4K2B, C3, and GIT1. For example, GBAS is localized to mitochondria and plays a role in oxidative phosphorylation and vesicular transport; it can facilitate NF- κB-mediated cytokine production [20]. *MFN2 is a* GTPase embedded in the outer membrane of the mitochondria, involving mitochondria fusion and mitophagy [21]. *(2) Genes involve in glucose metabolisms, such as OGDH, OGDHL, PYGB, and GYS1. OGDHL* is a rate-limiting enzyme in the Krebs cycle that is pivotal in mitochondrial metabolism [22]. *PYGB* regulates glycogen mobilization [23]. Under normal conditions, main source of cardiac ATP is from fatty acid oxidation (FAO), but less from glucose metabolism. With mitochondria dysfunction or metabolic perturbations, cardiac metabolic profiles are impaired by reduction in FAO and concomitant increases in glucose utilization [24, 25]. (3) Gene clusters engage in cardiovascular functions. SCN5A is a tetrodotoxin-resistant voltage-gated sodium channel subunit and primarily expressed in cardiac muscle for the action potential initiation in an electrocardiogram [26]. SLC6A6 is a sodium and chloride ion-dependent taurine neurotransmitter transporter, which deficiency leads to cardiomyopathy [27]. SCARA5 has a function in scavenger receptor activity and ferritin receptor activity, which genetic locus is associated with the susceptibility to venous thromboembolism [28]. PER1 is a member of the Period family of genes for circadian rhythms, which can regulate cardiovascular rhythm centrally (through brain) and peripherally (in the heart) [29-31]. The upregulation of PER1 is probably due to the long-term stress or chronic inflammatory consequence in HDF mice after viral administration. PCSK6 is upregulated in human atherosclerotic plaques associated with smooth muscle cells (SMCs); it induces SMC migration and modulates vascular remodeling [32]. These data demonstrate that long-term chronic inflammations persist in HFD mice after viral administration, leading to cardiac metabolic and functional perturbations.

The Spp virus is a replication-incompetent virus. The dosage administered in this study is supraphysiological. The viral burdens were detected in the heart, the liver, kidneys, spleen (around 29~35 Ct values in a real time RT-PCR assay) and lungs (26~30 Ct values) at 2 hpi. At 24 hpi, the Spp virus was hardly detectable in these tissues (33~36 Ct values or negative). Therefore, the viral particles were rapidly and effectively removed from the body. The residual viral burdens retained in the tissues at 2 hpi were in the pathological ranges from the nasal and pharyngeal swab samples in patients (overall median (interquartile range) Ct value: 28.7 (23.9–33.4)) [33]. Our data reflects parts of pathological spectra of PACS. Firstly, exposure of replication-incompetent Spp virus is transient and one time. The viral particles are rapidly cleared and undetectable 3~4 dpi in mice, which can be equivalent to the “viral-free” stage in discharged COVID-19 patients. The timelines from 6 to 24 wpi in this animal model is long enough to mimic potential cardiac sequelae in PASC in patients. Secondly, obesity is associated with and contributes to PACS [34-36]. Our data has demonstrated that cholesterol facilitates viral uptake in host cells and heart is the vulnerable organ for viral accumulation under obesity. Lastly and more importantly, the cardiac functional impairments and fibrotic development have been revealed as long-term sequelae in this model.

The first case series about persistent symptoms after acute COVID-19 was reported in July 2020 in an Italian hospitalized cohort [37], which was posited to be caused by the Wuhan original strain before the Alpha variant emerged to be the dominant variant in the world until January 2021 [38]. Different variants of SARS-CoV-2 may lead to different clinical aspects of PACS. For example, patients infected by the Wuhan variant showed higher incidences of dyspnea than those infected with Alpha or Delta variant [39]. Another report shows that Wuhan variant causes higher occurrences of dysgeusia and anosmia as compared to the Alpha variant in patients [40]. Patients infected by the Omicron variant were less likely to experience PACS than those infected by the delta variant with an odds ratio ranging from 0.24 (0.20–0.32) to 0.50 (0.43–0.59) [41]. It could be expected that PACS from individuals infected with the Wuhan variant would lead to higher burden and health costs than those infected by other variants [39]. The underlying explanations could be as follows. First, the Darwinian evolution of the mutations of Spike protein has led to less pathogenicity but higher transmissibility among VOCs [42, 43]. The Wuhan variant exhibited the highest pathogenicity which led to higher fatalities and likely PACS as well. Second, administration of antiviral medications, e.g., remdesivir, can reduce the risk of PACS [44]. Third, vaccination can reduce PACS [45]. These selective pressures reflected and contributed to the viral evolution and its adaption to a new host.

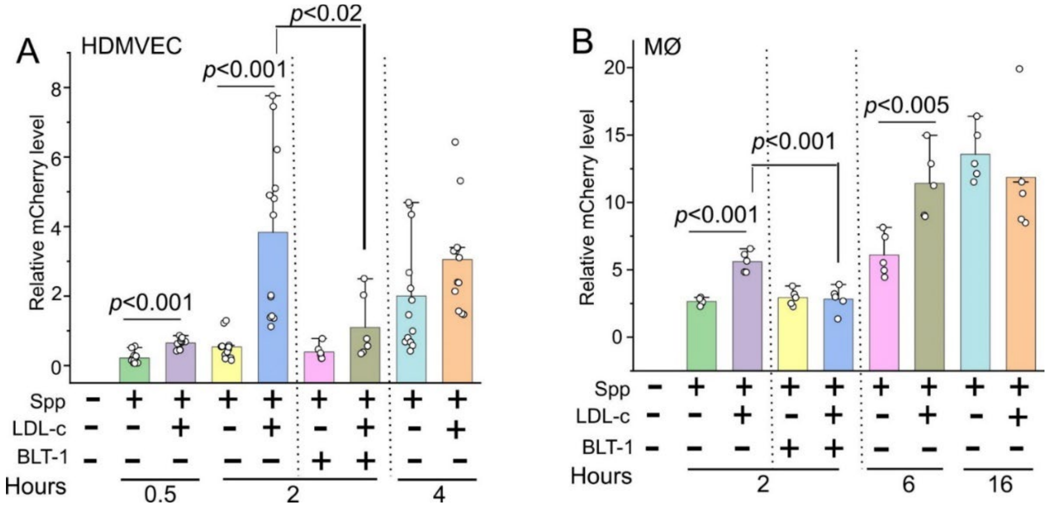

**Supplementary Fig 1** LDL-c significantly increases cellular uptake of Spp virus in HDMVEC (A) at 0.5 and 2 hpi and in MØ (B) at 2 and 6 hpi, which can be blocked with BLT-1, an inhibitor of SR-B1.

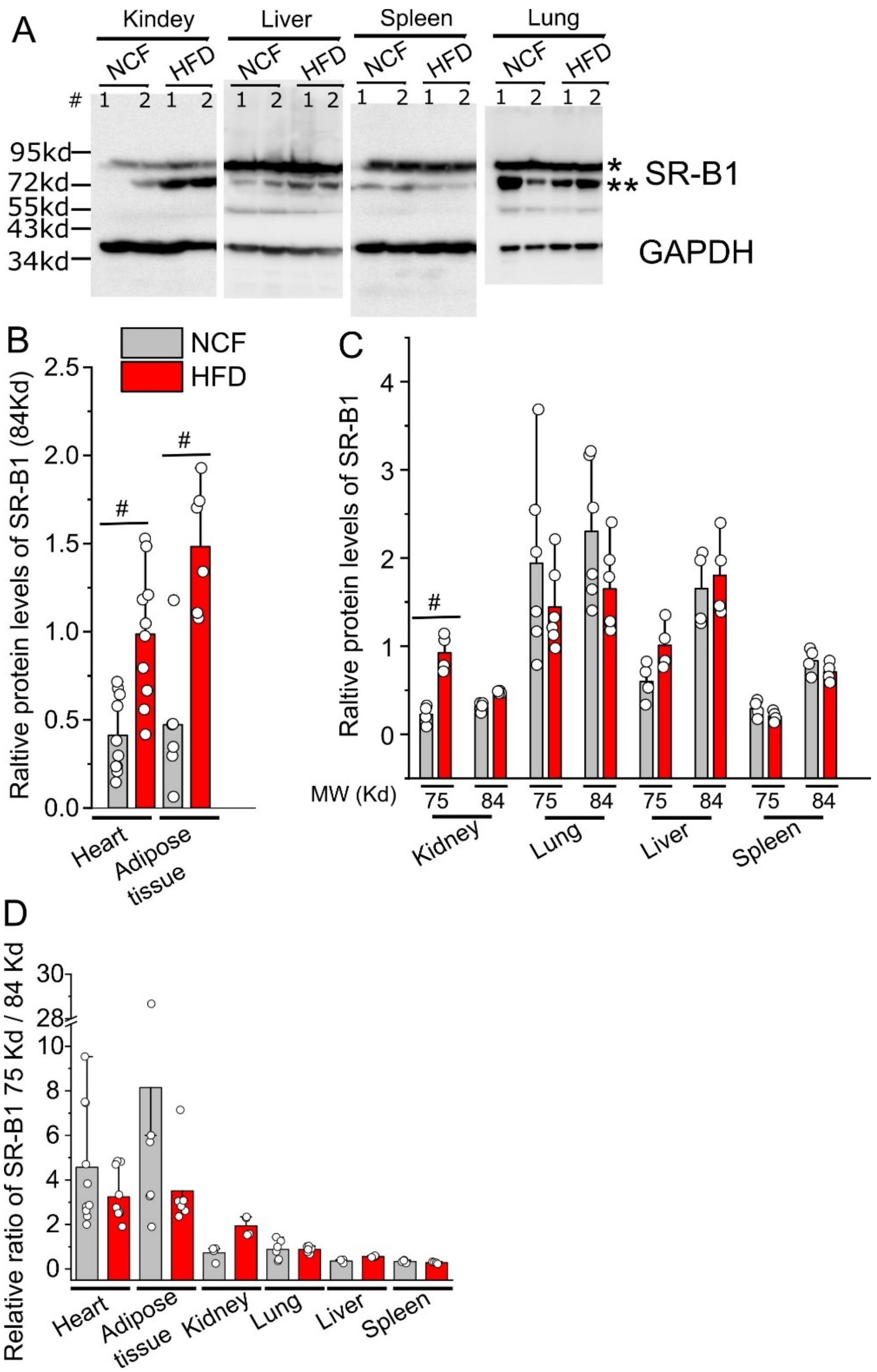

**Supplementary Fig 2** (A) The relative protein levels of SR-B1 in kidney, liver, spleen, and lungs in HFD mice and the NCF group by Western blot. Blots from two animals per group were shown. SR-B1 shows two bands in the blots, 75Kd (**) and 84Kd (*), probably due to the distinct post-translational modifications. (B) Relative protein levels of SR-B1 with the 84Kd MW band in hearts and adipose tissues. (C) Relative protein levels of SR-B1 in kidney, liver, spleen, and lungs. (D) Relative ratios of SR-B1 75Kd / 84 Kd MW bands among various tissues. # indicates $p < 0.01$ in HFD vs NCF group for SR-B1. The data from aorta is absent due to the very limited tissue collected.

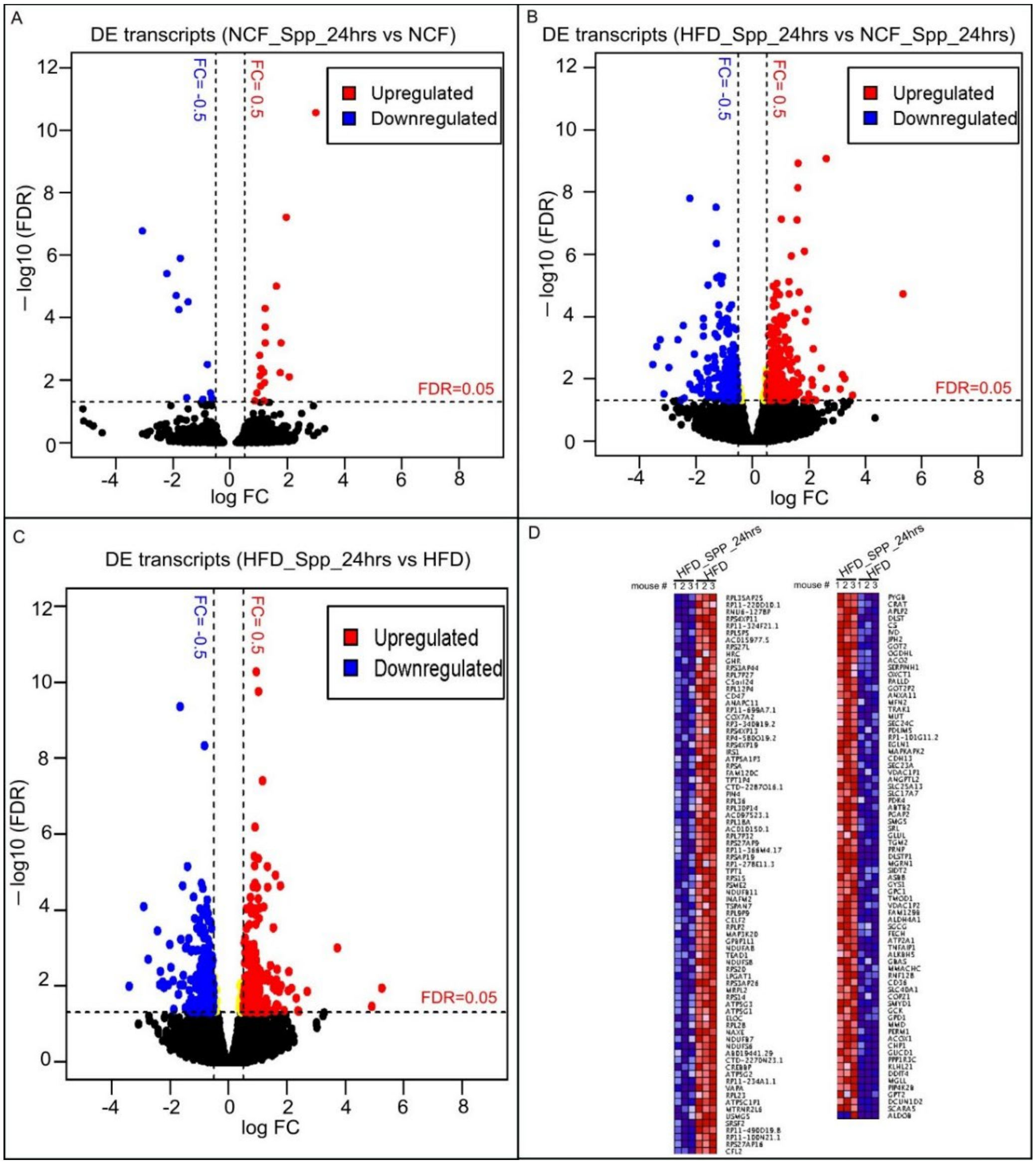

**Supplementary Fig 3** Acute changes in cardiac transcriptomes in normal and obese mice 24 hpi. (A) Volcano plot showing the cardiac DEs in NCF mice 24 hpi as compared with control NCF mice without viral administration. (B) Volcano plot showing the cardiac DEs of 24 hpi in HFD mice as compared with NCF mice. (C) Volcano plot showing the cardiac DEs in HFD mice 24 hpi as compared with HFD control mice. (D) Ranked heatmap showing the individual downregulated (left panel) and upregulated DEs (right panel), which is ranked by FDR in HFD mice 24 hpi as compared with HFD control mice. FDR < 0.05 (horizontal dashed line) is used for cut-off threshold. Right vertical dashed line: fold changes (FC) > 0.5 (increase); left vertical dashed line: FC < - 0.5 (decrease). Blue dots represent downregulated DEs and red dots represent upregulated DEs.

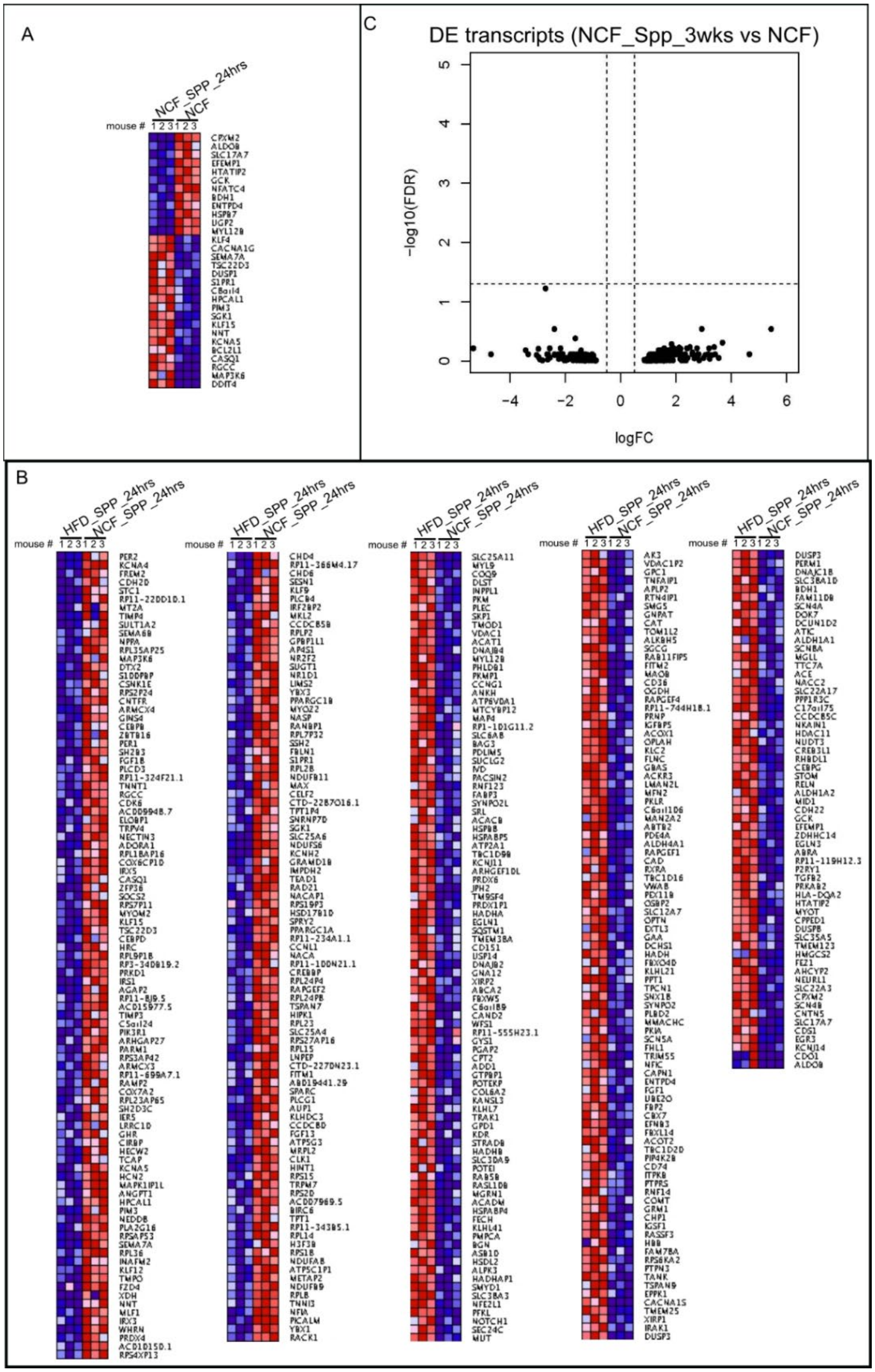

**Supplementary Fig 4** (A) Ranked heatmap showing the individual dysregulated DEs, which is ranked by FDR in NCF mice 24 hpi as compared with NCF control mice. (B) Ranked heatmap showing the individual dysregulated DEs, which is ranked by FDR in HFD mice 24 hpi as compared with NCF mice 24 hpi. (C) Volcano plot showing that there is no significant cardiac DEs in NCF mice 3 wpi as compared with control NCF mice without viral administration.

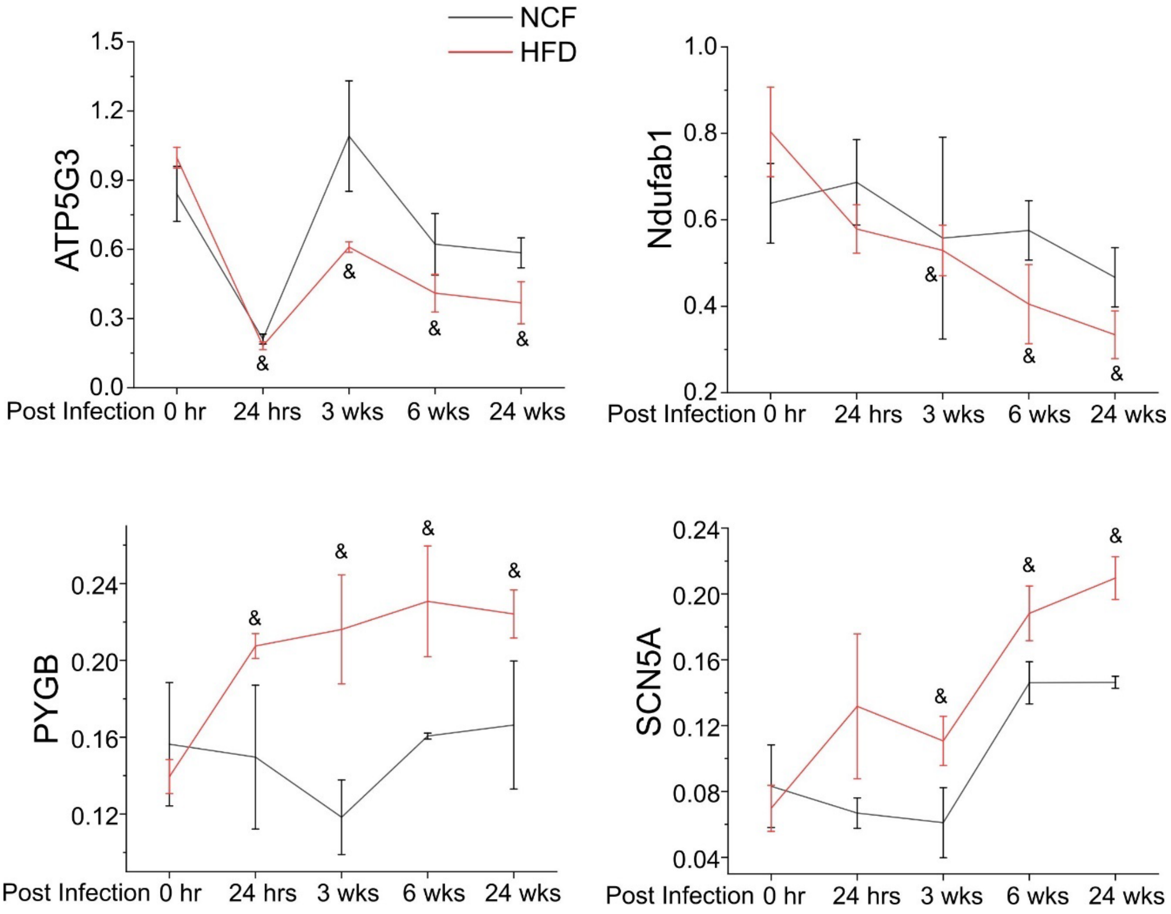

**Supplementary Fig 5** Relative changes of some representative cardiac transcripts of DEs in HDF mice during the time course of post viral administration**.** & indicates $p < 0.05$ at each post-administration time point as compared to before administration (0 hr) in HFD mice.

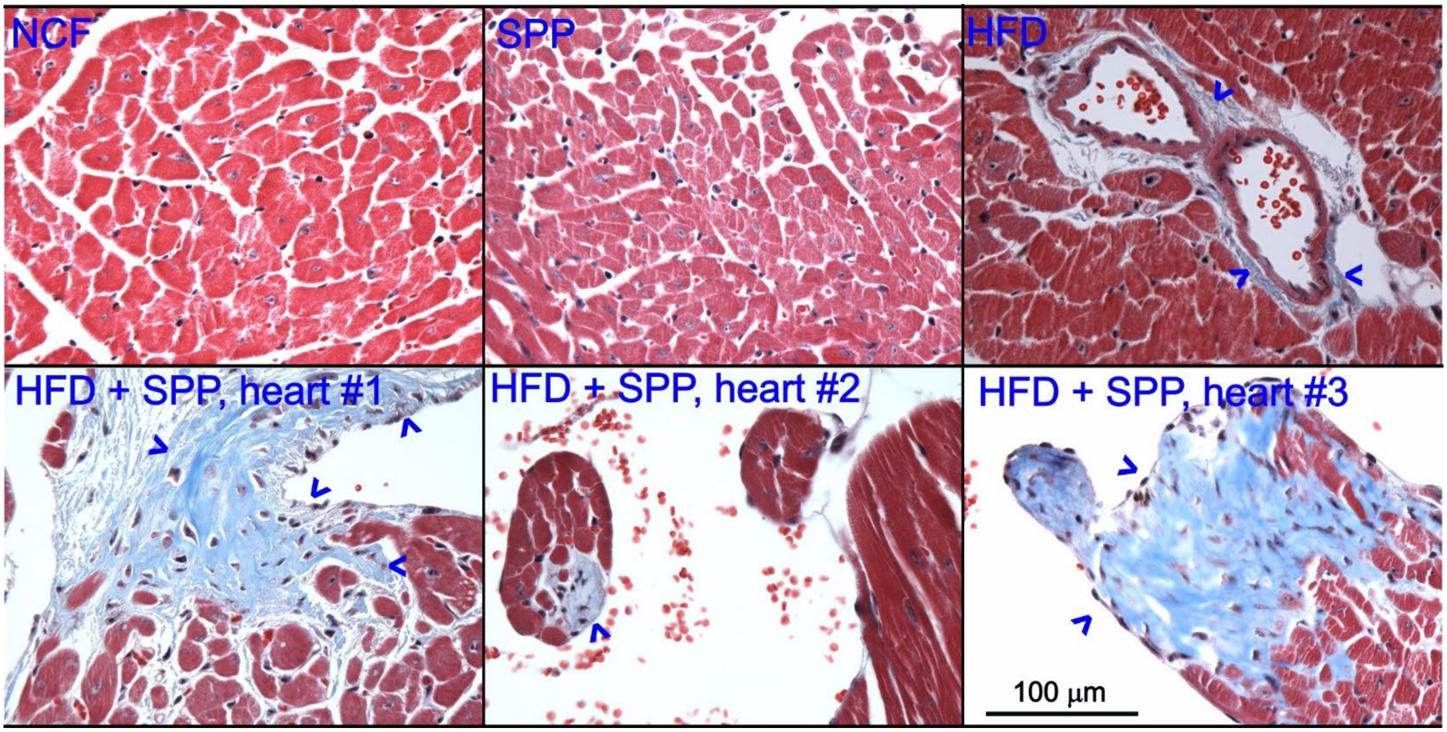

**Supplementary Fig 6** Heart cross section showing the cardiac fibrosis using Masson's trichrome staining. Focal fibrosis was evident in all three examined hearts in the HFD+SPP group but not in the control groups (6 wpi).

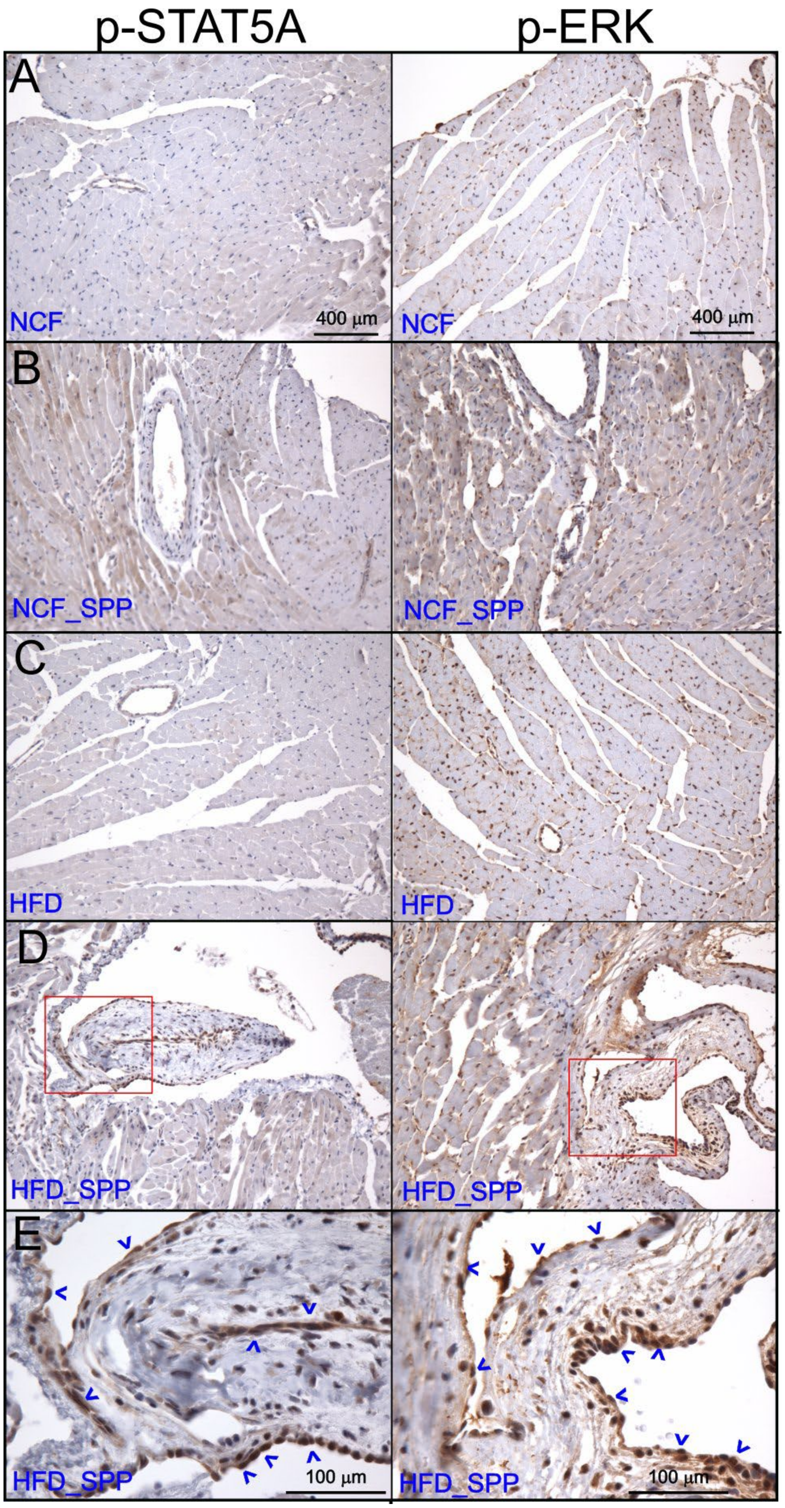

**Supplementary Fig 7** IHC staining on heart sections (6 wpi) using p-STAT5A and p-ERK antibodies. The boxed areas in (D) were shown in a higher magnification in (E). The arrow heads indicate the positive cells in the focal fibrotic regions.

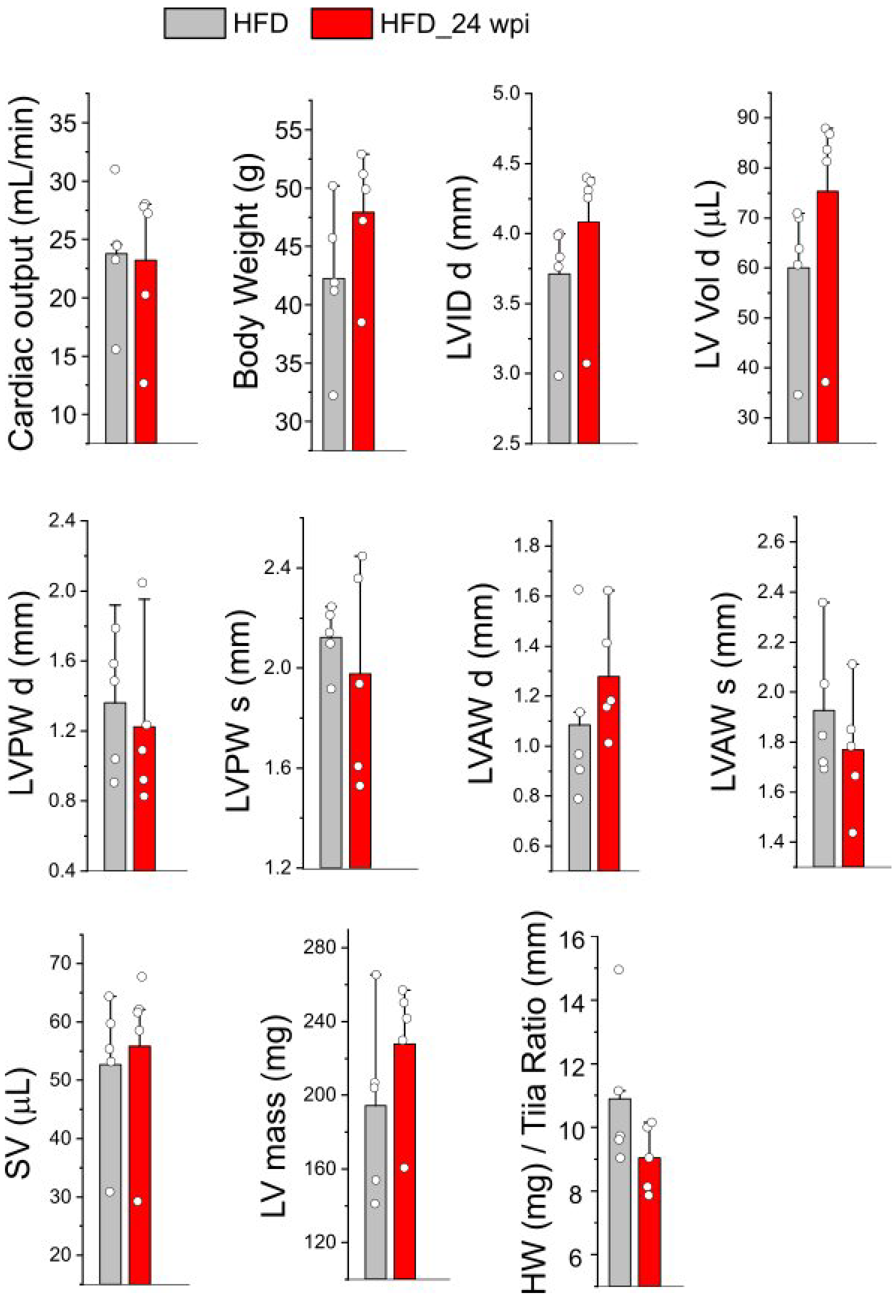

**Supplementary Fig 8** Echocardiogram showing the cardiac functions in the HFD mice 24 wpi as compared with the age-matched HFD controls (13 months old). There were no significant in cardiac output (mL/min), body weight (g) left ventricular end-diastolic diameter (LVID d), left ventricular end-diastolic volume (LV Vol d), left ventricular end-diastolic posterior wall thickness (LVPW d), left ventricular end-systolic posterior wall thickness (LVPW s), left ventricular end-diastolic anterior wall thickness (LVAW d), left ventricular end-systolic anterior wall thickness (LVAW s), and stroke volume (SV), LV mass, and heart weight-to-tibia length.

**Supplementary table 1** Source of antibodies in this study

| Antibodies | Catalog | Vendor | Dilutions for WB or IHC |
|---|---|---|---|
| Spike S1 subunit | GTX635671 | Genetex | 1:100 (IHC) |
| MRC1 | A02285-2 | Boster Bio | 1:200 (IHC) |
| SR-B1 | NB400-104 | Novus Biologicals | 1:2000 |
| CD31 | ZRB1216 | Sigma | 1:200 (IHC) |
| STAT5 | A21228 | Abclonal | 1:1000 |
| GAPDH | G8795 | Sigma | 1:2000 |
| Beta-actin | A1978 | Sigma | 1:2000 |
| Beta-tubulin | MA1-118 | Thermofisher | 1:2000 |
| PYGB | A13539 | Abclonal | 1:1000 |
| OGDH | A6391 | Abclonal | 1:1000 |
| NDUFS3 | A8013 | Abclonal | 1:1000 |
| NDUFA2 | A8136 | Abclonal | 1:1000 |
| ATP5C1 | A15257 | Abclonal | 1:1000 |
| p-STAT5A | AP0758 | Abclonal | 1:200 (IHC) |
| p-ERK | sc-7383 | Santa Cruz Biotech | 1:200 (IHC) |

**Supplementary table 2** List of primers

| species | ENSEMBL | SYMBOL | ENTREZID | 5 primers | 3 primers | size |
|---|---|---|---|---|---|---|
| mouse | ENSMUSG00000038717 | Atp5l | 27425 | TAGCCAACAATGCCACGTTT | TTACAGTTAAGGAAGCTGTG | 100 |
| mouse | ENSMUSG00000018770 | Atp5g3 | 228033 | CTCTCCAGTCCTAGTCTCTG | GGAGAGAGGAAGCGGGAGA | 188 |
| mouse | ENSMUSG00000014294 | Ndufa2 | 17991 | TTCAGGCTTTGCCGCTTAGC | TGCTTTTGGCCAAGAGAAGA | 93 |
| mouse | ENSMUSG00000030869 | Ndufab1 | 70316 | CAAGACATACAGAACTCGGT | GCTCGCGCAGGTGCCTGGA | 103 |
| mouse | ENSMUSG00000038615 | Nfe2l1 | 18023 | AAGCTCTCCCCAGTCTCTCC | CATCGCCAGAAGGAGCAGG | 131 |
| mouse | ENSMUSG00000032511 | Scn5a | 20271 | ACTTTCACCTTGGTCCAGTAC | CCGAAGGGGACCTGCCTCT | 100 |
| mouse | ENSMUSG00000033059 | Pygb | 110078 | AAGGAGAGTGCGTTGTCATAG | GAGGGGTTCTCCTGCCCA | 102 |
| mouse | ENSMUSG00000020893 | Per1 | 18626 | ATTAGTCAGCCCTCAGAGACA | CGAAACAGGGAAGGTGAAGA | 98 |
| mouse | ENSMUSG00000020456 | Ogdh | 18293 | GACATTTTCACAAAACAAACCAGC | TGGGATTCTCCAACCAGGC | 146 |

#### **Supplementary table 3** Cardiac DEs for NCF_Spp_24hrs vs NCF_CTL

| Ensembl ID | Gene symbol | NCF_Spp_24hrs-1 | NCF_Spp_24hrs-2 | NCF_Spp_24hrs-3 | NCF-CTL-1 | NCF-CTL-2 | NCF-CTL-3 | logFC | logCPM | PValue | FDR |
|---|---|---|---|---|---|---|---|---|---|---|---|
| ENSG00000121898 | CPXM2 | 8.421097163 | 4.817468434 | 5.424727771 | 61.68102601 | 53.21796724 | 49.97904554 | -3.13309 | 4.983498 | 4.00E-29 | 4.55E-25 |
| ENSG00000168209 | DDIT4 | 29.00600134 | 17.34288636 | 30.57573835 | 3.304340679 | 3.130468661 | 3.010785876 | 2.992887 | 3.944127 | 4.77E-15 | 2.72E-11 |
| ENSG00000102760 | RGCC | 37.4270985 | 27.45957007 | 27.61679593 | 8.44442618 | 8.34791643 | 6.623728927 | 1.958697 | 4.347837 | 1.60E-11 | 6.09E-08 |
| ENSG00000136872 | ALDOB | 4.678387313 | 0.481746843 | 2.958942421 | 26.43472543 | 30.26119706 | 12.64530068 | -3.07241 | 3.814489 | 5.89E-11 | 1.68E-07 |
| ENSG00000106633 | GCK | 20.11706544 | 9.634936868 | 15.78102624 | 60.94672808 | 52.17447769 | 39.14021639 | -1.74582 | 5.093367 | 5.53E-10 | 1.26E-06 |
| ENSG00000104888 | SLC17A7 | 3.74270985 | 6.744455808 | 8.876827262 | 25.33327854 | 43.30481648 | 21.67765831 | -2.21594 | 4.259694 | 2.05E-09 | 3.89E-06 |
| ENSG00000130037 | KCNA5 | 27.60248514 | 34.68577273 | 28.109953 | 9.913022037 | 12.00012987 | 7.225886103 | 1.613798 | 4.388355 | 6.01E-09 | 9.79E-06 |
| ENSG00000115380 | EFEMP1 | 6.081903506 | 3.372227904 | 4.931570701 | 19.82604407 | 16.69583286 | 16.86040091 | -1.88824 | 3.622241 | 1.37E-08 | 1.96E-05 |
| ENSG00000161267 | BDH1 | 72.98284208 | 53.95564646 | 32.05520956 | 167.7870767 | 121.0447882 | 152.9479225 | -1.47294 | 6.660341 | 2.47E-08 | 3.12E-05 |
| ENSG00000118515 | SGK1 | 112.2812955 | 83.82395075 | 77.42566001 | 34.14485368 | 42.7830717 | 40.94668791 | 1.221889 | 6.049223 | 4.43E-08 | 5.05E-05 |
| ENSG00000109854 | HTATIP2 | 4.210548581 | 7.707949495 | 8.383670192 | 24.59898061 | 28.17421795 | 17.46255808 | -1.7966 | 4.01015 | 5.34E-08 | 5.53E-05 |
| ENSG00000163884 | KLF15 | 54.73713156 | 47.6929375 | 60.16516255 | 18.35744822 | 24.52200451 | 27.69923006 | 1.22443 | 5.312901 | 2.12E-07 | 0.000201 |
| ENSG00000143318 | CASQ1 | 29.00600134 | 24.56908901 | 14.7947121 | 9.545873073 | 4.695702992 | 4.817257402 | 1.77733 | 3.972161 | 7.83E-07 | 0.000644 |
| ENSG00000112992 | NNT | 404.6805025 | 396.4776521 | 625.3231649 | 256.6371261 | 209.2196555 | 142.7112505 | 1.225113 | 8.410991 | 7.92E-07 | 0.000644 |
| ENSG00000138623 | SEMA7A | 65.49742238 | 68.88979861 | 54.74043478 | 32.67625783 | 26.08723884 | 33.72080181 | 1.0298 | 5.583611 | 2.12E-06 | 0.001608 |
| ENSG00000173641 | HSPB7 | 431.8151489 | 346.8577273 | 398.4709126 | 678.4912861 | 742.9645622 | 624.4369907 | -0.79747 | 9.07175 | 4.44E-06 | 0.003159 |
| ENSG00000170989 | S1PR1 | 62.22255126 | 47.6929375 | 61.64463376 | 34.14485368 | 20.3480463 | 25.29060136 | 1.083737 | 5.426704 | 6.44E-06 | 0.004316 |
| ENSG00000176907 | C8orf4 | 43.50900201 | 36.6127601 | 34.52099491 | 23.13038475 | 13.04361942 | 13.24745785 | 1.173706 | 4.833394 | 8.86E-06 | 0.00561 |
| ENSG00000171552 | BCL2L1 | 14.9708394 | 13.97065846 | 30.08258128 | 6.975830323 | 4.173958215 | 6.021571752 | 1.748689 | 3.773847 | 9.69E-06 | 0.005811 |
| ENSG00000157514 | TSC22D3 | 258.2469797 | 137.2978504 | 182.4681159 | 86.64715559 | 88.17486729 | 107.1839772 | 1.038157 | 7.173171 | 1.28E-05 | 0.007267 |
| ENSG00000142733 | MAP3K6 | 14.50300067 | 8.189696338 | 29.09626714 | 4.772936537 | 3.652213438 | 3.612943051 | 2.065179 | 3.537283 | 1.45E-05 | 0.007881 |
| ENSG00000198355 | PIM3 | 56.60848648 | 36.6127601 | 62.6309479 | 27.90332129 | 25.04374929 | 13.84961503 | 1.197181 | 5.257605 | 2.31E-05 | 0.01198 |
| ENSG00000120129 | DUSP1 | 94.03558498 | 58.29136805 | 95.17931453 | 40.38638608 | 49.04400902 | 28.30138724 | 1.064609 | 5.953111 | 3.14E-05 | 0.015572 |
| ENSG00000169764 | UGP2 | 391.5810181 | 336.2592967 | 343.2373208 | 621.9503456 | 518.6143082 | 585.8989315 | -0.689 | 8.868002 | 5.63E-05 | 0.025688 |
| ENSG00000006283 | CACNA1G | 38.36277596 | 46.72944381 | 51.28833529 | 22.02893786 | 27.1307284 | 22.27981548 | 0.933805 | 5.153478 | 5.64E-05 | 0.025688 |
| ENSG00000118680 | MYL12B | 282.1067549 | 290.0115997 | 290.4695143 | 489.7767184 | 426.7872275 | 435.3596377 | -0.6491 | 8.531725 | 8.53E-05 | 0.036289 |
| ENSG00000100968 | NFATC4 | 3.274871119 | 4.335721591 | 2.958942421 | 8.811575144 | 11.47838509 | 10.83882915 | -1.52216 | 2.969313 | 8.60E-05 | 0.036289 |
| ENSG00000197217 | ENTPD4 | 32.28087246 | 22.64210164 | 36.00046612 | 70.85975012 | 57.39192545 | 46.96825967 | -0.95579 | 5.508345 | 0.000101 | 0.040953 |
| ENSG00000136826 | KLF4 | 46.78387313 | 53.47389962 | 60.16516255 | 26.06757647 | 34.43515527 | 28.30138724 | 0.858013 | 5.408414 | 0.000117 | 0.045944 |
| ENSG00000115756 | HPCAL1 | 18.71354925 | 20.23336742 | 27.61679593 | 13.21736272 | 8.34791643 | 6.623728927 | 1.182163 | 4.077938 | 0.000121 | 0.04603 |

### Supplementary table 4 Cardiac DEs for HFD_Spp_24hr vs NCF_Spp_24hrs

| Ensembl ID | Gene symbol | HFD_Spp_24hr_1 | HFD_Spp_24hr_2 | HFD_Spp_24hr_3 | NCF_Spp_24hr-1 | NCF_Spp_24hr-2 | NCF_Spp_24hr-3 | logFC | logCPM | PValue | FDR |
|---|---|---|---|---|---|---|---|---|---|---|---|
| ENSG00000104888 | SLC17A7 | 47.55113902 | 43.83370118 | 54.43800474 | 3.881250009 | 6.968587561 | 9.182534619 | 2.858438 | 4.82051 | 1.84E-27 | 2.08E-23 |
| ENSG00000121898 | CPXM2 | 39.62594918 | 33.0175931 | 37.64330115 | 8.73281252 | 4.977562544 | 5.611548934 | 2.499835 | 4.48443 | 5.47E-21 | 3.10E-17 |
| ENSG00000177098 | SCN4B | 21.44463132 | 16.50879655 | 20.84859756 | 2.910937507 | 1.493268763 | 5.101408121 | 2.61654 | 3.61399 | 2.22E-13 | 8.39E-10 |
| ENSG00000174429 | ABRA | 48.01732665 | 39.84881926 | 41.69719512 | 13.09921878 | 12.44390636 | 16.8346468 | 1.615299 | 4.88266 | 4.17E-13 | 1.18E-09 |
| ENSG00000129521 | EGLN3 | 73.65764671 | 80.83617621 | 90.34392276 | 27.16875006 | 36.83396282 | 15.81436518 | 1.610538 | 5.77383 | 3.21E-12 | 7.27E-09 |
| ENSG00000175206 | NPPA | 38.69357391 | 8.539032698 | 19.11121443 | 116.4375003 | 105.0265697 | 91.82534619 | -2.215897 | 6.02138 | 8.45E-12 | 1.60E-08 |
| ENSG00000036448 | MYOM2 | 125.8706621 | 105.3147366 | 94.39781673 | 216.379688 | 240.9140271 | 339.7537809 | -1.287827 | 7.55866 | 1.90E-11 | 3.08E-08 |
| ENSG00000108947 | EFNB3 | 144.5181676 | 117.2693824 | 134.3576287 | 63.55546889 | 63.21504431 | 67.84872802 | 1.027214 | 6.63369 | 5.21E-11 | 7.37E-08 |
| ENSG00000106633 | GCK | 51.28064012 | 40.4180881 | 50.38411077 | 20.8617188 | 9.955125088 | 16.32450599 | 1.588108 | 5.01629 | 6.25E-11 | 7.87E-08 |
| ENSG00000163884 | KLF15 | 27.03888297 | 18.78587194 | 23.1651084 | 56.76328138 | 49.27786918 | 62.23717908 | -1.273287 | 5.3528 | 3.97E-10 | 4.50E-07 |
| ENSG00000109854 | HTATIP2 | 20.04606841 | 26.75563579 | 29.53551321 | 4.36640626 | 7.96410007 | 8.672393807 | 1.847612 | 4.08043 | 7.76E-10 | 7.99E-07 |
| ENSG00000119938 | PPP1R3C | 32.63313462 | 41.5566258 | 36.48504573 | 13.09921878 | 12.94166261 | 16.32450599 | 1.377596 | 4.71394 | 1.20E-09 | 1.13E-06 |
| ENSG00000225024 | AC015977.5 | 34.49788517 | 20.49367848 | 33.58940718 | 64.04062514 | 65.20606932 | 70.90957289 | -1.166067 | 5.62772 | 5.77E-09 | 5.03E-06 |
| ENSG00000236929 | RP11-699A7.1 | 89.97421402 | 55.21907811 | 75.2866023 | 161.0718754 | 162.2685389 | 136.7177377 | -1.054912 | 6.84383 | 6.50E-09 | 5.26E-06 |
| ENSG00000157514 | TSC22D3 | 74.12383435 | 78.55910082 | 96.13519986 | 267.8062506 | 141.8605325 | 188.7521005 | -1.267842 | 7.15575 | 7.47E-09 | 5.64E-06 |
| ENSG00000161267 | BDH1 | 131.9311014 | 128.6547593 | 144.2027998 | 75.68437517 | 55.74870049 | 33.15915279 | 1.29478 | 6.57891 | 1.06E-08 | 7.53E-06 |
| ENSG00000196821 | C6orf106 | 110.9526577 | 104.1761989 | 115.825542 | 58.21875013 | 60.72626303 | 62.23717908 | 0.868895 | 6.42885 | 1.31E-08 | 8.59E-06 |
| ENSG00000145675 | PIK3R1 | 86.24471292 | 78.55910082 | 99.03083841 | 225.1125005 | 138.3762387 | 199.9751984 | -1.094936 | 7.1213 | 1.36E-08 | 8.59E-06 |
| ENSG00000102760 | RGCC | 14.9180044 | 6.831226158 | 9.84517107 | 38.81250009 | 28.3721065 | 28.56788548 | -1.568674 | 4.49646 | 1.63E-08 | 9.72E-06 |
| ENSG00000082641 | NFE2L1 | 789.7218578 | 794.6993098 | 682.2124424 | 472.5421886 | 410.1511536 | 475.9613777 | 0.738593 | 9.2409 | 1.87E-08 | 1.06E-05 |
| ENSG00000116688 | MFN2 | 482.9703924 | 649.5357539 | 589.5520088 | 297.4007819 | 286.2098463 | 359.6492726 | 0.867669 | 8.7969 | 2.97E-08 | 1.60E-05 |
| ENSG00000169860 | P2RY1 | 17.71513022 | 18.21660309 | 16.21557588 | 5.336718762 | 4.479806289 | 6.631830558 | 1.657336 | 3.62302 | 3.18E-08 | 1.64E-05 |
| ENSG00000007314 | SCN4A | 46.1525761 | 72.29714351 | 72.97009146 | 28.62421881 | 18.41698141 | 30.09830792 | 1.302704 | 5.5011 | 3.95E-08 | 1.88E-05 |
| ENSG00000136872 | ALDOB | 17.24894258 | 22.20148501 | 308.6750694 | 4.851562511 | 0.497756254 | 3.060844873 | 5.336169 | 5.87172 | 4.07E-08 | 1.88E-05 |
| ENSG00000144476 | ACKR3 | 120.7425981 | 117.2693824 | 109.4551372 | 57.24843763 | 70.68138812 | 64.27774233 | 0.85542 | 6.50435 | 4.15E-08 | 1.88E-05 |
| ENSG00000172403 | SYNPO2 | 76.45477254 | 79.69763851 | 77.60311314 | 45.11953135 | 42.30928162 | 32.13887117 | 0.963494 | 5.8995 | 4.54E-08 | 1.98E-05 |
| ENSG00000084234 | APLP2 | 421.4336242 | 459.3999592 | 444.7700813 | 245.4890631 | 237.4297333 | 302.0033608 | 0.75584 | 8.46134 | 6.71E-08 | 2.82E-05 |
| ENSG00000128591 | FLNC | 629.3533105 | 773.6363624 | 841.4725626 | 479.3343761 | 407.1646161 | 355.5681461 | 0.852666 | 9.18403 | 1.07E-07 | 4.23E-05 |
| ENSG00000172399 | MYOZ2 | 619.0971825 | 553.8985877 | 542.0635366 | 870.8554707 | 890.9836953 | 1092.72162 | -0.734285 | 9.57537 | 1.08E-07 | 4.23E-05 |
| ENSG00000169047 | IRS1 | 20.51225605 | 20.49367848 | 23.74423611 | 38.32734384 | 58.73523802 | 49.99379959 | -1.183188 | 5.19018 | 1.13E-07 | 4.26E-05 |
| ENSG00000115593 | SMYD1 | 358.0321055 | 370.5940191 | 320.8367513 | 199.8843755 | 194.1249392 | 235.6850552 | 0.737418 | 8.13272 | 1.28E-07 | 4.66E-05 |
| ENSG00000119138 | KLF9 | 119.3440352 | 114.992307 | 122.1959468 | 230.4492193 | 173.2191765 | 223.9518165 | -0.815543 | 7.36899 | 1.57E-07 | 5.57E-05 |
| ENSG00000184545 | DUSP8 | 14.9180044 | 12.52391462 | 17.95295901 | 4.36640626 | 5.475318798 | 1.530422436 | 1.9701 | 3.37035 | 1.72E-07 | 5.89E-05 |
| ENSG00000148175 | STOM | 25.17413242 | 19.92440963 | 25.48161924 | 10.18828127 | 4.977562544 | 9.692675431 | 1.500833 | 4.07012 | 2.30E-07 | 7.68E-05 |
| ENSG00000219023 | RP3-340B19.2 | 14.45181676 | 10.81610808 | 12.74080962 | 31.05000007 | 29.36761901 | 27.54760386 | -1.196647 | 4.47463 | 2.62E-07 | 8.49E-05 |
| ENSG00000175931 | UBE2O | 45.22020083 | 54.64980927 | 45.17196138 | 24.25781255 | 21.90127519 | 25.50704061 | 1.012872 | 5.20602 | 3.04E-07 | 9.56E-05 |
| ENSG00000081248 | CACNA1S | 36.36263572 | 27.89417348 | 43.43457825 | 16.01015629 | 14.93268763 | 15.30422436 | 1.215458 | 4.72755 | 3.59E-07 | 0.00011 |
| ENSG00000214263 | RPSAP53 | 33.09932226 | 29.03271117 | 31.27289634 | 63.55546889 | 55.74870049 | 57.13577096 | -0.913084 | 5.53073 | 3.99E-07 | 0.00012 |
| ENSG00000179094 | PER1 | 32.63313462 | 15.37025886 | 33.58940718 | 144.5765628 | 53.75767547 | 73.46027695 | -1.730613 | 5.91743 | 4.00E-07 | 0.00012 |
| ENSG00000153179 | RASSF3 | 39.15976154 | 50.0956585 | 41.69719512 | 15.03984378 | 20.40800643 | 24.48675898 | 1.124532 | 5.02507 | 4.27E-07 | 0.00012 |
| ENSG00000178053 | MLF1 | 58.73964232 | 68.31226158 | 50.38411077 | 119.833594 | 111.497401 | 94.37605025 | -0.87601 | 6.4118 | 4.39E-07 | 0.00012 |
| ENSG00000147255 | IGSF1 | 49.41588957 | 62.61957312 | 39.95981199 | 26.68359381 | 19.91025018 | 22.95633655 | 1.124522 | 5.23808 | 4.55E-07 | 0.00012 |
| ENSG00000105953 | OGDH | 1177.589972 | 1755.055854 | 1504.573791 | 847.0828144 | 799.3965445 | 911.6216313 | 0.794175 | 10.1878 | 4.90E-07 | 0.00013 |
| ENSG00000138796 | HADH | 169.6923 | 152.5640509 | 144.2027998 | 62.10000014 | 77.15221943 | 104.5788665 | 0.938145 | 6.89734 | 5.10E-07 | 0.00013 |
| ENSG00000169116 | PARM1 | 24.70794478 | 21.06294732 | 19.11121443 | 38.32734384 | 52.26440671 | 46.93295472 | -1.071636 | 5.12896 | 5.28E-07 | 0.00013 |
| ENSG00000120729 | MYOT | 10.72231566 | 12.52391462 | 16.21557588 | 4.36640626 | 2.986537526 | 3.060844873 | 1.885952 | 3.19379 | 5.90E-07 | 0.00014 |
| ENSG00000138623 | SEMA7A | 37.29501099 | 31.30978656 | 35.32679031 | 67.92187515 | 71.17914438 | 56.62563015 | -0.908777 | 5.67856 | 5.91E-07 | 0.00014 |
| ENSG00000022267 | FHL1 | 95.56846567 | 122.392802 | 111.1925203 | 69.37734391 | 39.82050035 | 56.11548934 | 0.989528 | 6.37644 | 5.99E-07 | 0.00014 |
| ENSG00000129559 | NEDD8 | 48.94970193 | 33.0175931 | 36.48504573 | 81.50625018 | 73.66792565 | 70.90957289 | -0.920875 | 5.8773 | 6.85E-07 | 0.00016 |
| ENSG00000187446 | CHP1 | 33.56550989 | 40.98735695 | 36.48504573 | 14.55468753 | 15.92820014 | 20.9157733 | 1.106928 | 4.79971 | 8.12E-07 | 0.00018 |
| ENSG00000159423 | ALDH4A1 | 60.60439286 | 69.45079928 | 54.43800474 | 32.50546882 | 34.34518155 | 32.13887117 | 0.896671 | 5.58738 | 8.40E-07 | 0.00019 |
| ENSG00000218682 | AC010150.1 | 125.4044745 | 77.42056313 | 99.60996612 | 193.0921879 | 174.7124453 | 184.1608332 | -0.862409 | 7.16812 | 8.53E-07 | 0.00019 |
| ENSG00000181904 | C5orf24 | 20.51225605 | 18.21660309 | 11.5825542 | 38.32734384 | 41.31376911 | 33.15915279 | -1.147284 | 4.8319 | 8.87E-07 | 0.00019 |
| ENSG00000157150 | TIMP4 | 5.128064012 | 9.108301545 | 12.16168191 | 64.5257814 | 12.44390636 | 66.31830558 | -2.445256 | 4.88701 | 9.32E-07 | 0.0002 |
| ENSG00000100234 | TIMP3 | 55.01014122 | 42.12589464 | 60.80840955 | 156.7054691 | 79.6410007 | 114.7816827 | -1.151425 | 6.43 | 9.80E-07 | 0.0002 |
| ENSG00000109906 | ZBTB16 | 32.16694698 | 47.24931426 | 53.27974932 | 184.8445317 | 57.73972551 | 197.4244943 | -1.731563 | 6.597 | 1.03E-06 | 0.00021 |
| ENSG00000149596 | JPH2 | 379.4767369 | 336.4378883 | 358.4800525 | 230.9343755 | 238.4252458 | 232.1140695 | 0.615683 | 8.21366 | 1.11E-06 | 0.00022 |
| ENSG00000146729 | GBAS | 348.2421652 | 354.6544914 | 301.7255369 | 144.5765628 | 179.6900078 | 232.1140695 | 0.853311 | 8.02758 | 1.11E-06 | 0.00022 |
| ENSG00000145494 | NDUFS6 | 120.2764105 | 108.7303497 | 113.5090312 | 182.9039067 | 184.6675704 | 176.508721 | -0.665784 | 7.21946 | 1.18E-06 | 0.00023 |
| ENSG00000080546 | SESN1 | 68.5295827 | 57.4961535 | 59.65015413 | 103.8234377 | 97.56022586 | 129.5757663 | -0.828637 | 6.44928 | 1.25E-06 | 0.00024 |
| ENSG00000112992 | NNT | 276.449269 | 271.5412398 | 254.8161924 | 419.6601572 | 409.6533974 | 646.8585498 | -0.878044 | 8.57406 | 1.38E-06 | 0.00026 |
| ENSG00000151729 | SLC25A4 | 9800.662702 | 8418.916971 | 8604.100387 | 13661.02972 | 12807.76618 | 13492.71434 | -0.575047 | 13.4424 | 1.41E-06 | 0.00026 |
| ENSG00000141959 | PFKL | 127.7354127 | 111.5766939 | 112.9299034 | 69.86250016 | 75.16119441 | 66.31830558 | 0.738763 | 6.56747 | 1.47E-06 | 0.00026 |
| ENSG00000117054 | ACADM | 190.2045561 | 181.596762 | 169.1052913 | 101.8828127 | 103.0355447 | 125.4946398 | 0.712608 | 7.19061 | 1.69E-06 | 0.0003 |
| ENSG00000130255 | RPL36 | 135.1944149 | 79.69763851 | 137.832395 | 213.4687505 | 228.4701208 | 220.3808308 | -0.90646 | 7.414 | 1.76E-06 | 0.00031 |
| ENSG00000249617 | RP11-366M4.1 | 123.5397239 | 78.55910082 | 88.02741192 | 170.2898441 | 172.223664 | 177.0188618 | -0.833676 | 7.09132 | 1.79E-06 | 0.00031 |
| ENSG00000102763 | VWA8 | 97.89940386 | 85.39032698 | 76.44485772 | 38.81250009 | 41.81152537 | 58.15605258 | 0.908692 | 6.07303 | 1.97E-06 | 0.00033 |
| ENSG00000240371 | RPS4XP13 | 98.83177913 | 64.89664851 | 96.13519986 | 163.4976566 | 137.8784825 | 166.8160456 | -0.84572 | 6.93875 | 2.14E-06 | 0.00036 |
| ENSG00000120833 | SOCS2 | 14.9180044 | 18.78587194 | 13.89906504 | 53.85234387 | 38.32723159 | 27.03746304 | -1.325584 | 4.8623 | 2.18E-06 | 0.00036 |
| ENSG00000121769 | FABP3 | 1977.567958 | 1908.758442 | 1861.31646 | 1358.922659 | 1157.781048 | 1345.751462 | 0.573509 | 10.6461 | 2.44E-06 | 0.00039 |
| ENSG00000172216 | CEBPB | 10.25612802 | 8.539032698 | 17.3738313 | 64.04062514 | 23.89230021 | 32.13887117 | -1.734266 | 4.77378 | 2.65E-06 | 0.00042 |
| ENSG00000146085 | MUT | 118.4116599 | 138.3323297 | 113.5090312 | 65.98125015 | 79.6410007 | 74.99069939 | 0.7462 | 6.63321 | 2.77E-06 | 0.00044 |
| ENSG00000130528 | HRC | 36.82882336 | 17.0780654 | 19.11121443 | 62.10000014 | 67.6948506 | 44.89239147 | -1.238113 | 5.418 | 2.93E-06 | 0.00045 |
| ENSG00000136560 | TANK | 25.17413242 | 23.90929155 | 24.32336382 | 13.09921878 | 10.45288134 | 9.182534619 | 1.158818 | 4.217 | 2.96E-06 | 0.00045 |
| ENSG00000198952 | SMG5 | 82.98139946 | 91.08301545 | 81.0778794 | 55.30781263 | 47.28684417 | 47.44309553 | 0.763628 | 6.09516 | 3.35E-06 | 0.00051 |
| ENSG00000150893 | FREM2 | 0 | 2.277075386 | 0 | 9.703125022 | 7.96410007 | 4.591267309 | -3.265898 | 2.38288 | 3.73E-06 | 0.00055 |
| ENSG00000108861 | DUSP3 | 19.57988077 | 24.4785604 | 19.69034214 | 8.73281252 | 9.955125088 | 7.652112182 | 1.263322 | 3.98841 | 3.76E-06 | 0.00055 |
| ENSG00000169253 | RP11-220D10.: | 1.398562912 | 0.569268847 | 1.15825542 | 8.247656269 | 4.479806289 | 8.162522994 | -2.638975 | 2.35893 | 3.82E-06 | 0.00056 |
| ENSG00000171298 | GAA | 55.47632885 | 48.38785196 | 64.28317581 | 34.93125008 | 23.89230021 | 29.07802629 | 0.932178 | 5.4431 | 4.13E-06 | 0.00059 |

| | | | | | | | | | | | |
|---|---|---|---|---|---|---|---|---|---|---|---|
| ENSG00000070159 | PTPN3 | 49.8820772 | 64.32737966 | 44.59283367 | 31.53515632 | 14.43493138 | 24.9968998 | 1.157411 | 5.28836 | 4.47E-06 | 0.00063 |
| ENSG00000198663 | C6orf89 | 212.115375 | 177.6118801 | 189.3747612 | 124.6851565 | 121.9502823 | 124.984499 | 0.641664 | 7.31635 | 5.07E-06 | 0.00071 |
| ENSG00000213856 | VDAC1P2 | 108.6217195 | 118.9771889 | 105.4012432 | 61.61484389 | 58.23748176 | 78.05154426 | 0.75081 | 6.48031 | 5.58E-06 | 0.00077 |
| ENSG00000188338 | SLC38A3 | 90.44040166 | 96.20643506 | 97.29345528 | 63.07031264 | 48.78011293 | 58.15605258 | 0.737432 | 6.25672 | 5.86E-06 | 0.0008 |
| ENSG00000185641 | CTD-2287O16. | 203.2578099 | 147.4406313 | 183.5834841 | 279.9351569 | 297.6582401 | 278.0267426 | -0.676853 | 7.86408 | 5.94E-06 | 0.0008 |
| ENSG00000055118 | KCNH2 | 123.5397239 | 124.1006085 | 118.1420528 | 217.8351567 | 170.7303953 | 191.8129454 | -0.665613 | 7.31242 | 6.28E-06 | 0.00084 |
| ENSG00000182255 | KCNA4 | 0 | 0.569268847 | 0.57912771 | 3.881250009 | 4.479806289 | 4.591267309 | -3.381903 | 1.77654 | 7.00E-06 | 0.00092 |
| ENSG00000113504 | SLC12A7 | 62.00295578 | 93.92935968 | 75.2866023 | 46.57500011 | 43.30479413 | 31.62873035 | 0.920889 | 5.89396 | 7.90E-06 | 0.00103 |
| ENSG00000011105 | TSPAN9 | 27.50507061 | 47.81858311 | 45.17196138 | 15.03984378 | 21.90127519 | 16.32450599 | 1.165352 | 4.88655 | 8.59E-06 | 0.00108 |
| ENSG00000109819 | PPARGC1A | 142.6534171 | 144.594287 | 133.778501 | 203.7656255 | 208.0621143 | 229.5633655 | -0.606266 | 7.47804 | 8.59E-06 | 0.00108 |
| ENSG00000149557 | FEZ1 | 8.857565111 | 7.969763851 | 5.7912771 | 0.970312502 | 1.991025018 | 2.040563249 | 2.155564 | 2.48108 | 8.71E-06 | 0.00108 |
| ENSG00000112695 | COX7A2 | 17.24894258 | 15.37025886 | 20.84859756 | 32.99062507 | 44.79806289 | 32.64901198 | -1.046639 | 4.83312 | 8.71E-06 | 0.00108 |
| ENSG00000074416 | MGLL | 14.9180044 | 17.0780654 | 16.21557588 | 7.277343766 | 4.479806289 | 7.14197137 | 1.336838 | 3.59294 | 9.33E-06 | 0.00115 |
| ENSG00000159592 | GPBP1L1 | 55.94251649 | 48.9571208 | 41.69719512 | 93.15000021 | 85.61407575 | 73.97041776 | -0.779244 | 6.08474 | 1.02E-05 | 0.00125 |
| ENSG00000197217 | ENTPD4 | 56.40870413 | 81.40544505 | 51.54236619 | 33.47578133 | 23.39454396 | 37.24027929 | 1.005588 | 5.5845 | 1.08E-05 | 0.0013 |
| ENSG00000177600 | RPLP2 | 559.8913525 | 335.8686195 | 517.161045 | 844.6570332 | 812.3382071 | 774.3937528 | -0.782216 | 9.32675 | 1.09E-05 | 0.0013 |
| ENSG00000014824 | SLC30A9 | 103.0274679 | 116.1308447 | 96.13519986 | 58.70390638 | 67.19709434 | 67.84872802 | 0.702282 | 6.42063 | 1.13E-05 | 0.00133 |
| ENSG00000182606 | TRAK1 | 173.8879888 | 216.8914305 | 185.3208672 | 135.3585941 | 108.5108635 | 115.2918235 | 0.679472 | 7.29118 | 1.15E-05 | 0.00133 |
| ENSG00000108107 | RPL28 | 376.679611 | 269.8334333 | 391.490332 | 552.59297 | 536.083486 | 593.2937645 | -0.695595 | 8.82856 | 1.15E-05 | 0.00133 |
| ENSG00000187642 | PERM1 | 20.51225605 | 26.18636694 | 22.00685298 | 5.336718762 | 9.955125088 | 12.7535203 | 1.288633 | 4.08203 | 1.36E-05 | 0.00155 |
| ENSG00000183873 | SCN5A | 298.360088 | 539.6668665 | 601.7136907 | 281.3906256 | 217.0217269 | 232.6242103 | 0.976287 | 8.50012 | 1.39E-05 | 0.00156 |
| ENSG00000138029 | HADHB | 2126.748002 | 2223.564115 | 1960.347298 | 1091.116409 | 1168.233929 | 1619.697079 | 0.702184 | 10.7305 | 1.39E-05 | 0.00156 |
| ENSG00000126368 | NR1D1 | 141.2548541 | 91.65228429 | 130.3037347 | 215.8945317 | 184.6675704 | 211.1982962 | -0.748935 | 7.35619 | 1.48E-05 | 0.0016 |
| ENSG00000142733 | MAP3K6 | 4.661876374 | 4.554150772 | 3.47476626 | 15.03984378 | 8.461856324 | 30.09830792 | -2.052927 | 3.61055 | 1.49E-05 | 0.0016 |
| ENSG00000095370 | SH2D3C | 19.57988077 | 26.18636694 | 20.26946985 | 47.06015636 | 32.85191279 | 55.60534852 | -1.039336 | 5.11942 | 1.49E-05 | 0.0016 |
| ENSG00000161533 | ACOX1 | 80.18427364 | 88.80594006 | 87.44828421 | 39.78281259 | 39.82050035 | 63.25746071 | 0.843746 | 6.07238 | 1.49E-05 | 0.0016 |
| ENSG00000141298 | SSH2 | 59.20582995 | 51.23419619 | 49.22585535 | 82.47656269 | 93.57817582 | 85.19351563 | -0.704253 | 6.15812 | 1.49E-05 | 0.0016 |
| ENSG00000136383 | ALPK3 | 130.5325385 | 132.6396412 | 162.7348865 | 101.3976565 | 76.15670692 | 78.05154426 | 0.733577 | 6.83717 | 1.51E-05 | 0.0016 |
| ENSG00000163092 | XIRP2 | 490.8955822 | 432.0750545 | 513.6862788 | 332.332032 | 332.5011779 | 261.7022366 | 0.632532 | 8.62468 | 1.56E-05 | 0.00163 |
| ENSG00000248610 | HSPA8P4 | 77.38714781 | 95.06789737 | 85.13177337 | 55.30781263 | 51.76665046 | 49.48365878 | 0.71382 | 6.12522 | 1.67E-05 | 0.00173 |
| ENSG00000150401 | DCUN1D2 | 18.18131786 | 30.74051771 | 21.42772527 | 8.247656269 | 7.466343816 | 12.24337949 | 1.319552 | 4.09948 | 1.75E-05 | 0.0018 |
| ENSG00000176842 | IRX5 | 10.72231566 | 3.984881926 | 8.10778794 | 18.92109379 | 17.4214689 | 22.95633655 | -1.341438 | 3.8984 | 1.87E-05 | 0.00191 |
| ENSG00000242299 | RP11-234A1.1 | 178.5498651 | 146.3020936 | 158.1018648 | 232.8750005 | 236.9319771 | 264.2529407 | -0.601279 | 7.6732 | 1.99E-05 | 0.00201 |
| ENSG00000169100 | SLC25A6 | 74.59002199 | 70.58933697 | 72.97009146 | 131.477344 | 104.0310572 | 112.2309787 | -0.671828 | 6.5783 | 2.05E-05 | 0.00205 |
| ENSG00000105048 | TNNT1 | 5.128064012 | 2.277075386 | 5.21214939 | 15.03984378 | 11.44839385 | 11.73323868 | -1.569951 | 3.26993 | 2.18E-05 | 0.00217 |
| ENSG00000204434 | POTEKP | 97.43321622 | 91.08301545 | 106.5594986 | 60.64453139 | 66.69933809 | 57.64591177 | 0.672319 | 6.33724 | 2.20E-05 | 0.00217 |
| ENSG00000147573 | TRIM55 | 24.70794478 | 28.46344233 | 30.69376863 | 13.09921878 | 13.93717512 | 14.79408355 | 0.993818 | 4.44414 | 2.29E-05 | 0.00223 |
| ENSG00000066926 | FECH | 75.05620963 | 89.94447775 | 81.0778794 | 44.14921885 | 51.2688942 | 54.07492609 | 0.716613 | 6.05969 | 2.35E-05 | 0.00227 |
| ENSG00000166016 | ABTB2 | 45.68638847 | 46.11077657 | 46.3302168 | 17.95078129 | 27.87435025 | 29.07802629 | 0.883886 | 5.18354 | 2.36E-05 | 0.00227 |
| ENSG00000115380 | EFEMP1 | 10.72231566 | 20.49367848 | 14.47819275 | 6.307031264 | 3.484293781 | 5.101408121 | 1.589056 | 3.44545 | 2.38E-05 | 0.00227 |
| ENSG00000095397 | WHRN | 21.91081896 | 20.49367848 | 23.1651084 | 42.20859385 | 34.84293781 | 43.36196903 | -0.874972 | 5.00802 | 2.56E-05 | 0.00241 |
| ENSG00000224631 | RPS27AP16 | 301.6234014 | 245.3548729 | 265.2404912 | 414.8085947 | 421.1017912 | 373.9332153 | -0.573323 | 8.40225 | 2.57E-05 | 0.00241 |
| ENSG00000152822 | GRM1 | 26.1065077 | 24.4785604 | 38.80155657 | 16.49531254 | 11.94615011 | 12.7535203 | 1.105768 | 4.49568 | 2.72E-05 | 0.00253 |
| ENSG00000125691 | RPL23 | 479.2408913 | 380.2715895 | 446.5074644 | 690.3773453 | 657.0382558 | 601.4560175 | -0.57667 | 9.08692 | 2.75E-05 | 0.00253 |
| ENSG00000178814 | OPLAH | 39.62594918 | 40.4180881 | 45.75108909 | 19.8914063 | 24.39005646 | 25.50704061 | 0.847027 | 5.0627 | 3.00E-05 | 0.00275 |
| ENSG00000168264 | IRF2BP2 | 31.23457171 | 26.75563579 | 36.48504573 | 52.88203137 | 50.77113795 | 60.19661583 | -0.793641 | 5.46688 | 3.06E-05 | 0.00277 |
| ENSG00000198355 | PIM3 | 25.64032006 | 28.46344233 | 30.11464092 | 58.70390638 | 37.82947533 | 64.78788314 | -0.93985 | 5.39565 | 3.18E-05 | 0.00285 |
| ENSG00000105426 | PTPRS | 46.61876374 | 79.12836967 | 56.75451558 | 35.90156258 | 32.85191279 | 18.87521005 | 1.049867 | 5.51363 | 3.20E-05 | 0.00285 |
| ENSG00000108666 | C17orf75 | 13.98562912 | 13.66245232 | 11.00342649 | 3.881250009 | 5.475318798 | 5.101408121 | 1.416452 | 3.29304 | 3.29E-05 | 0.00291 |
| ENSG00000187079 | TEAD1 | 111.4188453 | 113.8537693 | 83.97351795 | 157.1906254 | 161.7707827 | 165.7957639 | -0.645481 | 7.06181 | 3.34E-05 | 0.00293 |
| ENSG00000128016 | ZFP36 | 8.857565111 | 8.539032698 | 8.68691565 | 23.7726563 | 13.93717512 | 28.05774467 | -1.325721 | 4.04261 | 3.36E-05 | 0.00293 |
| ENSG00000117592 | PRDX6 | 136.1267901 | 130.9318347 | 144.2027998 | 97.03125022 | 80.13875696 | 91.82534619 | 0.611589 | 6.83573 | 3.73E-05 | 0.00323 |
| ENSG00000113140 | SPARC | 222.8376907 | 236.2465713 | 224.7015515 | 306.1335944 | 364.3575782 | 327.0002606 | -0.544894 | 8.13647 | 3.81E-05 | 0.00327 |
| ENSG00000168334 | XIRP1 | 27.50507061 | 68.88153043 | 44.01370596 | 15.52500004 | 22.8967877 | 19.89549167 | 1.258282 | 5.0717 | 3.85E-05 | 0.00328 |
| ENSG00000197296 | FITM2 | 44.75401319 | 55.78834696 | 58.49189871 | 30.07968757 | 29.86537526 | 32.13887117 | 0.780543 | 5.4129 | 3.96E-05 | 0.00334 |
| ENSG00000132326 | PER2 | 0 | 0.569268847 | 0.57912771 | 7.762500018 | 2.488781272 | 4.081126497 | -3.525093 | 1.87959 | 4.12E-05 | 0.00346 |
| ENSG00000171867 | PRNP | 47.08495138 | 44.97223888 | 42.85545054 | 30.56484382 | 20.40800643 | 24.48675898 | 0.837134 | 5.16899 | 4.26E-05 | 0.00355 |
| ENSG00000157637 | SLC38A10 | 21.44463132 | 13.09318347 | 20.26946985 | 7.277343766 | 8.959612579 | 6.121689746 | 1.293397 | 3.78978 | 4.39E-05 | 0.00363 |
| ENSG00000276293 | PIP4K2B | 22.3770066 | 23.90929155 | 31.27289634 | 10.67343752 | 12.44390636 | 14.28394274 | 1.042723 | 4.31936 | 4.65E-05 | 0.00382 |
| ENSG00000091428 | RAPGEF4 | 38.69357391 | 42.12589464 | 37.64330115 | 21.34687505 | 24.88781272 | 20.40563249 | 0.826993 | 4.98751 | 4.76E-05 | 0.00388 |
| ENSG00000163833 | FBXO40 | 70.86052089 | 126.3776839 | 92.6604336 | 35.41640633 | 57.73972551 | 56.11548934 | 0.954218 | 6.20426 | 4.93E-05 | 0.00397 |
| ENSG00000165416 | SUGT1 | 38.69357391 | 34.72539964 | 30.69376863 | 51.42656262 | 63.21504431 | 61.72703827 | -0.751985 | 5.58424 | 4.96E-05 | 0.00397 |
| ENSG00000132763 | MMACHC | 25.17413242 | 31.30978656 | 26.06074695 | 11.64375003 | 15.43044389 | 14.79408355 | 0.973443 | 4.43149 | 4.98E-05 | 0.00397 |
| ENSG00000177707 | NECTIN3 | 7.459002199 | 3.415613079 | 3.47476626 | 11.15859378 | 15.92820014 | 13.77380193 | -1.452089 | 3.37906 | 5.19E-05 | 0.00412 |
| ENSG00000119471 | HSDL2 | 129.6001632 | 154.8411263 | 125.0915854 | 71.80312516 | 71.17914438 | 103.5585849 | 0.732146 | 6.78273 | 5.39E-05 | 0.00425 |
| ENSG00000178035 | IMPDH2 | 62.00295578 | 51.23419619 | 60.80840955 | 96.06093772 | 84.12080699 | 93.86590944 | -0.651748 | 6.24628 | 5.55E-05 | 0.00434 |
| ENSG00000159167 | STC1 | 0.932375275 | 0.569268847 | 0.57912771 | 10.18828127 | 3.484293781 | 4.081126497 | -2.958332 | 2.15178 | 5.59E-05 | 0.00434 |
| ENSG00000069535 | MAOB | 52.21301539 | 58.06542235 | 74.70747459 | 33.47578133 | 34.34518155 | 39.28084254 | 0.782445 | 5.62655 | 5.75E-05 | 0.00443 |
| ENSG00000270885 | RASL10B | 53.14539067 | 59.77322889 | 53.27974932 | 35.41640633 | 32.85191279 | 33.15915279 | 0.709155 | 5.50653 | 5.85E-05 | 0.00448 |
| ENSG00000174996 | KLC2 | 40.09213682 | 37.00247502 | 42.27632283 | 18.92109379 | 20.40800643 | 27.03746304 | 0.84788 | 4.99189 | 5.93E-05 | 0.00451 |
| ENSG00000101333 | PLCB4 | 35.89644808 | 44.97223888 | 40.5389397 | 65.98125015 | 57.73972551 | 87.74421969 | -0.803069 | 5.82306 | 6.06E-05 | 0.00453 |
| ENSG00000176485 | PLA2G16 | 17.24894258 | 14.80099001 | 23.1651084 | 34.44609383 | 31.35864403 | 38.26056091 | -0.916652 | 4.79135 | 6.09E-05 | 0.00453 |
| ENSG00000176986 | SEC24C | 78.78571072 | 112.1459628 | 96.71432757 | 48.51562511 | 57.24196925 | 65.80816477 | 0.741975 | 6.27064 | 6.10E-05 | 0.00453 |
| ENSG00000143318 | CASQ1 | 13.05325385 | 8.539032698 | 5.7912771 | 30.07968757 | 25.38556897 | 15.30422436 | -1.34104 | 4.1432 | 6.12E-05 | 0.00453 |
| ENSG00000213326 | RPS7P11 | 6.992814561 | 4.554150772 | 7.52866023 | 12.12890628 | 17.91922516 | 17.34478761 | -1.295403 | 3.61132 | 6.23E-05 | 0.00457 |
| ENSG00000111252 | SH2B3 | 3.263313462 | 2.846344233 | 2.89563855 | 10.67343752 | 8.461856324 | 9.182534619 | -1.625538 | 2.8781 | 6.26E-05 | 0.00457 |
| ENSG00000146477 | SLC22A3 | 5.128064012 | 5.123419619 | 6.94953252 | 0 | 1.991025018 | 1.020281624 | 2.442709 | 2.10507 | 6.32E-05 | 0.00457 |
| ENSG00000163349 | HIPK1 | 107.2231566 | 116.1308447 | 107.7177541 | 174.1710941 | 148.3313638 | 171.9174537 | -0.579115 | 7.11653 | 6.33E-05 | 0.00457 |
| ENSG00000182492 | BGN | 107.2231566 | 130.3625659 | 166.2096528 | 79.08046893 | 88.60061328 | 75.5008402 | 0.72727 | 6.76095 | 6.51E-05 | 0.00465 |
| ENSG00000102858 | MGRN1 | 100.6965297 | 106.4532743 | 102.5056047 | 52.39687512 | 57.24196925 | 79.5819667 | 0.711184 | 6.39131 | 6.54E-05 | 0.00465 |
| ENSG00000130037 | KCNA5 | 13.51944149 | 13.66245232 | 20.84859756 | 28.62421881 | 35.83845032 | 29.07802629 | -0.970189 | 4.62752 | 6.57E-05 | 0.00465 |

| | | | | | | | | | | | |
|---|---|---|---|---|---|---|---|---|---|---|---|
| ENSG00000072163 | LIMS2 | 54.07776594 | 47.81858311 | 56.17538787 | 77.62500018 | 77.64997568 | 110.1904154 | -0.745426 | 6.16547 | 6.86E-05 | 0.00483 |
| ENSG00000251596 | HADHAP1 | 137.9915407 | 127.5162216 | 116.9837974 | 65.98125015 | 64.70831307 | 99.47745837 | 0.735121 | 6.68632 | 7.04E-05 | 0.00493 |
| ENSG00000107263 | RAPGEF1 | 55.01014122 | 105.8840055 | 90.92305047 | 41.72343759 | 42.30928162 | 50.5039404 | 0.897899 | 6.0202 | 7.26E-05 | 0.00504 |
| ENSG00000185551 | NR2F2 | 30.30219643 | 35.86393733 | 40.5389397 | 66.4664064 | 59.23299427 | 53.05464446 | -0.752424 | 5.60641 | 7.29E-05 | 0.00504 |
| ENSG00000075239 | ACAT1 | 551.4999751 | 523.15807 | 489.3629149 | 352.2234383 | 379.7880221 | 379.0346234 | 0.493928 | 8.80336 | 7.33E-05 | 0.00504 |
| ENSG00000159640 | ACE | 13.98562912 | 26.75563579 | 26.63987466 | 10.67343752 | 9.955125088 | 5.101408121 | 1.364556 | 4.01979 | 7.54E-05 | 0.00514 |
| ENSG00000123240 | OPTN | 23.77556951 | 27.89417348 | 30.11464092 | 13.09921878 | 15.92820014 | 13.77380193 | 0.923471 | 4.43231 | 7.90E-05 | 0.00536 |
| ENSG00000175602 | CCDC85B | 34.03169753 | 29.03271117 | 28.9563855 | 64.04062514 | 45.29581915 | 50.5039404 | -0.79072 | 5.43416 | 8.18E-05 | 0.00552 |
| ENSG00000165688 | PMPCA | 66.19864451 | 60.91176658 | 54.43800474 | 33.47578133 | 34.84293781 | 42.34168741 | 0.718744 | 5.63326 | 8.66E-05 | 0.00578 |
| ENSG00000135218 | CD36 | 202.7916223 | 255.6017121 | 180.6878455 | 99.45703148 | 107.5153509 | 162.2247783 | 0.791846 | 7.39888 | 8.67E-05 | 0.00578 |
| ENSG00000060138 | YBX3 | 44.75401319 | 30.17124887 | 45.75108909 | 64.04062514 | 68.19260685 | 69.88929126 | -0.738313 | 5.78226 | 8.77E-05 | 0.00581 |
| ENSG00000161714 | PLCD3 | 4.661876374 | 5.123419619 | 2.31651084 | 8.247656269 | 13.43941887 | 15.30422436 | -1.578873 | 3.22005 | 8.92E-05 | 0.00587 |
| ENSG00000130347 | RTN4IP1 | 61.0705805 | 55.21907811 | 55.59626016 | 35.41640633 | 40.3182566 | 26.01718142 | 0.756158 | 5.54029 | 9.88E-05 | 0.00647 |
| ENSG00000175048 | ZDHHC14 | 10.72231566 | 9.108301545 | 7.52866023 | 3.881250009 | 2.488781272 | 2.550704061 | 1.605138 | 2.81551 | 0.0001 | 0.00654 |
| ENSG00000116497 | S100PBP | 2.330938187 | 1.70780654 | 1.73738313 | 9.703125022 | 8.959612579 | 4.591267309 | -1.960668 | 2.58044 | 0.0001 | 0.00658 |
| ENSG00000138411 | HECW2 | 13.98562912 | 11.38537693 | 12.74080962 | 28.13906256 | 19.91025018 | 27.54760386 | -0.97768 | 4.33183 | 0.00011 | 0.00698 |
| ENSG00000118680 | MYL12B | 449.8710701 | 375.1481699 | 445.349209 | 292.5492194 | 299.6492651 | 300.4729384 | 0.50955 | 8.49753 | 0.00011 | 0.00703 |
| ENSG00000135439 | AGAP2 | 9.789940386 | 8.539032698 | 10.42429878 | 24.25781255 | 14.43493138 | 26.52732223 | -1.174772 | 4.07253 | 0.00011 | 0.00703 |
| ENSG00000115307 | AUP1 | 110.4864701 | 105.8840055 | 104.2429878 | 159.1312504 | 157.2909764 | 148.4509763 | -0.534773 | 7.04582 | 0.00011 | 0.00717 |
| ENSG00000147123 | NDUFB11 | 108.6217195 | 75.71275659 | 127.4080962 | 170.7750004 | 162.7662952 | 170.8971721 | -0.693491 | 7.10095 | 0.00012 | 0.00726 |
| ENSG00000143772 | ITPKB | 31.70075934 | 61.48103543 | 52.1214939 | 29.59453132 | 23.39454396 | 16.8346468 | 1.044997 | 5.18842 | 0.00012 | 0.00726 |
| ENSG00000120802 | TMPO | 18.18131786 | 14.80099001 | 14.47819275 | 33.96093758 | 26.38108148 | 28.56788548 | -0.893447 | 4.58185 | 0.00012 | 0.00741 |
| ENSG00000179388 | EGR3 | 3.263313462 | 2.846344233 | 4.63302168 | 0.485156251 | 0.497756254 | 0 | 3.193327 | 1.54697 | 0.00012 | 0.00741 |
| ENSG00000099622 | CIRBP | 34.96407281 | 22.20148501 | 35.32679031 | 49.48593761 | 48.78011293 | 85.70365644 | -0.985366 | 5.56394 | 0.00012 | 0.00741 |
| ENSG00000187486 | KCNJ11 | 133.3296643 | 108.1610808 | 131.4619902 | 74.71406267 | 87.10734452 | 83.66309319 | 0.605037 | 6.70021 | 0.00012 | 0.00764 |
| ENSG00000084774 | CAD | 27.97125825 | 25.61709809 | 35.90591802 | 16.01015629 | 14.43493138 | 17.34478761 | 0.898928 | 4.56856 | 0.00013 | 0.00789 |
| ENSG00000256356 | HSPA8P5 | 227.4995671 | 210.6294732 | 188.7956335 | 156.7054691 | 141.3627762 | 118.3526684 | 0.590759 | 7.44997 | 0.00013 | 0.008 |
| ENSG00000125875 | TBC1D20 | 22.3770066 | 41.5566258 | 27.79813008 | 15.52500004 | 14.93268763 | 13.77380193 | 1.038725 | 4.54815 | 0.00014 | 0.00829 |
| ENSG00000135766 | EGLN1 | 89.50802639 | 92.79082199 | 79.91962398 | 57.24843763 | 51.76665046 | 61.72703827 | 0.619832 | 6.19086 | 0.00014 | 0.00844 |
| ENSG00000005339 | CREBBP | 76.92096017 | 93.36009083 | 99.60996612 | 134.3882816 | 129.4166261 | 141.309005 | -0.590296 | 6.82784 | 0.00014 | 0.00844 |
| ENSG00000063660 | GPC1 | 329.128472 | 288.6193052 | 395.5442259 | 178.0523442 | 158.2864889 | 265.7833631 | 0.750969 | 8.07653 | 0.00015 | 0.00883 |
| ENSG00000244019 | RPS2P24 | 0.932375275 | 2.846344233 | 3.47476626 | 10.67343752 | 5.973075053 | 9.182534619 | -1.86007 | 2.72127 | 0.00015 | 0.00895 |
| ENSG00000128052 | KDR | 142.6534171 | 169.6421163 | 139.5697781 | 76.65468767 | 118.9637448 | 83.66309319 | 0.693332 | 6.93828 | 0.00015 | 0.00904 |
| ENSG00000099822 | HCN2 | 11.65469094 | 10.24683924 | 12.74080962 | 18.92109379 | 23.89230021 | 24.9968998 | -0.963011 | 4.18664 | 0.00015 | 0.00904 |
| ENSG00000239474 | KLHL41 | 47.08495138 | 61.48103543 | 55.01713245 | 30.07968757 | 37.82947533 | 31.11858954 | 0.717752 | 5.47732 | 0.00016 | 0.00909 |
| ENSG00000184304 | PRKD1 | 5.594251649 | 8.539032698 | 5.7912771 | 16.49531254 | 13.43941887 | 15.30422436 | -1.188145 | 3.58208 | 0.00016 | 0.0092 |
| ENSG00000109501 | WFS1 | 93.70371512 | 92.79082199 | 99.03083841 | 72.28828141 | 48.78011293 | 60.19661583 | 0.653558 | 6.29747 | 0.00016 | 0.00927 |
| ENSG00000131791 | PRKAB2 | 8.857565111 | 11.38537693 | 6.94953252 | 2.425781255 | 0.995512509 | 4.591267309 | 1.746204 | 2.7651 | 0.00017 | 0.00946 |
| ENSG00000084754 | HADHA | 534.2510325 | 536.8205223 | 498.0498306 | 293.0343757 | 299.1515089 | 429.5385638 | 0.619346 | 8.75696 | 0.00017 | 0.00946 |
| ENSG00000238008 | COX6CP10 | 10.72231566 | 2.277075386 | 6.94953252 | 18.92109379 | 14.93268763 | 18.36506924 | -1.342426 | 3.73114 | 0.00017 | 0.00956 |
| ENSG00000019582 | CD74 | 16.78275495 | 21.06294732 | 19.11121443 | 7.277343766 | 10.9506376 | 9.182534619 | 1.044481 | 3.89944 | 0.00017 | 0.00971 |
| ENSG00000067225 | PKM | 1734.218011 | 1592.814233 | 1469.826128 | 1216.286722 | 1094.068247 | 1172.303586 | 0.462125 | 10.4314 | 0.00018 | 0.00982 |
| ENSG00000182324 | KCNJ14 | 2.330938187 | 3.984881926 | 5.21214939 | 0 | 0 | 1.020281624 | 3.27274 | 1.59124 | 0.00018 | 0.00983 |
| ENSG00000173991 | TCAP | 275.9830814 | 191.2743324 | 236.8632334 | 709.2984391 | 355.8957219 | 320.8785708 | -0.976383 | 8.4509 | 0.00018 | 0.00988 |
| ENSG00000228224 | NACAP1 | 48.94970193 | 44.97223888 | 51.54236619 | 83.44687519 | 73.17016939 | 68.86900964 | -0.632349 | 5.97788 | 0.00018 | 0.0099 |
| ENSG00000175662 | TOM1L2 | 35.89644808 | 35.86393733 | 41.11806741 | 23.28750005 | 20.40800643 | 22.44619573 | 0.766206 | 4.9402 | 0.00018 | 0.00996 |
| ENSG00000259330 | INAFM2 | 25.17413242 | 29.03271117 | 35.32679031 | 75.68437517 | 44.30030664 | 44.93295472 | -0.905232 | 5.45732 | 0.00018 | 0.00996 |
| ENSG00000048740 | CELF2 | 68.5295827 | 62.61957312 | 45.75108909 | 107.2195315 | 92.08490706 | 85.19351563 | -0.679529 | 6.28947 | 0.00019 | 0.01055 |
| ENSG00000197226 | TBC1D9B | 77.85333545 | 78.55910082 | 83.97351795 | 49.97109386 | 51.2688942 | 57.13577096 | 0.600969 | 6.07269 | 0.0002 | 0.01058 |
| ENSG00000111642 | CHD4 | 20.51225605 | 17.0780654 | 18.53208672 | 37.84218759 | 36.33620657 | 27.03746304 | -0.844431 | 4.77834 | 0.0002 | 0.0106 |
| ENSG00000152558 | TMEM123 | 4.661876374 | 9.108301545 | 6.37040481 | 1.455468753 | 2.986537526 | 0 | 2.108712 | 2.32098 | 0.0002 | 0.01078 |
| ENSG00000231006 | RPL7P32 | 79.25189836 | 43.83370118 | 68.91619749 | 112.5562503 | 102.5377884 | 103.5585849 | -0.724266 | 6.43391 | 0.0002 | 0.01078 |
| ENSG00000111199 | TRPV4 | 1.86475055 | 5.123419619 | 3.47476626 | 9.703125022 | 8.461856324 | 11.73323868 | -1.535176 | 2.96292 | 0.0002 | 0.01085 |
| ENSG00000242262 | RP11-100N21.[illegible] | 287.1715847 | 250.4782925 | 272.7691514 | 412.8679697 | 484.8145918 | 328.530683 | -0.596913 | 8.4125 | 0.00021 | 0.01109 |
| ENSG00000205476 | CCDC85C | 11.65469094 | 11.38537693 | 23.74423611 | 5.336718762 | 6.470831307 | 5.101408121 | 1.445507 | 3.51243 | 0.00021 | 0.0111 |
| ENSG00000107331 | ABCA2 | 67.59720743 | 75.14348774 | 75.2866023 | 49.97109386 | 40.3182566 | 49.99379959 | 0.632786 | 5.91985 | 0.00021 | 0.01115 |
| ENSG00000243964 | RPL23AP65 | 11.65469094 | 9.677570391 | 16.21557588 | 20.8617188 | 32.85191279 | 23.46647736 | -1.040799 | 4.34053 | 0.00021 | 0.01122 |
| ENSG00000065989 | PDE4A | 34.49788517 | 53.51127157 | 32.43115176 | 19.8914063 | 22.39903145 | 22.44619573 | 0.89009 | 4.98421 | 0.00022 | 0.01169 |
| ENSG00000184792 | OSBP2 | 27.97125825 | 27.89417348 | 33.01027947 | 12.61406253 | 13.43941887 | 20.9157733 | 0.918182 | 4.55333 | 0.00023 | 0.01189 |
| ENSG00000013441 | CLK1 | 111.885033 | 104.7454678 | 110.6133926 | 146.5171878 | 165.7528327 | 156.6132293 | -0.51766 | 7.06477 | 0.00023 | 0.01198 |
| ENSG00000147536 | GINS4 | 2.330938187 | 1.138537693 | 1.73738313 | 7.277343766 | 5.475318798 | 6.121689746 | -1.791274 | 2.3593 | 0.00024 | 0.01226 |
| ENSG00000128918 | ALDH1A2 | 6.992814561 | 16.50879655 | 15.05732046 | 3.396093758 | 4.479806289 | 5.101408121 | 1.53987 | 3.22504 | 0.00024 | 0.01237 |
| ENSG00000014216 | CAPN1 | 21.91081896 | 25.04782925 | 17.95295901 | 13.58437503 | 9.457368833 | 9.182534619 | 1.003948 | 4.09766 | 0.00024 | 0.01239 |
| ENSG00000072954 | TMEM38A | 151.0447945 | 162.2416213 | 150.5732046 | 103.3382815 | 78.14773194 | 119.8830909 | 0.621947 | 7.004 | 0.00024 | 0.01239 |
| ENSG00000071242 | RPS6KA2 | 11.65469094 | 14.23172116 | 14.47819275 | 6.307031264 | 6.968587561 | 4.591267309 | 1.154253 | 3.40572 | 0.00025 | 0.01248 |
| ENSG00000196531 | NACA | 241.4851962 | 179.3196867 | 244.3918936 | 354.6492196 | 367.3441157 | 286.6991364 | -0.599773 | 8.13074 | 0.00025 | 0.0126 |
| ENSG00000124177 | CHD6 | 34.49788517 | 42.12589464 | 36.48504573 | 87.3281252 | 46.29133166 | 67.3385872 | -0.831663 | 5.74289 | 0.00026 | 0.01274 |
| ENSG00000151929 | BAG3 | 256.8693882 | 212.3372798 | 257.1327032 | 155.7351566 | 155.7977076 | 187.2216781 | 0.543299 | 7.68002 | 0.00026 | 0.01287 |
| ENSG00000101871 | MID1 | 7.459002199 | 7.969763851 | 9.84517107 | 2.910937507 | 1.991025018 | 3.570985685 | 1.547718 | 2.70988 | 0.00026 | 0.01294 |
| ENSG00000125952 | MAX | 62.00295578 | 58.06542235 | 63.12492039 | 117.892969 | 74.1656819 | 102.0281624 | -0.682356 | 6.33585 | 0.00029 | 0.01407 |
| ENSG00000163110 | PDLIM5 | 450.8034454 | 473.6316803 | 445.349209 | 316.807032 | 258.335496 | 364.7506807 | 0.543394 | 8.59159 | 0.00029 | 0.01407 |
| ENSG00000113578 | FGF1 | 22.84319423 | 43.26443234 | 29.53551321 | 16.01015629 | 18.41698141 | 12.7535203 | 1.005988 | 4.61647 | 0.00029 | 0.01408 |
| ENSG00000012232 | EXTL3 | 26.1065077 | 47.81858311 | 45.17196138 | 18.43593754 | 23.39454396 | 20.40563249 | 0.923754 | 4.94787 | 0.0003 | 0.01437 |
| ENSG00000047849 | MAP4 | 162.6994855 | 182.7352997 | 178.9504624 | 129.0515628 | 105.5243259 | 127.535203 | 0.532537 | 7.21485 | 0.0003 | 0.01437 |
| ENSG00000154188 | ANGPT1 | 21.91081896 | 14.23172116 | 12.74080962 | 24.25781255 | 35.83845032 | 35.70985685 | -0.947107 | 4.66503 | 0.0003 | 0.01437 |
| ENSG00000189056 | RELN | 9.789940386 | 10.24683924 | 7.52866023 | 3.396093758 | 1.493268763 | 4.591267309 | 1.529809 | 2.8348 | 0.0003 | 0.01437 |
| ENSG00000169567 | HINT1 | 108.6217195 | 117.2693824 | 120.4585637 | 159.6164066 | 179.1922516 | 156.1030885 | -0.516557 | 7.14319 | 0.0003 | 0.01437 |
| ENSG00000250500 | RP11-119H12.[illegible] | 6.992814561 | 8.539032698 | 6.37040481 | 1.940625004 | 1.991025018 | 3.060844873 | 1.623229 | 2.52665 | 0.0003 | 0.0144 |
| ENSG00000213585 | VDAC1 | 325.3989709 | 351.2388783 | 309.2541971 | 217.8351567 | 230.4611458 | 257.1109693 | 0.483133 | 8.14331 | 0.00031 | 0.01489 |
| ENSG00000226313 | MTCYBP12 | 2130.943691 | 2273.659773 | 2533.683731 | 1317.684378 | 1764.048166 | 1721.2151 | 0.530581 | 10.9349 | 0.00032 | 0.01491 |
| ENSG00000171033 | PKIA | 27.50507061 | 44.97223888 | 24.90249153 | 16.01015629 | 14.43493138 | 18.87521005 | 0.975192 | 4.65799 | 0.00032 | 0.01491 |

| | | | | | | | | | | | |
|---|---|---|---|---|---|---|---|---|---|---|---|
| ENSG00000118922 | KLF12 | 13.51944149 | 14.23172116 | 11.00342649 | 27.65390631 | 22.39903145 | 22.44619573 | -0.897555 | 4.30143 | 0.00032 | 0.01491 |
| ENSG00000101557 | USP14 | 63.40151869 | 78.55910082 | 74.70747459 | 42.20859385 | 48.78011293 | 48.97351797 | 0.626091 | 5.91206 | 0.00032 | 0.01491 |
| ENSG00000167680 | SEMA6B | 1.86475055 | 0 | 1.73738313 | 3.396093758 | 6.968587561 | 7.652112182 | -2.217143 | 2.23463 | 0.00032 | 0.01495 |
| ENSG00000157184 | CPT2 | 61.0705805 | 59.20396004 | 56.17538787 | 33.96093758 | 33.34966904 | 44.38225066 | 0.662224 | 5.61243 | 0.00032 | 0.01495 |
| ENSG00000154122 | ANKH | 146.3829182 | 136.0552543 | 133.778501 | 91.69453146 | 94.07593208 | 104.5788665 | 0.521268 | 6.89094 | 0.00032 | 0.01496 |
| ENSG00000175920 | DOK7 | 13.51944149 | 11.95464578 | 10.42429878 | 5.821875013 | 2.488781272 | 6.121689746 | 1.312505 | 3.22705 | 0.00033 | 0.01504 |
| ENSG00000131477 | RAMP2 | 15.85037967 | 11.38537693 | 19.69034214 | 38.32734384 | 20.40800643 | 38.77070172 | -1.05242 | 4.65873 | 0.00033 | 0.01514 |
| ENSG00000109079 | TNFAIP1 | 29.8360088 | 35.29466849 | 31.85202405 | 18.92109379 | 19.91025018 | 18.36506924 | 0.755131 | 4.7313 | 0.00033 | 0.01518 |
| ENSG00000164754 | RAD21 | 38.69357391 | 45.54150772 | 33.58940718 | 63.07031264 | 61.72177554 | 59.17633421 | -0.642404 | 5.68582 | 0.00033 | 0.01518 |
| ENSG00000152137 | HSPB8 | 340.3169753 | 285.2036921 | 280.8769393 | 191.1515629 | 169.2371265 | 242.8270266 | 0.588864 | 7.98043 | 0.00034 | 0.0152 |
| ENSG00000198812 | LRRC10 | 50.34826484 | 30.17124887 | 49.80498306 | 100.427344 | 48.78011293 | 113.2512603 | -1.005908 | 6.06135 | 0.00035 | 0.0158 |
| ENSG00000170989 | S1PR1 | 32.63313462 | 34.15613079 | 42.85545054 | 64.5257814 | 49.27786918 | 63.76760152 | -0.700397 | 5.61466 | 0.00035 | 0.01582 |
| ENSG00000128928 | IVD | 158.9699844 | 153.1333197 | 148.2566938 | 101.8828127 | 87.60510077 | 121.9236541 | 0.565 | 7.01695 | 0.00035 | 0.01582 |
| ENSG00000171823 | FBXL14 | 15.85037967 | 14.80099001 | 18.53208672 | 8.73281252 | 6.470831307 | 8.672393807 | 1.033494 | 3.71011 | 0.00036 | 0.01597 |
| ENSG00000167588 | GPD1 | 53.14539067 | 55.78834696 | 69.4953252 | 32.99062507 | 33.8474253 | 43.36196903 | 0.691576 | 5.61119 | 0.00036 | 0.01597 |
| ENSG00000019144 | PHLDB1 | 132.8634767 | 133.7781789 | 126.2498408 | 96.54609397 | 85.61407575 | 92.84562781 | 0.514674 | 6.81014 | 0.00036 | 0.01597 |
| ENSG00000236480 | PKMP1 | 137.525353 | 137.7630609 | 130.8828625 | 90.72421896 | 88.60061328 | 104.0687257 | 0.520029 | 6.85566 | 0.00037 | 0.01607 |
| ENSG00000084628 | NKAIN1 | 10.25612802 | 11.95464578 | 8.10778794 | 4.851562511 | 1.493268763 | 4.591267309 | 1.456063 | 2.96809 | 0.00037 | 0.01609 |
| ENSG00000134240 | HMGCS2 | 13.05325385 | 10.24683924 | 69.4953252 | 5.336718762 | 5.973075053 | 9.692675431 | 2.129964 | 4.27197 | 0.00037 | 0.01615 |
| ENSG00000157613 | CREB3L1 | 7.459002199 | 8.539032698 | 9.26604336 | 2.910937507 | 2.488781272 | 3.570985685 | 1.466456 | 2.72857 | 0.00037 | 0.01623 |
| ENSG00000124181 | PLCG1 | 94.16990276 | 99.62204814 | 82.23613482 | 136.8140628 | 122.9457948 | 140.7988642 | -0.535755 | 6.83242 | 0.00038 | 0.01644 |
| ENSG00000113441 | LNPEP | 54.54395358 | 50.66492734 | 52.70062161 | 77.13984392 | 78.64548819 | 79.07182588 | -0.57025 | 6.05846 | 0.00038 | 0.01645 |
| ENSG00000167291 | TBC1D16 | 20.97844368 | 29.60198002 | 32.43115176 | 14.55468753 | 12.44390636 | 16.8346468 | 0.907271 | 4.45298 | 0.00038 | 0.01645 |
| ENSG00000156427 | FGF18 | 2.797125825 | 6.261957312 | 2.89563855 | 12.12890628 | 16.42595639 | 7.14197137 | -1.583668 | 3.18032 | 0.00039 | 0.01657 |
| ENSG00000100266 | PACSIN2 | 183.6779291 | 165.0879655 | 167.9470359 | 93.63515646 | 124.9368198 | 130.0859071 | 0.569279 | 7.18116 | 0.00039 | 0.01666 |
| ENSG00000013561 | RNF14 | 20.51225605 | 21.06294732 | 13.31993733 | 9.703125022 | 9.457368833 | 6.631830558 | 1.088722 | 3.85082 | 0.0004 | 0.01709 |
| ENSG00000234152 | ELOBP1 | 3.263313462 | 4.554150772 | 2.89563855 | 6.307031264 | 11.94615011 | 13.26366112 | -1.540884 | 3.02379 | 0.00041 | 0.01748 |
| ENSG00000102683 | SGCG | 74.12383435 | 75.71275659 | 67.75794207 | 43.17890635 | 27.37659399 | 56.62563015 | 0.775662 | 5.8658 | 0.00043 | 0.01787 |
| ENSG00000109756 | RAPGEF2 | 60.13820523 | 46.11077657 | 56.75451558 | 84.41718769 | 80.63651321 | 80.60224832 | -0.586906 | 6.11567 | 0.00043 | 0.01787 |
| ENSG00000236476 | RP11-8J9.5 | 8.857565111 | 4.554150772 | 5.7912771 | 15.03984378 | 11.94615011 | 17.34478761 | -1.168643 | 3.55762 | 0.00044 | 0.01819 |
| ENSG00000161011 | SQSTM1 | 239.154258 | 221.4455813 | 172.5800576 | 124.2000003 | 121.9502823 | 166.3059048 | 0.620362 | 7.4531 | 0.00044 | 0.01836 |
| ENSG00000143627 | PKLR | 21.91081896 | 25.04782925 | 27.21900237 | 11.15859378 | 12.94166261 | 16.32450599 | 0.868827 | 4.31765 | 0.00044 | 0.01844 |
| ENSG00000100478 | AP4S1 | 28.43744588 | 18.21660309 | 28.37725779 | 46.57500011 | 43.80255039 | 37.24027929 | -0.759173 | 5.12964 | 0.00045 | 0.01844 |
| ENSG00000213923 | CSNK1E | 0.466187637 | 2.846344233 | 2.31651084 | 5.821875013 | 9.457368833 | 5.611548934 | -1.921052 | 2.45531 | 0.00045 | 0.01866 |
| ENSG00000161016 | RPL8 | 463.8566992 | 393.364773 | 451.7196138 | 586.5539076 | 611.7424366 | 582.0706667 | -0.44314 | 9.01154 | 0.00046 | 0.01898 |
| ENSG00000203825 | RP11-744H18. | 31.70075934 | 34.15613079 | 23.74423611 | 16.49531254 | 14.93268763 | 18.87521005 | 0.835478 | 4.60026 | 0.00046 | 0.01898 |
| ENSG00000104852 | SNRNP70 | 39.62594918 | 28.46344233 | 31.27289634 | 59.67421889 | 49.77562544 | 50.5039404 | -0.676389 | 5.47542 | 0.00047 | 0.0191 |
| ENSG00000068724 | TTC7A | 11.1885033 | 11.95464578 | 20.84859756 | 7.762500018 | 2.986537526 | 6.121689746 | 1.359035 | 3.45528 | 0.00047 | 0.01915 |
| ENSG00000122756 | CNTFR | 1.398562912 | 1.138537693 | 2.31651084 | 5.821875013 | 5.475318798 | 6.631830558 | -1.858557 | 2.2871 | 0.00047 | 0.01915 |
| ENSG00000214121 | PRDX1P1 | 65.26626924 | 87.66740237 | 72.39096375 | 45.11953135 | 52.26440671 | 48.97351797 | 0.618108 | 5.97096 | 0.00049 | 0.01963 |
| ENSG00000077942 | FBLN1 | 73.65764671 | 58.06542235 | 90.92305047 | 97.51640647 | 116.9727198 | 147.9408355 | -0.702333 | 6.6246 | 0.00049 | 0.01967 |
| ENSG00000261150 | EPPK1 | 11.65469094 | 19.92440963 | 16.79470359 | 9.703125022 | 6.470831307 | 4.591267309 | 1.193016 | 3.62645 | 0.0005 | 0.01994 |
| ENSG00000162783 | IER5 | 22.84319423 | 10.81610808 | 11.00342649 | 39.29765634 | 25.88332523 | 27.54760386 | -1.027 | 4.60021 | 0.00051 | 0.02019 |
| ENSG00000118515 | SGK1 | 66.19864451 | 63.75811081 | 46.90934451 | 116.4375003 | 86.60958826 | 80.09210751 | -0.675083 | 6.28505 | 0.00051 | 0.0202 |
| ENSG00000163517 | HDAC11 | 13.05325385 | 11.38537693 | 6.94953252 | 3.396093758 | 5.475318798 | 2.550704061 | 1.456234 | 3.02671 | 0.00052 | 0.02055 |
| ENSG00000113558 | SKP1 | 241.0190085 | 222.0148501 | 214.8563804 | 155.7351566 | 178.1967391 | 153.0422436 | 0.478195 | 7.60799 | 0.00053 | 0.02076 |
| ENSG00000115461 | IGFBP5 | 21.44463132 | 27.32490463 | 29.53551321 | 16.49531254 | 11.94615011 | 14.79408355 | 0.842748 | 4.39761 | 0.00053 | 0.02076 |
| ENSG00000168175 | MAPK1IP1L | 12.58706621 | 7.400495005 | 9.26604336 | 16.98046879 | 19.91025018 | 20.9157733 | -0.954738 | 3.97256 | 0.00053 | 0.02076 |
| ENSG00000224004 | ATP5C1P1 | 499.7531473 | 398.4881926 | 439.5579319 | 582.1875013 | 639.1190306 | 625.9427765 | -0.464365 | 9.05558 | 0.00053 | 0.02081 |
| ENSG00000186350 | RXRA | 34.96407281 | 20.49367848 | 34.7476626 | 17.46562504 | 13.93717512 | 16.8346468 | 0.906043 | 4.58783 | 0.00053 | 0.02081 |
| ENSG00000131779 | PEX11B | 25.64032006 | 17.64733424 | 23.74423611 | 12.61406253 | 12.44390636 | 10.71295706 | 0.909178 | 4.17932 | 0.00054 | 0.02081 |
| ENSG00000149972 | CNTN5 | 5.594251649 | 10.81610808 | 2.89563855 | 2.425781255 | 0 | 0.510140812 | 2.619796 | 2.21583 | 0.00054 | 0.02081 |
| ENSG00000138363 | ATIC | 10.25612802 | 10.81610808 | 10.42429878 | 4.36640626 | 1.991025018 | 6.121689746 | 1.324902 | 3.04622 | 0.00054 | 0.02098 |
| ENSG00000163624 | CDS1 | 4.661876374 | 2.277075386 | 2.89563855 | 0.485156251 | 0 | 0.510140812 | 3.109134 | 1.50091 | 0.00055 | 0.02119 |
| ENSG00000162413 | KLHL21 | 48.94970193 | 33.58686195 | 58.49189871 | 27.16875006 | 14.43493138 | 31.11858954 | 0.954741 | 5.18904 | 0.00056 | 0.02126 |
| ENSG00000087274 | ADD1 | 83.91377474 | 139.4708674 | 116.4046697 | 66.4664064 | 69.18811936 | 78.05154426 | 0.664444 | 6.53703 | 0.00056 | 0.02126 |
| ENSG00000172340 | SUCLG2 | 123.0735363 | 127.5162216 | 103.6638601 | 74.22890642 | 78.64548819 | 89.78478294 | 0.54757 | 6.64953 | 0.00056 | 0.0214 |
| ENSG00000153879 | CEBPG | 7.459002199 | 9.677570391 | 6.94953252 | 3.396093758 | 2.986537526 | 2.040563249 | 1.484897 | 2.67 | 0.00057 | 0.02141 |
| ENSG00000264217 | RPL35AP25 | 1.398562912 | 0 | 1.73738313 | 5.336718762 | 5.475318798 | 3.570985685 | -2.108971 | 2.00775 | 0.00058 | 0.02194 |
| ENSG00000151176 | PLBD2 | 33.09932226 | 21.63221617 | 41.11806741 | 13.58437503 | 13.93717512 | 21.42591411 | 0.969853 | 4.64396 | 0.00058 | 0.02194 |
| ENSG00000213862 | CTD-2270N23. | 165.0304236 | 117.2693824 | 156.9436094 | 200.3695317 | 246.8871022 | 205.5867473 | -0.56945 | 7.51801 | 0.00059 | 0.02231 |
| ENSG00000166341 | DCHS1 | 23.77556951 | 39.84881926 | 44.01370596 | 16.98046879 | 23.39454396 | 15.30422436 | 0.937253 | 4.80138 | 0.00061 | 0.02277 |
| ENSG00000113328 | CCNG1 | 351.039291 | 466.8004542 | 357.9009248 | 251.3109381 | 276.7524774 | 290.7802629 | 0.521248 | 8.38015 | 0.00061 | 0.02277 |
| ENSG00000091073 | DTX2 | 1.398562912 | 1.70780654 | 0.57912771 | 6.792187515 | 3.982050035 | 4.591267309 | -1.992421 | 2.09698 | 0.00062 | 0.02285 |
| ENSG00000148985 | PGAP2 | 44.75401319 | 58.63469119 | 48.64672764 | 32.99062507 | 29.86537526 | 33.15915279 | 0.657135 | 5.39777 | 0.00062 | 0.02285 |
| ENSG00000238082 | AC009948.7 | 2.797125825 | 1.70780654 | 3.47476626 | 9.217968771 | 6.470831307 | 8.162252994 | -1.555636 | 2.68163 | 0.00063 | 0.0234 |
| ENSG00000114982 | KANSL3 | 47.08495138 | 68.31226158 | 53.85887703 | 33.47578133 | 32.85191279 | 39.28084254 | 0.674742 | 5.54143 | 0.00064 | 0.02343 |
| ENSG00000217027 | TPT1P4 | 73.65764671 | 50.0956585 | 86.86891565 | 114.4968753 | 126.9278449 | 95.39633187 | -0.676502 | 6.53113 | 0.00066 | 0.02436 |
| ENSG00000033627 | ATP6V0A1 | 90.44040166 | 93.92935968 | 94.39781673 | 64.5257814 | 69.68587561 | 58.6661934 | 0.529532 | 6.31256 | 0.00067 | 0.02437 |
| ENSG00000233762 | AC007969.5 | 360.3630437 | 258.4480563 | 302.8837923 | 424.5117197 | 455.4469728 | 429.028423 | -0.504236 | 8.54341 | 0.00067 | 0.0245 |
| ENSG00000155846 | PPARGC1B | 30.76838407 | 26.18636694 | 19.11121443 | 38.81250009 | 49.27786918 | 39.79098335 | -0.73586 | 5.13932 | 0.0007 | 0.02535 |
| ENSG00000136842 | TMOD1 | 178.0836775 | 212.9065486 | 201.5364431 | 138.2695316 | 151.8156576 | 133.6568928 | 0.481607 | 7.41083 | 0.00071 | 0.02567 |
| ENSG00000116906 | GNPAT | 40.55832446 | 37.00247502 | 31.27289634 | 18.43593754 | 18.91473767 | 27.03746304 | 0.763928 | 4.89853 | 0.00072 | 0.02607 |
| ENSG00000162599 | NFIA | 217.7096267 | 236.2465713 | 249.0249153 | 315.3515632 | 320.0572716 | 311.1858954 | -0.430551 | 8.10877 | 0.00075 | 0.02698 |
| ENSG00000072506 | HSD17B10 | 71.79289616 | 54.64980927 | 85.71090108 | 103.8234377 | 98.05798211 | 126.0047806 | -0.627252 | 6.51067 | 0.00076 | 0.02714 |
| ENSG00000170464 | DNAJC18 | 7.925189836 | 10.81610808 | 9.84517107 | 2.910937507 | 3.982050035 | 4.591267309 | 1.293095 | 2.9223 | 0.00077 | 0.02738 |
| ENSG00000163660 | CCNL1 | 55.94251649 | 42.12589464 | 44.59283367 | 76.65468767 | 77.64997568 | 63.25746071 | -0.601263 | 5.9382 | 0.00077 | 0.0274 |
| ENSG00000092969 | TGFB2 | 6.526626924 | 8.539032698 | 5.7912771 | 3.881250009 | 1.493268763 | 1.020281624 | 1.659287 | 2.45983 | 0.00078 | 0.02763 |
| ENSG00000159069 | FBXW5 | 45.68638847 | 52.37273388 | 60.22928184 | 30.56484382 | 34.84293781 | 36.21999766 | 0.633217 | 5.46293 | 0.00078 | 0.02776 |
| ENSG00000233383 | RP11-324F21.1 | 2.797125825 | 1.138537693 | 2.31651084 | 5.821875013 | 5.973075053 | 7.652112182 | -1.575533 | 2.43028 | 0.00079 | 0.02806 |
| ENSG00000196296 | ATP2A1 | 777.1347916 | 725.2485105 | 718.6974881 | 502.6218761 | 358.3845032 | 610.1284113 | 0.594544 | 9.26732 | 0.0008 | 0.02815 |

| | | | | | | | | | | | |
|---|---|---|---|---|---|---|---|---|---|---|---|
| ENSG00000091542 | ALKBH5 | 29.8360088 | 37.00247502 | 46.3302168 | 19.8914063 | 20.40800643 | 25.50704061 | 0.772845 | 4.93428 | 0.0008 | 0.02815 |
| ENSG00000148400 | NOTCH1 | 32.16694698 | 39.84881926 | 49.22585535 | 22.8023438 | 27.37659399 | 21.93605492 | 0.738825 | 5.04371 | 0.00081 | 0.02838 |
| ENSG00000126882 | FAM78A | 13.51944149 | 16.50879655 | 12.16168191 | 4.851562511 | 9.457368833 | 4.591267309 | 1.148646 | 3.47294 | 0.00081 | 0.02838 |
| ENSG00000102401 | ARMCX3 | 9.789940386 | 12.52391462 | 8.10778794 | 26.19843756 | 24.39005646 | 13.26366112 | -1.068785 | 4.0786 | 0.00083 | 0.02863 |
| ENSG00000240463 | RPS19P3 | 181.346991 | 106.4532743 | 131.4619902 | 214.439063 | 240.4162709 | 195.3839311 | -0.628861 | 7.4892 | 0.00083 | 0.02863 |
| ENSG00000196876 | SCN8A | 10.25612802 | 7.400495005 | 12.74080962 | 5.336718762 | 3.484293781 | 3.060844873 | 1.334938 | 2.99934 | 0.00083 | 0.02863 |
| ENSG00000100226 | GTPBP1 | 35.43026044 | 43.26443234 | 42.27632283 | 23.28750005 | 25.38556897 | 27.03746304 | 0.669526 | 5.07048 | 0.00083 | 0.02863 |
| ENSG00000123131 | PRDX4 | 14.45181676 | 15.9395277 | 12.74080962 | 32.02031257 | 26.87883774 | 19.89549167 | -0.868204 | 4.42653 | 0.00083 | 0.02865 |
| ENSG00000023171 | GRAMD1B | 33.09932226 | 37.00247502 | 38.22242886 | 60.15937514 | 45.29581915 | 64.78788314 | -0.65512 | 5.57145 | 0.00085 | 0.02897 |
| ENSG00000196440 | ARMCX4 | 2.330938187 | 2.846344233 | 1.73738313 | 13.09921878 | 5.973075053 | 5.611548934 | -1.810189 | 2.67998 | 0.00086 | 0.02945 |
| ENSG00000165458 | INPPL1 | 231.6952558 | 246.4934105 | 239.1797442 | 189.2109379 | 165.7528327 | 173.9580169 | 0.438753 | 7.70447 | 0.00087 | 0.02947 |
| ENSG00000135631 | RAB11FIP5 | 24.70794478 | 29.60198002 | 32.43115176 | 15.03984378 | 15.43044389 | 19.89549167 | 0.775921 | 4.56503 | 0.00088 | 0.02947 |
| ENSG00000178209 | PLEC | 610.705805 | 576.1000727 | 748.812129 | 473.0273448 | 426.0793537 | 490.2453205 | 0.477996 | 9.11622 | 0.00088 | 0.02947 |
| ENSG00000103269 | RHBDL1 | 6.992814561 | 6.261957312 | 9.26604336 | 2.910937507 | 2.986537526 | 2.040563249 | 1.471877 | 2.5899 | 0.00088 | 0.02947 |
| ENSG00000146926 | ASB10 | 38.22738627 | 30.17124887 | 31.85202405 | 17.95078129 | 20.90576268 | 21.93605492 | 0.729024 | 4.7982 | 0.00088 | 0.02947 |
| ENSG00000148411 | NACC2 | 8.391377474 | 8.539032698 | 10.42429878 | 2.425781255 | 2.488781272 | 5.611548934 | 1.364639 | 2.85432 | 0.00088 | 0.02947 |
| ENSG00000101335 | MYL9 | 444.2768185 | 396.2111172 | 425.0797391 | 311.4703132 | 322.5460528 | 317.817726 | 0.411538 | 8.53335 | 0.00088 | 0.02947 |
| ENSG00000154518 | ATP5G3 | 5034.826484 | 3779.945141 | 4960.228836 | 6719.414078 | 7461.366253 | 5619.711187 | -0.523432 | 12.4505 | 0.00088 | 0.02947 |
| ENSG00000129682 | FGF13 | 61.53676814 | 53.51127157 | 64.28317581 | 87.81328145 | 81.13426946 | 90.80506456 | -0.53292 | 6.21635 | 0.00089 | 0.02952 |
| ENSG00000186815 | TPCN1 | 17.24894258 | 23.34002271 | 30.69376863 | 10.67343752 | 14.93268763 | 10.71295706 | 0.956257 | 4.22474 | 0.00089 | 0.02952 |
| ENSG00000122550 | KLHL7 | 71.32670853 | 72.86641236 | 60.80840955 | 29.59453132 | 48.78011293 | 49.99379959 | 0.678082 | 5.81881 | 0.00093 | 0.03069 |
| ENSG00000244244 | RPS3AP42 | 7.925189836 | 5.123419619 | 7.52866023 | 17.46562504 | 14.93268763 | 11.22309787 | -1.068917 | 3.56886 | 0.00094 | 0.03087 |
| ENSG00000115756 | HPCAL1 | 13.05325385 | 14.80099001 | 7.52866023 | 19.40625004 | 20.90576268 | 28.56788548 | -0.946777 | 4.21228 | 0.00094 | 0.03087 |
| ENSG00000115268 | RPS15 | 110.0202824 | 92.79082199 | 128.5663516 | 159.1312504 | 151.8156576 | 162.7349191 | -0.51561 | 7.08048 | 0.00094 | 0.03087 |
| ENSG00000101542 | CDH20 | 0 | 0 | 1.15825542 | 3.396093758 | 5.973075053 | 1.530422436 | -3.131406 | 1.62454 | 0.00095 | 0.0309 |
| ENSG00000138459 | SLC35A5 | 6.992814561 | 5.123419619 | 4.05389397 | 1.455468753 | 1.991025018 | 0.510140812 | 1.994745 | 2.12913 | 0.00095 | 0.03108 |
| ENSG00000111540 | RAB5B | 53.6115783 | 83.11325159 | 50.96323848 | 38.32734384 | 33.8474253 | 42.34168741 | 0.708996 | 5.6764 | 0.00096 | 0.03116 |
| ENSG00000162616 | DNAJB4 | 160.3685473 | 155.4103951 | 152.8897154 | 106.249219 | 97.0624696 | 128.0453438 | 0.501141 | 7.06855 | 0.00097 | 0.03116 |
| ENSG00000139914 | FITM1 | 86.71090056 | 57.4961535 | 82.23613482 | 111.5859378 | 111.9951572 | 113.2512603 | -0.568276 | 6.572 | 0.00097 | 0.03116 |
| ENSG00000114988 | LMAN2L | 22.3770066 | 28.46344233 | 22.58598069 | 14.55468753 | 9.457368833 | 16.32450599 | 0.858024 | 4.30976 | 0.00099 | 0.03175 |
| ENSG00000272325 | NUDT3 | 8.857565111 | 7.400495005 | 8.68691565 | 4.851562511 | 0.995512509 | 3.060844873 | 1.463598 | 2.72582 | 0.001 | 0.03197 |
| ENSG00000164068 | RNF123 | 88.57565111 | 106.4532743 | 130.8828625 | 70.83281266 | 72.67241314 | 74.99069939 | 0.572096 | 6.51468 | 0.001 | 0.03197 |
| ENSG00000133112 | TPT1 | 408.3803704 | 363.7627929 | 406.5476524 | 556.9593763 | 631.6526868 | 463.2078574 | -0.486634 | 8.88568 | 0.001 | 0.03203 |
| ENSG00000135924 | DNAJB2 | 72.2590838 | 50.0956585 | 56.75451558 | 39.29765634 | 39.82050035 | 37.24027929 | 0.628704 | 5.65216 | 0.00101 | 0.03209 |
| ENSG00000112651 | MRPL2 | 79.718086 | 67.17372389 | 68.91619749 | 110.130469 | 107.5153509 | 92.84562781 | -0.520919 | 6.47517 | 0.00101 | 0.03209 |
| ENSG00000177697 | CD151 | 77.38714781 | 85.39032698 | 72.39096375 | 50.45625011 | 38.82498784 | 63.25746071 | 0.624794 | 6.03286 | 0.00105 | 0.0332 |
| ENSG00000144712 | CAND2 | 57.80726704 | 54.64980927 | 65.44143123 | 33.96093758 | 31.85640028 | 47.95323634 | 0.64485 | 5.62802 | 0.00106 | 0.03353 |
| ENSG00000149654 | CDH22 | 6.992814561 | 6.261957312 | 12.16168191 | 4.36640626 | 2.488781272 | 1.530422436 | 1.55748 | 2.7102 | 0.00107 | 0.03353 |
| ENSG00000132780 | NASP | 22.3770066 | 16.50879655 | 16.21557588 | 32.50546882 | 27.87435025 | 32.13887117 | -0.730839 | 4.69056 | 0.00107 | 0.03353 |
| ENSG00000130821 | SLC6A8 | 121.2087857 | 114.4230382 | 109.4551372 | 70.83281266 | 71.17914438 | 95.39633187 | 0.541743 | 6.61455 | 0.00107 | 0.03353 |
| ENSG00000237793 | RPL18AP16 | 3.263313462 | 3.984881926 | 7.52866023 | 11.15859378 | 8.959612579 | 17.85492843 | -1.372898 | 3.30099 | 0.00107 | 0.03354 |
| ENSG00000073921 | PICALM | 345.4450393 | 371.1632879 | 345.7392429 | 468.6609386 | 464.4065853 | 462.6977166 | -0.39405 | 8.68241 | 0.00109 | 0.03381 |
| ENSG00000129596 | CDO1 | 0.932375275 | 2.277075386 | 39.95981199 | 0.970312502 | 0.995512509 | 1.530422436 | 3.550601 | 3.0405 | 0.00109 | 0.03381 |
| ENSG00000129991 | TNNI3 | 4223.193807 | 3223.200209 | 3830.929802 | 5112.576574 | 5268.749953 | 4873.375178 | -0.435703 | 12.1109 | 0.00116 | 0.03581 |
| ENSG00000235485 | RP1-101G11.2 | 290.4348981 | 322.2061671 | 313.3080911 | 194.5476567 | 179.6900078 | 265.7833631 | 0.532642 | 8.03213 | 0.00116 | 0.03581 |
| ENSG00000124702 | KLHDC3 | 60.60439286 | 63.18884197 | 63.7040481 | 90.72421896 | 79.14324445 | 101.5180216 | -0.533839 | 6.27833 | 0.00116 | 0.03581 |
| ENSG00000188846 | RPL14 | 186.475055 | 161.1030836 | 191.1121443 | 268.2914069 | 268.7883774 | 215.7895635 | -0.482564 | 7.75814 | 0.00117 | 0.03581 |
| ENSG00000119689 | DLST | 430.2911893 | 458.2614215 | 374.6956284 | 304.1929694 | 329.5146404 | 300.4729384 | 0.43557 | 8.52023 | 0.00117 | 0.03594 |
| ENSG00000091986 | CCDC80 | 70.86052089 | 59.20396004 | 62.54579268 | 102.8531252 | 94.57368833 | 82.13267076 | -0.53382 | 6.3209 | 0.00118 | 0.03617 |
| ENSG00000181524 | RPL24P4 | 46.61876374 | 36.43320618 | 44.59283367 | 73.25859392 | 57.73972551 | 61.72703827 | -0.59029 | 5.77132 | 0.0012 | 0.03628 |
| ENSG00000092096 | SLC22A17 | 7.459002199 | 8.539032698 | 7.52866023 | 1.940625004 | 2.986537526 | 4.081126497 | 1.370268 | 2.67023 | 0.0012 | 0.03628 |
| ENSG00000104812 | GYS1 | 92.30515221 | 95.63716622 | 108.2968818 | 62.10000014 | 44.30030664 | 81.62252994 | 0.654925 | 6.3485 | 0.0012 | 0.03628 |
| ENSG00000196547 | MAN2A2 | 20.04606841 | 35.29466849 | 37.64330115 | 15.03984378 | 16.92371265 | 18.36506924 | 0.870916 | 4.61877 | 0.0012 | 0.03632 |
| ENSG00000177508 | IRX3 | 15.85037967 | 11.95464578 | 9.84517107 | 26.68359381 | 18.41698141 | 24.9968998 | -0.87575 | 4.26122 | 0.00122 | 0.03683 |
| ENSG00000185739 | SRL | 317.0075934 | 512.3419619 | 355.0052862 | 280.4203131 | 262.8153023 | 248.4385755 | 0.579849 | 8.3663 | 0.00123 | 0.03699 |
| ENSG00000119421 | NDUFA8 | 134.7282272 | 113.8537693 | 132.6202456 | 187.2703129 | 176.2057141 | 163.2450599 | -0.465176 | 7.25306 | 0.00124 | 0.03705 |
| ENSG00000105810 | CDK6 | 1.398562912 | 3.984881926 | 1.73738313 | 6.307031264 | 7.96410007 | 6.631830558 | -1.561921 | 2.52109 | 0.00124 | 0.0371 |
| ENSG00000221869 | CEBPD | 5.594251649 | 7.969763851 | 9.26604336 | 25.22812506 | 9.457368833 | 18.87521005 | -1.239984 | 3.79226 | 0.00124 | 0.0371 |
| ENSG00000112964 | GHR | 10.72231566 | 14.80099001 | 7.52866023 | 23.28750005 | 27.87435025 | 15.30422436 | -1.005359 | 4.15039 | 0.00125 | 0.0371 |
| ENSG00000213309 | RPL9P18 | 4.661876374 | 2.846344233 | 5.21214939 | 9.217968771 | 9.457368833 | 11.22309787 | -1.21167 | 3.03788 | 0.00125 | 0.0371 |
| ENSG00000225200 | AB019441.29 | 120.7425981 | 91.08301545 | 99.60996612 | 142.6359378 | 182.6765454 | 133.6568928 | -0.555697 | 7.01903 | 0.00128 | 0.03776 |
| ENSG00000111142 | METAP2 | 184.1441168 | 154.8411263 | 165.0513973 | 229.964063 | 255.8467148 | 209.6678738 | -0.462589 | 7.65227 | 0.0013 | 0.03845 |
| ENSG00000108528 | SLC25A11 | 582.2683591 | 560.160545 | 538.5887703 | 427.9078135 | 419.6085224 | 441.2718025 | 0.383688 | 8.95391 | 0.00131 | 0.0386 |
| ENSG00000136158 | SPRY2 | 37.76119863 | 27.89417348 | 32.43115176 | 54.82265637 | 52.26440671 | 44.38225066 | -0.618837 | 5.42066 | 0.00132 | 0.0386 |
| ENSG00000163485 | ADORA1 | 2.797125825 | 3.415613079 | 3.47476626 | 10.18828127 | 6.470831307 | 8.672393807 | -1.382875 | 2.79342 | 0.00132 | 0.0386 |
| ENSG00000165092 | ALDH1A1 | 21.44463132 | 14.23172116 | 53.85887703 | 9.703125022 | 10.45288134 | 15.30422436 | 1.327846 | 4.4243 | 0.00132 | 0.0386 |
| ENSG00000119673 | ACOT2 | 15.85037967 | 15.37025886 | 12.74080962 | 6.307031264 | 4.977562544 | 10.20281624 | 1.036588 | 3.56849 | 0.00132 | 0.0386 |
| ENSG00000141905 | NFIC | 22.84319423 | 37.00247502 | 48.06759993 | 12.61406253 | 14.93268763 | 26.01718142 | 1.002441 | 4.78211 | 0.00136 | 0.03938 |
| ENSG00000008988 | RPS20 | 87.64327584 | 71.72787466 | 71.23270833 | 119.833594 | 103.0355447 | 106.1092889 | -0.507637 | 6.56268 | 0.00137 | 0.03981 |
| ENSG00000101337 | TM9SF4 | 43.82163792 | 55.21907811 | 63.7040481 | 33.96093758 | 34.84293781 | 36.73013847 | 0.616492 | 5.50667 | 0.0014 | 0.04054 |
| ENSG00000149582 | TMEM25 | 10.72231566 | 9.108301545 | 8.10778794 | 3.396093758 | 4.479806289 | 4.081126497 | 1.225685 | 2.93385 | 0.00141 | 0.04061 |
| ENSG00000131238 | PPT1 | 20.97844368 | 21.63221617 | 21.42772527 | 5.821875013 | 11.94615011 | 15.30422436 | 0.9549 | 4.09314 | 0.00141 | 0.04061 |
| ENSG00000236801 | RPL24P8 | 80.18427364 | 52.37273388 | 80.49875169 | 113.5265628 | 107.5153509 | 98.45717674 | -0.581184 | 6.49223 | 0.00142 | 0.04072 |
| ENSG00000233597 | RP11-343B5.1 | 92.30515221 | 73.4356812 | 85.71090108 | 116.9226565 | 115.9772073 | 119.37295 | -0.482617 | 6.67018 | 0.00143 | 0.04085 |
| ENSG00000142173 | COL6A2 | 46.1525761 | 70.58933697 | 58.49189871 | 36.87187508 | 43.30479413 | 29.07802629 | 0.673516 | 5.58975 | 0.00145 | 0.04138 |
| ENSG00000132475 | H3F3B | 204.1901852 | 177.0426113 | 191.1121443 | 228.023438 | 305.6223402 | 265.2732223 | -0.479728 | 7.84397 | 0.00145 | 0.04138 |
| ENSG00000115760 | BIRC6 | 187.4074302 | 215.183624 | 163.8931419 | 263.4398443 | 304.1290714 | 234.1546328 | -0.500997 | 7.84071 | 0.00146 | 0.04145 |
| ENSG00000174748 | RPL15 | 45.22020083 | 50.66492734 | 54.43800474 | 63.55546889 | 73.17016939 | 86.21379725 | -0.571119 | 5.98405 | 0.00146 | 0.0415 |
| ENSG00000093010 | COMT | 15.85037967 | 15.9395277 | 22.00685298 | 8.247656269 | 3.982050035 | 12.7535203 | 1.098924 | 3.80523 | 0.00147 | 0.04166 |
| ENSG00000147853 | AK3 | 39.62594918 | 46.11077657 | 27.79813008 | 21.8320313 | 20.40800643 | 25.50704061 | 0.746936 | 4.96094 | 0.00148 | 0.04177 |
| ENSG00000197165 | SULT1A2 | 0.932375275 | 0.569268847 | 1.15825542 | 3.396093758 | 2.488781272 | 9.182534619 | -2.411525 | 2.00948 | 0.00149 | 0.04202 |

| | | | | | | | | | | | |
|---|---|---|---|---|---|---|---|---|---|---|---|
| ENSG00000147684 | NDUFB9 | 314.2104676 | 239.0929155 | 284.9308333 | 369.6890633 | 377.7969971 | 395.8692702 | -0.446174 | 8.37296 | 0.00152 | 0.04243 |
| ENSG00000169122 | FAM110B | 12.12087857 | 9.108301545 | 8.10778794 | 3.396093758 | 5.973075053 | 2.550704061 | 1.299298 | 2.98038 | 0.00152 | 0.04243 |
| ENSG00000130957 | FBP2 | 13.98562912 | 11.95464578 | 15.63644817 | 8.247656269 | 4.479806289 | 7.652112182 | 1.020713 | 3.49406 | 0.00152 | 0.04243 |
| ENSG00000238193 | RP11-555H23.: | 601.8482399 | 580.6542235 | 589.5520088 | 300.3117194 | 294.1739463 | 532.0768671 | 0.65378 | 8.9185 | 0.00154 | 0.04301 |
| ENSG00000121691 | CAT | 139.3901036 | 148.5791689 | 229.9137009 | 101.8828127 | 68.6903631 | 133.6568928 | 0.765927 | 7.10423 | 0.00155 | 0.04314 |
| ENSG00000099901 | RANBP1 | 16.31656731 | 15.9395277 | 14.47819275 | 26.68359381 | 27.87435025 | 22.95633655 | -0.724822 | 4.45056 | 0.00157 | 0.04347 |
| ENSG00000178996 | SNX18 | 13.98562912 | 22.77075386 | 16.21557588 | 7.762500018 | 10.45288134 | 8.672393807 | 0.963164 | 3.82331 | 0.00161 | 0.04439 |
| ENSG00000159314 | ARHGAP27 | 4.661876374 | 7.969763851 | 5.21214939 | 10.67343752 | 12.44390636 | 14.79408355 | -1.092904 | 3.37382 | 0.00161 | 0.04447 |
| ENSG00000231500 | RPS18 | 292.7658363 | 245.3548729 | 324.8906453 | 408.9867197 | 356.8912344 | 428.5182822 | -0.468633 | 8.42662 | 0.00162 | 0.04447 |
| ENSG00000184216 | IRAK1 | 7.925189836 | 9.677570391 | 12.74080962 | 4.851562511 | 2.488781272 | 5.101408121 | 1.258654 | 3.00285 | 0.00169 | 0.04624 |
| ENSG00000082146 | STRADB | 49.8820772 | 31.87905541 | 45.17196138 | 25.22812506 | 24.39005646 | 29.07802629 | 0.695853 | 5.14036 | 0.00169 | 0.04624 |
| ENSG00000074964 | ARHGEF10L | 72.2590838 | 63.18884197 | 67.75794207 | 42.6937501 | 34.84293781 | 56.11548934 | 0.607172 | 5.83446 | 0.00171 | 0.04653 |
| ENSG00000092439 | TRPM7 | 59.20582995 | 69.45079928 | 62.54579268 | 82.96171894 | 100.5467634 | 88.76450131 | -0.512015 | 6.2921 | 0.00171 | 0.04653 |
| ENSG00000174804 | FZD4 | 10.25612802 | 19.92440963 | 9.84517107 | 25.71328131 | 22.8967877 | 24.9968998 | -0.888379 | 4.32437 | 0.00172 | 0.04653 |
| ENSG00000238181 | AHCYP2 | 2.797125825 | 4.554150772 | 4.63302168 | 1.455468753 | 0.497756254 | 0.510140812 | 2.158903 | 1.74607 | 0.00172 | 0.04653 |
| ENSG00000237541 | HLA-DQA2 | 5.128064012 | 7.969763851 | 5.21214939 | 0 | 2.488781272 | 2.550704061 | 1.82849 | 2.26897 | 0.00176 | 0.04754 |
| ENSG00000146535 | GNA12 | 48.94970193 | 68.88153043 | 53.85887703 | 36.38671883 | 31.35864403 | 42.85182822 | 0.629831 | 5.57994 | 0.00176 | 0.04754 |
| ENSG00000166317 | SYNPO2L | 47.55113902 | 48.38785196 | 53.27974932 | 35.90156258 | 29.36761901 | 34.68957523 | 0.574481 | 5.40629 | 0.00177 | 0.04754 |
| ENSG00000125148 | MT2A | 0.466187637 | 1.138537693 | 6.37040481 | 28.13906256 | 2.488781272 | 14.79408355 | -2.518458 | 3.33248 | 0.00179 | 0.04798 |
| ENSG00000156298 | TSPAN7 | 44.28782556 | 33.58686195 | 37.06417344 | 53.85234387 | 54.25543173 | 64.78788314 | -0.579605 | 5.62019 | 0.0018 | 0.04822 |
| ENSG00000158125 | XDH | 37.29501099 | 34.72539964 | 35.32679031 | 65.4960939 | 35.83845032 | 97.43689512 | -0.885829 | 5.70648 | 0.0018 | 0.04822 |
| ENSG00000244734 | HBB | 193.4678695 | 401.9038057 | 854.7925 | 196.4882817 | 237.4297333 | 228.5430838 | 1.129364 | 8.45936 | 0.00181 | 0.04828 |
| ENSG00000065978 | YBX1 | 972.001224 | 879.5203679 | 871.0080758 | 1104.700784 | 1227.466923 | 1194.749782 | -0.373003 | 10.0262 | 0.00183 | 0.04873 |
| ENSG00000107954 | NEURL1 | 3.263313462 | 2.846344233 | 4.05389397 | 0.485156251 | 0.497756254 | 1.020281624 | 2.241317 | 1.58827 | 0.00184 | 0.04882 |
| ENSG00000204628 | RACK1 | 643.3389396 | 578.9464169 | 644.5691412 | 788.378908 | 782.4728319 | 846.8337482 | -0.372696 | 9.48233 | 0.00186 | 0.04914 |
| ENSG00000088682 | COQ9 | 324.000408 | 302.2817575 | 272.1900237 | 230.9343755 | 203.582308 | 234.1546328 | 0.427234 | 8.03432 | 0.00186 | 0.04914 |
| ENSG00000076555 | ACACB | 180.4146157 | 277.8031971 | 255.9744478 | 162.0421879 | 128.9188699 | 183.6506924 | 0.58748 | 7.63466 | 0.00187 | 0.04921 |
| ENSG00000196834 | POTEI | 59.20582995 | 38.14101272 | 60.80840955 | 39.29765634 | 31.35864403 | 26.52732223 | 0.702694 | 5.44407 | 0.00187 | 0.04929 |
| ENSG00000103381 | CPPED1 | 4.195688737 | 5.123419619 | 6.94953252 | 0.485156251 | 2.986537526 | 0.510140812 | 1.964836 | 2.10541 | 0.00191 | 0.04998 |
| ENSG00000100307 | CBX7 | 14.45181676 | 9.677570391 | 13.31993733 | 4.851562511 | 6.470831307 | 7.14197137 | 1.026294 | 3.36526 | 0.00191 | 0.04998 |
| ENSG00000186260 | MKL2 | 20.04606841 | 11.95464578 | 17.3738313 | 31.53515632 | 32.85191279 | 21.93605492 | -0.793411 | 4.57587 | 0.00191 | 0.04998 |

#### Suppl table 5 Cardiac DEs for HFD_Spp_24hrs vs HFD_CTL

| Ensembl ID | Gene Symbol | HFD_Spp_24hrs-1 | HFD_Spp_24hrs-2 | HFD_Spp_24hrs-3 | HFD-CTL-1 | HFD-CTL-2 | HFD-CTL-3 | logFC | logCPM | PValue | FDR |
|---|---|---|---|---|---|---|---|---|---|---|---|
| ENSG00000196296 | ATP2A1 | 764.0262141 | 713.8262261 | 707.1660134 | 366.395546 | 410.7073383 | 355.9094 | 0.949101 | 9.114289 | 4.69E-15 | 5.30E-11 |
| ENSG00000115593 | SMYD1 | 351.9928809 | 364.7573573 | 315.6889375 | 179.4302095 | 176.9399539 | 151.0239 | 1.025279 | 8.010551 | 3.09E-14 | 1.74E-10 |
| ENSG00000225024 | AC015977.5 | 33.91598071 | 20.17091377 | 33.05046638 | 92.77625007 | 71.03428807 | 112.4758 | -1.66077 | 5.94853 | 1.17E-13 | 4.38E-10 |
| ENSG00000133112 | TPT1 | 401.4918798 | 358.0337194 | 400.0246103 | 689.9350576 | 670.3053729 | 694.9211 | -0.82534 | 9.068156 | 1.65E-12 | 4.66E-09 |
| ENSG00000161533 | ACOX1 | 78.83173895 | 87.407293 | 86.04517971 | 40.97225257 | 40.03750782 | 31.68333 | 1.159741 | 5.952999 | 1.75E-11 | 3.94E-08 |
| ENSG00000136842 | TMOD1 | 175.0797923 | 209.5533819 | 198.3027983 | 101.2532678 | 113.0090947 | 97.16222 | 0.90541 | 7.229609 | 3.48E-10 | 6.55E-07 |
| ENSG00000181227 | DLSTP1 | 113.6643678 | 114.8621478 | 114.5369611 | 64.04857873 | 61.9935605 | 60.19833 | 0.879475 | 6.482051 | 2.39E-09 | 3.85E-06 |
| ENSG00000135218 | CD36 | 199.3709677 | 251.5761189 | 177.7887157 | 92.77625007 | 120.1125235 | 98.74638 | 1.016753 | 7.301688 | 3.13E-09 | 4.42E-06 |
| ENSG00000102858 | MGRN1 | 98.99799775 | 104.776691 | 100.860906 | 56.04250639 | 50.36976791 | 57.03 | 0.892127 | 6.307513 | 5.46E-09 | 6.85E-06 |
| ENSG00000181904 | C5orf24 | 20.1662588 | 17.92970113 | 11.39671254 | 36.26279825 | 45.84940412 | 51.22139 | -1.40706 | 4.972589 | 6.41E-09 | 7.20E-06 |
| ENSG00000119938 | PPP1R3C | 32.08268446 | 40.9021307 | 35.89964451 | 16.48309011 | 13.56109136 | 12.67333 | 1.330331 | 4.721354 | 7.01E-09 | 7.20E-06 |
| ENSG00000276293 | PIP4K2B | 21.99955505 | 23.53273273 | 30.77112387 | 8.477017773 | 6.457662552 | 9.505 | 1.61032 | 4.161151 | 1.29E-08 | 1.22E-05 |
| ENSG00000063660 | GPC1 | 323.5767889 | 284.0737022 | 389.1977334 | 157.2957742 | 169.1907589 | 206.4697 | 0.902933 | 8.000639 | 2.42E-08 | 1.98E-05 |
| ENSG00000104812 | GYS1 | 90.7481646 | 94.13093092 | 106.5592623 | 57.92628812 | 48.43246914 | 48.58111 | 0.902179 | 6.239566 | 2.58E-08 | 1.98E-05 |
| ENSG00000185641 | CTD-2287O16.1 | 199.8292917 | 145.1185185 | 180.6378938 | 298.5794038 | 304.8016725 | 400.2661 | -0.93142 | 7.999538 | 2.63E-08 | 1.98E-05 |
| ENSG00000130528 | HRC | 36.20760103 | 16.80909481 | 18.8045757 | 89.00868662 | 47.78670289 | 78.68027 | -1.57253 | 5.626099 | 3.43E-08 | 2.31E-05 |
| ENSG00000150401 | DCUN1D2 | 17.87463848 | 30.25637065 | 21.08391821 | 7.535126909 | 7.103428807 | 5.280555 | 1.776506 | 3.989915 | 3.48E-08 | 2.31E-05 |
| ENSG00000135821 | GLUL | 274.5361141 | 217.9579293 | 301.4430468 | 135.6322844 | 148.5262387 | 155.7764 | 0.852802 | 7.692272 | 3.89E-08 | 2.44E-05 |
| ENSG00000146729 | GBAS | 342.3680755 | 349.0688688 | 296.8843618 | 158.7086105 | 215.040163 | 128.8455 | 0.980023 | 7.96319 | 4.31E-08 | 2.52E-05 |
| ENSG00000162413 | KLHL21 | 48.12402668 | 33.05788645 | 57.55339835 | 19.77970814 | 16.14415638 | 18.48194 | 1.339289 | 5.062459 | 4.47E-08 | 2.52E-05 |
| ENSG00000105640 | RPL18A | 194.329403 | 144.5582153 | 223.9454015 | 356.0347465 | 332.5696214 | 346.4044 | -0.87885 | 8.062377 | 5.04E-08 | 2.71E-05 |
| ENSG00000236929 | RP11-699A7.1 | 88.45654428 | 54.34940654 | 74.07863154 | 114.9106854 | 188.5637465 | 196.4367 | -1.1971 | 6.908545 | 8.88E-08 | 4.56E-05 |
| ENSG00000162889 | MAPKAPK2 | 264.9113088 | 245.4127842 | 224.5152371 | 139.3998478 | 162.7330963 | 135.1822 | 0.753339 | 7.618596 | 9.50E-08 | 4.66E-05 |
| ENSG00000133135 | RNF128 | 53.16559138 | 64.99516659 | 62.11208336 | 24.96010789 | 31.64254651 | 32.73944 | 1.015378 | 5.521697 | 1.08E-07 | 5.08E-05 |
| ENSG00000119421 | NDUFA8 | 132.4556544 | 112.060632 | 130.4923586 | 197.3261359 | 205.3536692 | 220.1992 | -0.72987 | 7.386216 | 1.21E-07 | 5.47E-05 |
| ENSG00000264217 | RPL35AP25 | 1.374972191 | 0 | 1.709506882 | 6.593236046 | 8.394961318 | 10.03306 | -2.90719 | 2.506302 | 1.88E-07 | 8.16E-05 |
| ENSG00000235485 | RP1-101G11.2 | 285.5358917 | 317.1315887 | 308.2810743 | 199.2099177 | 149.8177712 | 191.6842 | 0.74724 | 7.926061 | 1.96E-07 | 8.17E-05 |
| ENSG00000163110 | PDLIM5 | 443.1993695 | 466.1722293 | 438.2035973 | 257.1362058 | 317.0712313 | 231.2883 | 0.744969 | 8.49131 | 2.04E-07 | 8.22E-05 |
| ENSG00000187446 | CHP1 | 32.99933258 | 40.34182754 | 35.89964451 | 19.77970814 | 14.20685761 | 12.67333 | 1.203396 | 4.762151 | 2.15E-07 | 8.36E-05 |
| ENSG00000091436 | MAP3K20 | 144.37208 | 133.3521521 | 166.9618388 | 251.4848606 | 224.0808906 | 275.1169 | -0.75756 | 7.645876 | 2.28E-07 | 8.56E-05 |
| ENSG00000167588 | GPD1 | 52.24894326 | 54.9097097 | 68.38027526 | 21.66348986 | 30.351014 | 32.21139 | 1.064651 | 5.468581 | 2.45E-07 | 8.93E-05 |
| ENSG00000213856 | VDAC1P2 | 106.7895068 | 117.1033605 | 103.7100841 | 44.26887059 | 68.45122305 | 62.31055 | 0.914722 | 6.405452 | 2.73E-07 | 9.26E-05 |
| ENSG00000125166 | GOT2 | 380.8672969 | 374.282511 | 377.8010208 | 258.5490421 | 248.6200083 | 233.9286 | 0.611958 | 8.292421 | 2.79E-07 | 9.26E-05 |
| ENSG00000168028 | RPSA | 37.58257322 | 24.09303589 | 37.03931577 | 66.87425132 | 66.51392429 | 62.31055 | -0.97897 | 5.646301 | 2.79E-07 | 9.26E-05 |
| ENSG00000198952 | SMG5 | 81.58168333 | 89.64850564 | 79.77698781 | 51.8039975 | 41.97480659 | 44.88472 | 0.84646 | 6.047906 | 2.90E-07 | 9.35E-05 |
| ENSG00000154518 | ATP5G3 | 4949.899887 | 3720.412984 | 4880.642147 | 6911.124212 | 7568.380511 | 7542.745 | -0.70046 | 12.53376 | 3.96E-07 | 0.00012 |
| ENSG00000004864 | SLC25A13 | 97.16470149 | 91.88971828 | 96.302221 | 51.33305207 | 59.41049548 | 54.91778 | 0.790205 | 6.253234 | 4.03E-07 | 0.00012 |
| ENSG00000159423 | ALDH4A1 | 59.58212827 | 68.35698555 | 53.56454896 | 33.90807109 | 30.351014 | 30.62722 | 0.929392 | 5.562089 | 4.52E-07 | 0.00013 |
| ENSG00000218682 | AC010150.1 | 123.2891731 | 76.20122979 | 98.01172788 | 169.06941 | 185.9806815 | 193.2683 | -0.87551 | 7.149593 | 4.57E-07 | 0.00013 |
| ENSG00000110717 | NDUFS8 | 153.5385613 | 128.3094237 | 173.2300307 | 252.4267515 | 244.7454107 | 252.4105 | -0.72014 | 7.656513 | 5.50E-07 | 0.00016 |
| ENSG00000106633 | GCK | 50.415647 | 39.78152438 | 49.57569957 | 20.721599 | 20.66452017 | 26.40278 | 1.042882 | 5.162214 | 6.12E-07 | 0.00017 |
| ENSG00000112695 | COX7A2 | 16.95799035 | 15.12818533 | 20.51408258 | 35.79185282 | 34.22561153 | 46.99694 | -1.15675 | 4.872089 | 6.24E-07 | 0.00017 |
| ENSG00000145494 | NDUFS6 | 118.2476084 | 107.0179036 | 111.6877829 | 168.5984646 | 178.8772527 | 192.7403 | -0.67836 | 7.201316 | 7.38E-07 | 0.00019 |
| ENSG00000224004 | ATP5C1P1 | 491.3233962 | 392.2122122 | 432.505241 | 675.3357493 | 665.7850091 | 693.865 | -0.6277 | 9.128169 | 7.74E-07 | 0.0002 |
| ENSG00000101558 | VAPA | 164.5383388 | 159.1260975 | 152.1461125 | 253.8395878 | 228.6012543 | 260.3314 | -0.64302 | 7.673778 | 7.86E-07 | 0.0002 |
| ENSG00000242058 | RPS4XP19 | 38.04089728 | 25.77394537 | 47.86619268 | 71.11276021 | 71.68005433 | 80.26444 | -0.99721 | 5.826822 | 8.09E-07 | 0.0002 |
| ENSG00000066926 | FECH | 73.79017425 | 88.52789932 | 79.77698781 | 44.73981602 | 51.01553416 | 31.15528 | 0.935873 | 5.965996 | 9.91E-07 | 0.00024 |
| ENSG00000182606 | TRAK1 | 170.9548757 | 213.4755041 | 182.3474007 | 123.3877031 | 109.1344971 | 112.4758 | 0.711575 | 7.258267 | 1.04E-06 | 0.00025 |
| ENSG00000169047 | IRS1 | 20.1662588 | 20.17091377 | 23.36326072 | 45.68170689 | 40.68327408 | 40.13222 | -0.9957 | 5.032163 | 1.24E-06 | 0.00029 |
| ENSG00000074416 | MGLL | 14.66637004 | 16.80909481 | 15.95539756 | 5.18039975 | 5.811896297 | 5.280555 | 1.534572 | 3.547723 | 1.30E-06 | 0.00029 |
| ENSG00000242706 | RPS27AP9 | 56.37385983 | 42.02273702 | 43.87734329 | 74.88032366 | 88.46997697 | 97.69027 | -0.86396 | 6.091572 | 1.39E-06 | 0.0003 |
| ENSG00000240371 | RPS4XP13 | 97.16470149 | 63.87456027 | 94.59271411 | 166.7146829 | 133.0278486 | 243.9617 | -1.08816 | 7.071096 | 1.42E-06 | 0.0003 |
| ENSG00000149596 | JPH2 | 373.0757878 | 331.1391677 | 352.7282532 | 248.1882426 | 220.8520593 | 224.4236 | 0.606958 | 8.194828 | 1.44E-06 | 0.0003 |
| ENSG00000100412 | ACO2 | 1433.637671 | 1660.738567 | 1513.483426 | 1063.865731 | 1053.244762 | 883.4369 | 0.618762 | 10.30964 | 1.45E-06 | 0.0003 |
| ENSG00000233055 | RP11-490D19.8 | 288.7441601 | 258.2997569 | 286.0574849 | 435.153579 | 408.1242733 | 405.0186 | -0.58322 | 8.442712 | 1.50E-06 | 0.00031 |
| ENSG00000251939 | RNU6-1278P | 0.916648127 | 1.12060632 | 2.849178136 | 6.593236046 | 9.040727573 | 11.08917 | -2.43741 | 2.637824 | 1.77E-06 | 0.00036 |
| ENSG00000115268 | RPS15 | 108.164479 | 91.32941512 | 126.5035092 | 157.7667197 | 200.1875391 | 215.4467 | -0.81272 | 7.235404 | 2.02E-06 | 0.0004 |
| ENSG00000217027 | TPT1P4 | 72.41520206 | 49.3066781 | 85.47534408 | 108.7883948 | 152.4008362 | 143.103 | -0.95963 | 6.68125 | 2.05E-06 | 0.0004 |
| ENSG00000084234 | APLP2 | 414.3249535 | 452.1646503 | 437.6337617 | 308.9402033 | 274.4506585 | 296.2392 | 0.565893 | 8.512284 | 2.86E-06 | 0.00054 |
| ENSG00000249617 | RP11-366M4.17 | 121.4558769 | 77.32183611 | 86.61501533 | 150.2315928 | 182.7518502 | 181.123 | -0.84069 | 7.068391 | 2.88E-06 | 0.00054 |
| ENSG00000185834 | RPL12P4 | 10.54145346 | 5.603031602 | 15.95539756 | 27.78578048 | 29.05948149 | 25.87472 | -1.3616 | 4.328102 | 3.07E-06 | 0.00057 |
| ENSG00000185088 | RPS27L | 4.583240636 | 5.042728442 | 5.128520645 | 11.7736358 | 14.85262387 | 19.53805 | -1.63241 | 3.474171 | 3.27E-06 | 0.0006 |
| ENSG00000112651 | MRPL2 | 78.37341488 | 66.11577291 | 67.81043964 | 107.8465039 | 115.5921597 | 124.0931 | -0.70507 | 6.559309 | 3.35E-06 | 0.0006 |
| ENSG00000100994 | PYGB | 622.8624025 | 592.2404404 | 589.7798741 | 442.2177605 | 402.312377 | 399.21 | 0.536619 | 8.992498 | 4.19E-06 | 0.00074 |
| ENSG00000062485 | CS | 515.1562475 | 528.3658801 | 495.18716 | 339.0807109 | 379.7105581 | 301.5197 | 0.594241 | 8.740037 | 4.79E-06 | 0.00082 |
| ENSG00000233383 | RP11-324F21.1 | 2.749944382 | 1.12060632 | 2.279342509 | 7.064181478 | 9.686493828 | 9.505 | -2.01314 | 2.676315 | 4.79E-06 | 0.00082 |
| ENSG00000198959 | TGM2 | 52.24894326 | 59.95243815 | 49.57569957 | 29.19861677 | 32.93407902 | 26.93083 | 0.865508 | 5.422274 | 4.84E-06 | 0.00082 |
| ENSG00000242299 | RP11-234A1.1 | 175.5381164 | 143.9979122 | 155.5651262 | 242.5368974 | 227.9554881 | 274.0608 | -0.64674 | 7.675193 | 5.08E-06 | 0.00084 |
| ENSG00000146085 | MUT | 116.4143122 | 136.1536679 | 111.6877829 | 67.34519675 | 69.74275556 | 83.43277 | 0.722478 | 6.623368 | 5.17E-06 | 0.00085 |
| ENSG00000197444 | OGDHL | 321.7434927 | 327.2170456 | 293.465348 | 211.4544989 | 224.7266568 | 181.123 | 0.612158 | 8.028367 | 5.73E-06 | 0.00092 |
| ENSG00000234031 | RPS3AP44 | 8.249833146 | 7.283941083 | 6.838027526 | 16.48309011 | 27.76794897 | 17.95389 | -1.43726 | 3.897852 | 5.85E-06 | 0.00093 |
| ENSG00000220069 | RPL7P27 | 10.0831294 | 3.922122122 | 9.117370035 | 25.43105332 | 18.7272214 | 19.01 | -1.43258 | 3.957513 | 6.04E-06 | 0.00095 |
| ENSG00000171867 | PRNP | 46.29073043 | 44.26394966 | 42.16783641 | 24.01821702 | 24.5391177 | 24.29055 | 0.867426 | 5.146605 | 6.58E-06 | 0.001 |

| | | | | | | | | | | | |
|---|---|---|---|---|---|---|---|---|---|---|---|
| ENSG00000136872 | ALDOB | 16.95799035 | 21.85182325 | 303.7223893 | 8.006072341 | 7.103428807 | 10.56111 | 3.724539 | 5.942701 | 6.63E-06 | 0.001 |
| ENSG00000112964 | GHR | 10.54145346 | 14.56788217 | 7.407863154 | 18.83781727 | 38.74597531 | 33.79555 | -1.47101 | 4.415358 | 6.66E-06 | 0.001 |
| ENSG00000183298 | RPSAP19 | 53.62391545 | 34.17849277 | 52.99471333 | 92.30530464 | 80.07501565 | 80.26444 | -0.8403 | 6.059534 | 7.03E-06 | 0.00104 |
| ENSG00000073905 | VDAC1P1 | 118.2476084 | 119.9048763 | 111.1179473 | 54.62967009 | 84.59537943 | 65.47889 | 0.782687 | 6.543916 | 7.17E-06 | 0.00105 |
| ENSG00000164587 | RPS14 | 1176.976195 | 834.8517088 | 1181.839091 | 1696.345445 | 1593.751118 | 1906.28 | -0.70214 | 10.45063 | 7.65E-06 | 0.00111 |
| ENSG00000141552 | ANAPC11 | 14.20804597 | 10.08545688 | 7.977698781 | 22.60538073 | 23.24758519 | 31.15528 | -1.2285 | 4.270238 | 8.03E-06 | 0.00115 |
| ENSG00000235907 | RP4-580O19.2 | 13.29139785 | 19.05030745 | 13.67605505 | 28.25672591 | 32.93407902 | 34.85167 | -1.06183 | 4.619397 | 8.22E-06 | 0.00116 |
| ENSG00000122359 | ANXA11 | 82.95665552 | 86.28668668 | 92.88320723 | 51.8039975 | 52.30706667 | 57.03 | 0.698917 | 6.16278 | 8.30E-06 | 0.00116 |
| ENSG00000219023 | RP3-340B19.2 | 14.20804597 | 10.64576004 | 12.5363838 | 23.54727159 | 27.12218272 | 31.68333 | -1.12128 | 4.388937 | 8.55E-06 | 0.00118 |
| ENSG00000196776 | CD47 | 10.54145346 | 6.723637923 | 14.24589068 | 21.19254443 | 30.351014 | 25.87472 | -1.28075 | 4.24776 | 9.05E-06 | 0.00122 |
| ENSG00000231006 | RPL7P32 | 77.91509082 | 43.14334334 | 67.81043964 | 105.0208313 | 114.9463934 | 126.7333 | -0.86865 | 6.496533 | 9.08E-06 | 0.00122 |
| ENSG00000147123 | NDUFB11 | 106.7895068 | 74.52032031 | 125.363838 | 181.3139913 | 162.0873301 | 192.2122 | -0.80499 | 7.144223 | 9.24E-06 | 0.00123 |
| ENSG00000135766 | EGLN1 | 87.99822022 | 91.32941512 | 78.63731655 | 49.92021578 | 59.41049548 | 44.88472 | 0.750353 | 6.1247 | 9.92E-06 | 0.0013 |
| ENSG00000102683 | SGCG | 72.87352612 | 74.52032031 | 66.67076838 | 43.32697973 | 45.20363787 | 23.7625 | 0.935082 | 5.794869 | 1.05E-05 | 0.00135 |
| ENSG00000119689 | DLST | 423.0331107 | 451.044044 | 368.6836508 | 284.4510408 | 282.1998535 | 272.4767 | 0.566803 | 8.443631 | 1.05E-05 | 0.00135 |
| ENSG00000159592 | GPBP1L1 | 54.99888764 | 48.18607178 | 41.02816516 | 76.29315996 | 78.78348314 | 89.24138 | -0.75264 | 6.040243 | 1.10E-05 | 0.00139 |
| ENSG00000140945 | CDH13 | 84.78995177 | 114.3018447 | 88.89435784 | 58.86817898 | 51.66130042 | 59.14222 | 0.755202 | 6.274099 | 1.11E-05 | 0.00139 |
| ENSG00000214389 | RPS3AP26 | 56.37385983 | 47.06546546 | 47.29635706 | 85.71206859 | 83.94961318 | 78.15222 | -0.71057 | 6.076291 | 1.16E-05 | 0.00144 |
| ENSG00000177981 | ASB8 | 51.33229513 | 41.46243386 | 46.72652143 | 21.66348986 | 29.05948149 | 24.81861 | 0.901928 | 5.208045 | 1.23E-05 | 0.0015 |
| ENSG00000185739 | SRL | 311.6603633 | 504.2728442 | 349.3092395 | 257.1362058 | 182.106084 | 206.9978 | 0.847013 | 8.24352 | 1.24E-05 | 0.0015 |
| ENSG00000004799 | PDK4 | 158.121802 | 193.3045903 | 160.1238112 | 126.6843212 | 80.7207819 | 83.96083 | 0.803675 | 7.078164 | 1.32E-05 | 0.00159 |
| ENSG00000242262 | RP11-100N21.1 | 282.3276232 | 246.5333905 | 268.3925804 | 362.1570371 | 388.1055194 | 431.4214 | -0.56639 | 8.369853 | 1.37E-05 | 0.00163 |
| ENSG00000213862 | CTD-2270N23.1 | 162.2467185 | 115.422451 | 154.425455 | 217.5767895 | 230.5385531 | 241.3214 | -0.67104 | 7.554038 | 1.40E-05 | 0.00164 |
| ENSG00000108107 | RPL28 | 370.3258434 | 265.583698 | 385.208884 | 553.3608824 | 501.1146141 | 587.7258 | -0.68529 | 8.797662 | 1.49E-05 | 0.00173 |
| ENSG00000005339 | CREBBP | 75.6234705 | 91.88971828 | 98.01172788 | 149.2897019 | 128.5074848 | 141.5189 | -0.6666 | 6.847273 | 1.55E-05 | 0.00178 |
| ENSG00000161547 | SRSF2 | 123.2891731 | 104.2163878 | 108.2687692 | 167.6565737 | 169.8365251 | 168.4497 | -0.58766 | 7.142843 | 1.79E-05 | 0.00201 |
| ENSG00000237550 | RPL9P9 | 99.91464587 | 73.39971399 | 105.419591 | 129.5099938 | 185.9806815 | 166.3375 | -0.78394 | 6.994102 | 1.80E-05 | 0.00201 |
| ENSG00000149577 | SIDT2 | 46.74905449 | 55.47001286 | 45.01701455 | 20.25065357 | 29.70524774 | 29.57111 | 0.899468 | 5.278245 | 1.81E-05 | 0.00201 |
| ENSG00000169253 | RP11-220D10.1 | 1.374972191 | 0.56030316 | 1.139671254 | 11.30269036 | 5.811896297 | 4.7525 | -2.7507 | 2.405878 | 1.83E-05 | 0.00201 |
| ENSG00000130255 | RPL36 | 132.9139785 | 78.44244243 | 135.6208793 | 167.6565737 | 247.3284758 | 241.3214 | -0.91544 | 7.391724 | 1.87E-05 | 0.00201 |
| ENSG00000225200 | AB019441.29 | 118.7059325 | 89.64850564 | 98.01172788 | 144.5802476 | 168.5449926 | 177.9547 | -0.67425 | 7.064828 | 1.87E-05 | 0.00201 |
| ENSG00000136859 | ANGPTL2 | 53.62391545 | 67.79668239 | 68.38027526 | 36.73374368 | 32.93407902 | 39.60416 | 0.784201 | 5.6699 | 1.88E-05 | 0.00201 |
| ENSG00000163382 | NAXE | 47.20737856 | 47.06546546 | 49.00586394 | 78.17694169 | 78.13771688 | 73.92777 | -0.68396 | 5.982506 | 1.89E-05 | 0.00201 |
| ENSG00000166016 | ABTB2 | 44.91575824 | 45.38455598 | 45.58685018 | 24.96010789 | 22.60181893 | 28.515 | 0.828552 | 5.189511 | 1.90E-05 | 0.00201 |
| ENSG00000173915 | USMG5 | 198.9126436 | 187.7015587 | 239.3309634 | 284.4510408 | 340.9645828 | 336.8994 | -0.61981 | 8.052562 | 1.94E-05 | 0.00203 |
| ENSG00000165410 | CFL2 | 400.5752316 | 384.3679679 | 344.7505545 | 546.7676464 | 507.5722766 | 581.3891 | -0.53376 | 8.851793 | 2.06E-05 | 0.00213 |
| ENSG00000109079 | TNFAIP1 | 29.33274007 | 34.73879593 | 31.3409595 | 15.54119925 | 15.49839013 | 17.95389 | 0.952869 | 4.652893 | 2.08E-05 | 0.00214 |
| ENSG00000116688 | MFN2 | 474.8237299 | 639.3059058 | 580.0926685 | 416.3157617 | 291.2405811 | 328.9786 | 0.705939 | 8.83391 | 2.12E-05 | 0.00216 |
| ENSG00000048740 | CELF2 | 67.37363736 | 61.63334763 | 45.01701455 | 87.12490489 | 97.51070454 | 115.1161 | -0.77677 | 6.321659 | 2.25E-05 | 0.00227 |
| ENSG00000128928 | IVD | 156.2885057 | 150.7215501 | 145.8779206 | 104.0789404 | 105.9056659 | 89.76944 | 0.597323 | 6.984056 | 2.28E-05 | 0.00227 |
| ENSG00000008988 | RPS20 | 86.16492397 | 70.59819819 | 70.08978215 | 103.607995 | 127.2159523 | 143.6311 | -0.71575 | 6.660546 | 2.41E-05 | 0.00238 |
| ENSG00000148985 | PGAP2 | 43.99911011 | 57.7112255 | 47.86619268 | 27.31483505 | 29.70524774 | 26.93083 | 0.831121 | 5.320009 | 2.49E-05 | 0.00244 |
| ENSG00000149257 | SERPINH1 | 240.1618093 | 258.2997569 | 202.8614833 | 162.0052286 | 158.2127325 | 136.2383 | 0.62008 | 7.601035 | 2.52E-05 | 0.00244 |
| ENSG00000083720 | OXCT1 | 357.0344456 | 368.1191763 | 275.8004436 | 218.9896258 | 229.2470206 | 202.7733 | 0.622133 | 8.11132 | 2.53E-05 | 0.00244 |
| ENSG00000095321 | CRAT | 340.0764552 | 321.614014 | 312.8397593 | 237.8274431 | 231.1843194 | 197.4928 | 0.548687 | 8.101713 | 2.62E-05 | 0.00249 |
| ENSG00000123684 | LPGAT1 | 44.45743417 | 40.9021307 | 50.14553519 | 69.22897848 | 73.6173531 | 79.20833 | -0.71179 | 5.9199 | 2.62E-05 | 0.00249 |
| ENSG00000159199 | ATP5G1 | 710.4022986 | 531.7276991 | 795.4905356 | 1073.755585 | 976.3985779 | 1242.515 | -0.69247 | 9.79681 | 2.72E-05 | 0.00256 |
| ENSG00000154582 | ELOC | 88.91486835 | 93.0103246 | 104.2799198 | 152.5863199 | 127.2159523 | 179.0108 | -0.68578 | 6.968881 | 2.84E-05 | 0.00265 |
| ENSG00000177600 | RPLP2 | 550.4472004 | 330.5788645 | 508.8632151 | 688.9931668 | 801.3959227 | 889.7736 | -0.77502 | 9.297599 | 2.98E-05 | 0.00273 |
| ENSG00000129116 | PALLD | 80.6650352 | 94.69123408 | 87.18485096 | 53.2168338 | 58.76472923 | 53.33361 | 0.667997 | 6.17666 | 2.98E-05 | 0.00273 |
| ENSG00000230849 | GOT2P2 | 95.7897293 | 118.2239668 | 103.7100841 | 56.51345182 | 68.45122305 | 72.34361 | 0.68885 | 6.44022 | 2.99E-05 | 0.00273 |
| ENSG00000176986 | SEC24C | 77.45676676 | 110.3797226 | 95.16254974 | 58.86817898 | 49.72400165 | 60.19833 | 0.734835 | 6.254934 | 3.19E-05 | 0.00288 |
| ENSG00000099795 | NDUFB7 | 156.2885057 | 110.3797226 | 168.10151 | 237.3564976 | 220.8520593 | 240.2653 | -0.68339 | 7.569442 | 3.22E-05 | 0.00289 |
| ENSG00000168209 | DDIT4 | 12.83307378 | 11.76636636 | 18.23474007 | 4.709454318 | 5.166130042 | 5.808611 | 1.434152 | 3.43488 | 3.28E-05 | 0.00291 |
| ENSG00000224631 | RPS27AP16 | 296.5356692 | 241.4906621 | 260.9847173 | 405.4840168 | 355.8172066 | 421.3883 | -0.56531 | 8.372719 | 3.31E-05 | 0.00292 |
| ENSG00000270672 | MTRNR2L6 | 622.4040784 | 759.7710853 | 887.2340715 | 1244.708776 | 1061.639724 | 1185.485 | -0.62277 | 9.90866 | 3.40E-05 | 0.00297 |
| ENSG00000184083 | FAM120C | 12.83307378 | 12.32666953 | 14.24589068 | 24.96010789 | 25.18488395 | 27.45889 | -0.97679 | 4.35778 | 3.54E-05 | 0.00307 |
| ENSG00000180211 | RP1-278E11.3 | 87.99822022 | 80.12335191 | 104.8497554 | 199.6808631 | 114.9463934 | 167.3936 | -0.82591 | 6.990937 | 3.81E-05 | 0.00326 |
| ENSG00000187079 | TEAD1 | 109.5394512 | 112.060632 | 82.62616594 | 136.5741752 | 175.0026552 | 191.6842 | -0.72155 | 7.081663 | 3.82E-05 | 0.00326 |
| ENSG00000213051 | RPL5P5 | 2.749944382 | 0 | 2.849178136 | 7.064181478 | 7.749195063 | 8.448889 | -1.97565 | 2.548504 | 3.87E-05 | 0.00329 |
| ENSG00000111481 | COPZ1 | 32.08268446 | 37.54031174 | 45.01701455 | 16.01214468 | 25.83065021 | 15.31361 | 1.019415 | 4.883392 | 4.20E-05 | 0.00354 |
| ENSG00000166123 | GPT2 | 17.41631442 | 7.844244243 | 17.66490444 | 3.767563455 | 3.874597531 | 5.808611 | 1.665188 | 3.404278 | 4.40E-05 | 0.00367 |
| ENSG00000104888 | SLC17A7 | 46.74905449 | 43.14334334 | 53.56454896 | 24.48916246 | 28.41371523 | 29.57111 | 0.801391 | 5.275252 | 4.64E-05 | 0.00385 |
| ENSG00000108960 | MMD | 19.70793474 | 27.45485485 | 21.65375383 | 10.83174493 | 10.33226008 | 11.08917 | 1.077649 | 4.162567 | 4.83E-05 | 0.00398 |
| ENSG00000243256 | RPL30P14 | 39.87419354 | 24.65333905 | 46.1566858 | 60.28101528 | 63.93085927 | 82.37666 | -0.89715 | 5.751684 | 4.90E-05 | 0.00399 |
| ENSG00000091542 | ALKBH5 | 29.33274007 | 36.41970542 | 45.58685018 | 21.66348986 | 18.08145515 | 16.36972 | 0.969814 | 4.854949 | 4.91E-05 | 0.00399 |
| ENSG00000100911 | PSME2 | 22.91620318 | 33.05788645 | 30.20128824 | 46.15265232 | 54.24436544 | 50.16528 | -0.8097 | 5.331804 | 4.98E-05 | 0.00401 |
| ENSG00000135390 | ATP5G2 | 469.7821652 | 348.5085657 | 491.1983106 | 612.7000068 | 660.6188791 | 784.6905 | -0.65171 | 9.135016 | 5.02E-05 | 0.00402 |
| ENSG00000234335 | RPS4XP11 | 0.916648127 | 1.12060632 | 1.139671254 | 5.651345182 | 5.811896297 | 5.280555 | -2.33751 | 2.11962 | 5.28E-05 | 0.0042 |
| ENSG00000156298 | TSPAN7 | 43.54078605 | 33.05788645 | 36.46948014 | 66.87425132 | 52.30706667 | 76.04 | -0.78691 | 5.71716 | 5.39E-05 | 0.00424 |
| ENSG00000233045 | AC097523.1 | 18.33296255 | 17.92970113 | 15.95539756 | 28.25672591 | 37.4544428 | 32.73944 | -0.89638 | 4.700573 | 5.41E-05 | 0.00424 |
| ENSG00000168079 | SCARA5 | 7.791509082 | 7.844244243 | 5.128520645 | 1.412836296 | 1.937298766 | 1.584167 | 2.063291 | 2.431854 | 5.44E-05 | 0.00424 |
| ENSG00000138449 | SLC40A1 | 26.58279569 | 21.29152009 | 29.63145261 | 14.12836296 | 12.9153251 | 11.08917 | 1.016842 | 4.351589 | 5.63E-05 | 0.00435 |
| ENSG00000100934 | SEC23A | 51.79061919 | 69.47759187 | 63.25175462 | 41.443198 | 34.87137778 | 31.68333 | 0.760184 | 5.638726 | 6.14E-05 | 0.00472 |
| ENSG00000125691 | RPL23 | 471.1571374 | 374.282511 | 439.3432686 | 576.4372086 | 615.4152412 | 813.7336 | -0.64187 | 9.101801 | 6.24E-05 | 0.00473 |

| | | | | | | | | | | | |
|---|---|---|---|---|---|---|---|---|---|---|---|
| ENSG00000132763 | MMACHC | 24.74949944 | 30.81667381 | 25.64260322 | 13.65741752 | 9.686493828 | 16.36972 | 1.005283 | 4.413256 | 6.25E-05 | 0.00473 |
| ENSG00000102309 | PIN4 | 29.33274007 | 14.56788217 | 22.79342509 | 42.8560343 | 39.39174157 | 45.94083 | -0.92756 | 5.071789 | 6.32E-05 | 0.00476 |
| ENSG00000263232 | ATP5A1P3 | 76.99844269 | 34.17849277 | 70.65961777 | 94.6600318 | 132.3820823 | 137.2944 | -0.99515 | 6.521684 | 6.42E-05 | 0.00477 |
| ENSG00000259330 | INAFM2 | 24.74949944 | 28.57546117 | 34.75997326 | 44.73981602 | 54.24436544 | 53.86166 | -0.79754 | 5.356891 | 6.43E-05 | 0.00477 |
| ENSG00000187642 | PERM1 | 20.1662588 | 25.77394537 | 21.65375383 | 13.18647209 | 8.394961318 | 9.505 | 1.085275 | 4.136903 | 6.48E-05 | 0.00478 |
| ENSG00000138867 | GUCD1 | 15.1246941 | 13.44727585 | 11.96654817 | 6.593236046 | 5.166130042 | 4.224444 | 1.32672 | 3.40507 | 6.66E-05 | 0.00488 |
| ENSG00000136830 | FAM129B | 27.95776788 | 28.01515801 | 25.0727676 | 13.65741752 | 14.85262387 | 14.2575 | 0.92658 | 4.443444 | 6.78E-05 | 0.00494 |
| ENSG00000175931 | UBE2O | 44.45743417 | 53.78910338 | 44.44717892 | 27.31483505 | 20.01875391 | 32.21139 | 0.82532 | 5.256847 | 6.93E-05 | 0.00501 |
| ENSG00000166317 | SYNPO2L | 46.74905449 | 47.62576862 | 52.4248777 | 28.72767134 | 32.28831276 | 26.40278 | 0.751056 | 5.324875 | 7.13E-05 | 0.00513 |
| ENSG00000239344 | RP11-771F20.1 | 6.416536891 | 2.801515801 | 3.419013763 | 14.59930839 | 9.040727573 | 14.2575 | -1.54892 | 3.265122 | 7.44E-05 | 0.00531 |
| ENSG00000227008 | RP3-417G15.1 | 46.74905449 | 25.77394537 | 39.8884939 | 62.164797 | 73.6173531 | 63.36666 | -0.81006 | 5.725119 | 7.47E-05 | 0.00531 |
| ENSG00000078699 | CBFA2T2 | 8.708157209 | 9.525153724 | 7.407863154 | 19.30876271 | 17.43568889 | 18.48194 | -1.1041 | 3.861049 | 7.72E-05 | 0.00542 |
| ENSG00000196821 | C6orf106 | 109.0811271 | 102.5354783 | 113.9671254 | 64.99046959 | 62.63932676 | 82.37666 | 0.629595 | 6.499 | 7.73E-05 | 0.00542 |
| ENSG00000011198 | ABHD5 | 35.74927696 | 38.66091806 | 33.620302 | 16.01214468 | 20.66452017 | 22.70639 | 0.871124 | 4.856795 | 7.84E-05 | 0.00546 |
| ENSG00000154122 | ANKH | 143.913756 | 133.9124553 | 131.6320299 | 90.89246835 | 83.30384692 | 101.3867 | 0.569231 | 6.850768 | 8.41E-05 | 0.00582 |
| ENSG00000129559 | NEDD8 | 48.12402668 | 32.49758329 | 35.89964451 | 90.89246835 | 50.36976791 | 76.56805 | -0.90317 | 5.837425 | 8.65E-05 | 0.00594 |
| ENSG00000112992 | NNT | 271.7861697 | 267.2646074 | 250.727676 | 181.7849367 | 202.1248379 | 149.9678 | 0.567743 | 7.792464 | 8.68E-05 | 0.00594 |
| ENSG00000119715 | ESRRB | 15.1246941 | 15.68848849 | 19.94424695 | 7.064181478 | 3.874597531 | 9.505 | 1.26537 | 3.700731 | 8.77E-05 | 0.00596 |
| ENSG00000147862 | NFIB | 396.9086391 | 358.5940226 | 353.2980889 | 512.8595753 | 483.0331589 | 558.1547 | -0.48665 | 8.797405 | 8.99E-05 | 0.00605 |
| ENSG00000111667 | USP5 | 159.4967741 | 155.2039754 | 120.2353173 | 102.1951587 | 96.21917203 | 77.09611 | 0.660041 | 6.902941 | 9.05E-05 | 0.00605 |
| ENSG00000167315 | ACAA2 | 118.2476084 | 75.64092663 | 100.2910704 | 150.7025382 | 168.5449926 | 149.4397 | -0.66511 | 7.001547 | 9.06E-05 | 0.00605 |
| ENSG00000256356 | HSPA8P5 | 223.6621431 | 207.3121693 | 185.7664145 | 128.5681029 | 157.5669663 | 128.3175 | 0.579141 | 7.434446 | 9.29E-05 | 0.00617 |
| ENSG00000022267 | FHL1 | 93.95643305 | 120.4651795 | 109.4084404 | 63.10668787 | 80.7207819 | 59.14222 | 0.678399 | 6.47079 | 9.42E-05 | 0.0062 |
| ENSG00000136942 | RPL35 | 127.8724138 | 133.3521521 | 148.1572631 | 190.2619545 | 189.2095128 | 213.8625 | -0.53753 | 7.393038 | 9.45E-05 | 0.0062 |
| ENSG00000065154 | OAT | 96.24805337 | 82.92486772 | 94.59271411 | 58.86817898 | 67.15969054 | 45.41278 | 0.6823 | 6.234504 | 0.0001 | 0.00668 |
| ENSG00000119446 | RBM18 | 23.37452725 | 22.97242957 | 21.08391821 | 12.24458123 | 12.9153251 | 5.280555 | 1.153935 | 4.122927 | 0.0001 | 0.00669 |
| ENSG00000175920 | DOK7 | 13.29139785 | 11.76636636 | 10.25704129 | 3.296618023 | 5.166130042 | 4.7525 | 1.441954 | 3.201619 | 0.00011 | 0.0069 |
| ENSG00000103249 | CLCN7 | 12.83307378 | 15.68848849 | 15.38556193 | 5.18039975 | 6.457662552 | 6.864722 | 1.240373 | 3.516374 | 0.00011 | 0.0069 |
| ENSG00000231096 | NDUFB4P3 | 9.624805337 | 12.32666953 | 21.65375383 | 21.19254443 | 41.97480659 | 43.82861 | -1.29716 | 4.684745 | 0.00011 | 0.00692 |
| ENSG00000197879 | MYO1C | 86.16492397 | 107.5782068 | 102.0005773 | 64.99046959 | 58.76472923 | 68.11916 | 0.616025 | 6.363707 | 0.00011 | 0.00699 |
| ENSG00000233762 | AC007969.5 | 354.2845012 | 254.3776347 | 298.024033 | 492.1379763 | 399.0835457 | 481.5866 | -0.59792 | 8.574401 | 0.00011 | 0.00699 |
| ENSG00000197226 | TBC1D9B | 76.54011863 | 77.32183611 | 82.62616594 | 52.74588837 | 34.22561153 | 55.97389 | 0.710811 | 6.011877 | 0.00011 | 0.00699 |
| ENSG00000108947 | EFNB3 | 142.0804597 | 115.422451 | 132.2018655 | 73.9384328 | 97.51070454 | 82.90472 | 0.624392 | 6.760627 | 0.00011 | 0.00699 |
| ENSG00000109861 | CTSC | 8.249833146 | 9.525153724 | 7.977698781 | 2.825672591 | 3.874597531 | 1.584167 | 1.642729 | 2.751526 | 0.00011 | 0.00699 |
| ENSG00000215464 | AP000354.2 | 11.45810159 | 14.56788217 | 21.65375383 | 27.78578048 | 36.16291029 | 31.15528 | -1.00232 | 4.619599 | 0.00011 | 0.00699 |
| ENSG00000162909 | CAPN2 | 44.91575824 | 50.98758758 | 48.43602831 | 29.19861677 | 32.28831276 | 25.34667 | 0.735176 | 5.306115 | 0.00011 | 0.00699 |
| ENSG00000113578 | FGF1 | 22.45787912 | 42.58304018 | 29.06161699 | 16.95403555 | 9.686493828 | 15.31361 | 1.131921 | 4.570489 | 0.00011 | 0.00699 |
| ENSG00000228224 | NACAP1 | 48.12402668 | 44.26394966 | 50.71537082 | 65.93236046 | 76.20041812 | 83.43277 | -0.65254 | 5.962829 | 0.00012 | 0.00725 |
| ENSG00000120992 | LYPLA1 | 21.08290693 | 18.49000429 | 10.25704129 | 7.535126909 | 6.457662552 | 5.280555 | 1.368271 | 3.674192 | 0.00012 | 0.00725 |
| ENSG00000169504 | CLIC4 | 38.04089728 | 50.98758758 | 34.19013763 | 24.96010789 | 21.95605268 | 20.06611 | 0.870388 | 5.036627 | 0.00012 | 0.00725 |
| ENSG00000099875 | MKNK2 | 116.4143122 | 99.73396252 | 147.0175918 | 70.64181478 | 64.57662552 | 87.12916 | 0.703293 | 6.625676 | 0.00012 | 0.00725 |
| ENSG00000144744 | UBA3 | 74.24849831 | 75.08062347 | 62.11208336 | 44.73981602 | 45.20363787 | 44.88472 | 0.65127 | 5.8799 | 0.00012 | 0.00725 |
| ENSG00000116497 | S100PBP | 2.291620318 | 1.680909481 | 1.709506882 | 8.947963205 | 7.103428807 | 5.808611 | -1.89986 | 2.502278 | 0.00012 | 0.00725 |
| ENSG00000137094 | DNAJB5 | 141.1638116 | 121.5857858 | 101.4307416 | 85.24112316 | 56.82743046 | 79.73639 | 0.709287 | 6.631724 | 0.00012 | 0.00725 |
| ENSG00000162852 | CNST | 24.74949944 | 20.73121693 | 21.08391821 | 36.26279825 | 40.68327408 | 38.02 | -0.77369 | 4.965475 | 0.00013 | 0.00751 |
| ENSG00000266563 | EIF1P5 | 46.74905449 | 26.33424853 | 58.6930696 | 64.51952416 | 87.82421071 | 109.3075 | -0.98569 | 6.053249 | 0.00013 | 0.00759 |
| ENSG00000172115 | CYCS | 96.70637743 | 80.12335191 | 119.0956461 | 155.4119925 | 147.2347062 | 151.5519 | -0.61957 | 6.977497 | 0.00013 | 0.00764 |
| ENSG00000175445 | LPL | 771.8177232 | 1028.716602 | 858.1724546 | 669.6844041 | 438.4752873 | 577.6928 | 0.655378 | 9.502585 | 0.00014 | 0.00781 |
| ENSG00000173599 | PC | 57.29050796 | 44.82425282 | 83.19600157 | 36.26279825 | 33.57984527 | 31.15528 | 0.870099 | 5.608982 | 0.00014 | 0.00781 |
| ENSG00000213261 | EEF1B2P6 | 48.58235075 | 41.46243386 | 43.30750767 | 67.34519675 | 75.55465186 | 66.535 | -0.64147 | 5.860232 | 0.00014 | 0.00782 |
| ENSG00000119471 | HSDL2 | 127.4140897 | 152.4024596 | 123.0844955 | 83.82828687 | 102.6768346 | 72.34361 | 0.643117 | 6.797886 | 0.00014 | 0.00782 |
| ENSG00000157870 | FAM213B | 0.458324064 | 1.12060632 | 2.279342509 | 4.709454318 | 5.811896297 | 7.920833 | -2.25655 | 2.234921 | 0.00014 | 0.00791 |
| ENSG00000174996 | KLC2 | 39.41586947 | 36.41970542 | 41.59800079 | 24.96010789 | 21.95605268 | 21.65028 | 0.768544 | 5.007102 | 0.00014 | 0.00797 |
| ENSG00000147853 | AK3 | 38.95754541 | 45.38455598 | 27.35211011 | 16.95403555 | 21.31028642 | 20.59417 | 0.933449 | 4.884857 | 0.00014 | 0.00802 |
| ENSG00000132589 | FLOT2 | 58.66548015 | 64.43486343 | 63.82159025 | 33.90807109 | 44.55787161 | 39.07611 | 0.677307 | 5.692817 | 0.00015 | 0.00811 |
| ENSG00000169567 | HINT1 | 106.7895068 | 115.422451 | 118.5258105 | 146.4640293 | 176.9399539 | 174.2583 | -0.54506 | 7.134944 | 0.00015 | 0.00813 |
| ENSG00000170417 | TMEM182 | 21.08290693 | 21.85182325 | 27.35211011 | 15.07025382 | 9.040727573 | 9.505 | 1.027977 | 4.20716 | 0.00015 | 0.00835 |
| ENSG00000111897 | SERINC1 | 149.4136447 | 196.6664092 | 143.5985781 | 90.42152291 | 125.9244198 | 81.84861 | 0.72083 | 7.046576 | 0.00015 | 0.00835 |
| ENSG00000213598 | RP11-112J1.1 | 23.83285131 | 8.404547404 | 13.10621943 | 35.79185282 | 27.76794897 | 36.96389 | -1.1252 | 4.677188 | 0.00016 | 0.00861 |
| ENSG00000183873 | SCN5A | 293.3274007 | 531.1673959 | 592.0592167 | 283.0382045 | 198.2502404 | 287.2622 | 0.879345 | 8.512241 | 0.00016 | 0.00861 |
| ENSG00000123505 | AMD1 | 128.7890619 | 161.9276133 | 105.9894267 | 76.76410539 | 94.92763952 | 73.92777 | 0.696144 | 6.755958 | 0.00016 | 0.0087 |
| ENSG00000235587 | GAPDHP65 | 556.8637373 | 563.104676 | 565.2769422 | 434.2116882 | 415.2277021 | 405.5466 | 0.424757 | 8.940119 | 0.00016 | 0.0087 |
| ENSG00000137872 | SEMA6D | 26.12447163 | 31.37697697 | 20.51408258 | 39.08847084 | 46.49517038 | 51.74944 | -0.80919 | 5.202717 | 0.00017 | 0.00889 |
| ENSG00000134531 | EMP1 | 18.33296255 | 21.29152009 | 9.117370035 | 40.03036171 | 30.351014 | 29.57111 | -1.03172 | 4.697371 | 0.00017 | 0.00889 |
| ENSG00000213939 | RP11-314A20.1 | 11.45810159 | 7.844244243 | 9.687205662 | 18.83781727 | 23.24758519 | 18.48194 | -1.03223 | 3.990615 | 0.00017 | 0.00894 |
| ENSG00000129991 | TNNI3 | 4151.957693 | 3172.436493 | 3769.462674 | 5604.721584 | 4750.256573 | 5475.408 | -0.51291 | 12.13205 | 0.00017 | 0.00894 |
| ENSG00000123562 | MORF4L2 | 83.41497958 | 68.91728871 | 80.91665906 | 109.2593402 | 116.8836922 | 118.8125 | -0.56015 | 6.605256 | 0.00017 | 0.00894 |
| ENSG00000107331 | ABCA2 | 66.45698923 | 73.96001715 | 74.07863154 | 51.8039975 | 43.91210536 | 41.18833 | 0.638563 | 5.899628 | 0.00017 | 0.00894 |
| ENSG00000164754 | RAD21 | 38.04089728 | 44.82425282 | 33.05046638 | 57.45534268 | 69.09698931 | 60.19833 | -0.68116 | 5.681427 | 0.00017 | 0.00894 |
| ENSG00000149806 | FAU | 159.4967741 | 111.5003289 | 173.7998663 | 195.9132996 | 252.4946058 | 275.1169 | -0.69951 | 7.611447 | 0.00017 | 0.00894 |
| ENSG00000243404 | RPL35AP32 | 45.83240636 | 33.05788645 | 34.19013763 | 51.8039975 | 64.57662552 | 79.20833 | -0.77766 | 5.712783 | 0.00017 | 0.00894 |
| ENSG00000107679 | PLEKHA1 | 27.49944382 | 30.25637065 | 24.50293197 | 13.18647209 | 14.20685761 | 16.89778 | 0.892773 | 4.473054 | 0.00017 | 0.00894 |
| ENSG00000090316 | MAEA | 60.4987764 | 57.7112255 | 50.71537082 | 32.02428937 | 42.62057284 | 25.34667 | 0.772613 | 5.519807 | 0.00018 | 0.00894 |
| ENSG00000161011 | SQSTM1 | 235.1202446 | 217.9579293 | 169.8110169 | 130.9228301 | 152.4008362 | 129.3736 | 0.598513 | 7.441254 | 0.00018 | 0.00933 |
| ENSG00000188846 | RPL14 | 183.3296255 | 158.5657943 | 188.045757 | 230.7632616 | 277.6794897 | 252.4105 | -0.5187 | 7.754789 | 0.00018 | 0.00933 |

| | | | | | | | | | | | |
|---|---|---|---|---|---|---|---|---|---|---|---|
| ENSG00000153179 | RASSF3 | 38.49922135 | 49.3066781 | 41.02816516 | 27.31483505 | 25.18488395 | 22.70639 | 0.767534 | 5.132814 | 0.00019 | 0.00941 |
| ENSG00000248448 | COX5BP1 | 558.6970336 | 464.4913198 | 663.2886701 | 860.8882494 | 781.3771688 | 823.7666 | -0.54858 | 9.436975 | 0.00019 | 0.00942 |
| ENSG00000114023 | FAM162A | 59.58212827 | 45.94485914 | 55.84389147 | 81.00261428 | 79.42924939 | 87.65722 | -0.61623 | 6.115113 | 0.00019 | 0.00942 |
| ENSG00000231586 | RPS23P9 | 1.374972191 | 0.56030316 | 1.709506882 | 7.064181478 | 3.228831276 | 8.976944 | -2.36164 | 2.307712 | 0.00019 | 0.00942 |
| ENSG00000175334 | BANF1 | 2.749944382 | 2.241212641 | 4.558685018 | 9.418908637 | 10.97802634 | 7.920833 | -1.55188 | 2.864727 | 0.00019 | 0.00951 |
| ENSG00000213923 | CSNK1E | 0.458324064 | 2.801515801 | 2.279342509 | 4.709454318 | 10.33226008 | 8.448889 | -2.07917 | 2.512581 | 0.00019 | 0.00955 |
| ENSG00000230807 | AC099535.4 | 1.833296255 | 2.801515801 | 3.419013763 | 8.006072341 | 7.103428807 | 10.03306 | -1.6581 | 2.714522 | 0.00019 | 0.00955 |
| ENSG00000255262 | ELOBP2 | 1.833296255 | 2.241212641 | 0.569835627 | 6.593236046 | 4.520363787 | 8.976944 | -2.06656 | 2.384456 | 0.0002 | 0.00955 |
| ENSG00000133639 | BTG1 | 70.5819058 | 64.99516659 | 80.34682343 | 96.0728681 | 121.404056 | 110.3636 | -0.59803 | 6.51469 | 0.0002 | 0.00955 |
| ENSG00000198795 | ZNF521 | 19.24961067 | 28.01515801 | 21.08391821 | 39.08847084 | 39.39174157 | 39.07611 | -0.78955 | 4.996745 | 0.0002 | 0.00963 |
| ENSG00000134201 | GSTM5 | 119.1642565 | 108.1385099 | 153.8556193 | 86.18301403 | 80.07501565 | 81.32055 | 0.618749 | 6.726466 | 0.0002 | 0.00972 |
| ENSG00000248610 | HSPA8P4 | 76.08179457 | 93.57062776 | 83.7658372 | 64.51952416 | 48.43246914 | 46.46889 | 0.656839 | 6.128763 | 0.0002 | 0.00975 |
| ENSG00000104388 | RAB2A | 159.0384501 | 152.4024596 | 152.1461125 | 195.9132996 | 249.9115408 | 221.2553 | -0.52079 | 7.564174 | 0.0002 | 0.00975 |
| ENSG00000240156 | COX6CP6 | 243.3700778 | 189.3824682 | 205.1408258 | 285.3929317 | 317.7169976 | 309.9686 | -0.51384 | 8.019575 | 0.00021 | 0.00977 |
| ENSG00000129521 | EGLN3 | 72.41520206 | 79.56304875 | 88.89435784 | 28.25672591 | 65.22239178 | 32.73944 | 0.9493 | 5.952144 | 0.00021 | 0.00977 |
| ENSG00000106244 | PDAP1 | 42.16581386 | 34.73879593 | 35.89964451 | 59.33912441 | 56.1816642 | 61.7825 | -0.64686 | 5.627829 | 0.00021 | 0.00978 |
| ENSG00000072954 | TMEM38A | 148.4969966 | 159.6864007 | 148.1572631 | 88.06679575 | 124.6328873 | 76.04 | 0.667785 | 6.966839 | 0.00021 | 0.00978 |
| ENSG00000273654 | CTB-52I2.4 | 247.9533184 | 245.4127842 | 202.8614833 | 165.3018466 | 172.4195901 | 139.4067 | 0.547499 | 7.620093 | 0.00021 | 0.00984 |
| ENSG00000178573 | MAF | 15.58301816 | 15.68848849 | 15.38556193 | 22.60538073 | 30.351014 | 38.02 | -0.95388 | 4.576431 | 0.00021 | 0.00984 |
| ENSG00000011114 | BTBD7 | 42.16581386 | 39.78152438 | 33.620302 | 63.5776333 | 60.05626174 | 58.08611 | -0.64799 | 5.661723 | 0.00021 | 0.00985 |
| ENSG00000218363 | SLC25A20P1 | 8.249833146 | 7.283941083 | 7.407863154 | 1.412836296 | 3.874597531 | 1.584167 | 1.796402 | 2.591118 | 0.00021 | 0.00985 |
| ENSG00000160803 | UBQLN4 | 38.04089728 | 45.94485914 | 38.74882265 | 23.07632616 | 18.08145515 | 28.515 | 0.800021 | 5.054556 | 0.00022 | 0.00985 |
| ENSG00000100234 | TIMP3 | 54.08223951 | 41.46243386 | 59.83274086 | 72.05465107 | 82.01231441 | 95.05 | -0.67749 | 6.095242 | 0.00022 | 0.00985 |
| ENSG00000143420 | ENSA | 50.87397106 | 38.66091806 | 36.46948014 | 29.66956221 | 20.66452017 | 18.48194 | 0.865231 | 5.080738 | 0.00022 | 0.00985 |
| ENSG00000174437 | ATP2A2 | 16039.05061 | 17455.68465 | 15733.7315 | 12355.72435 | 12590.50468 | 12658.02 | 0.388595 | 13.8211 | 0.00022 | 0.01008 |
| ENSG00000178035 | IMPDH2 | 60.95710046 | 50.42728442 | 59.83274086 | 96.54381353 | 83.30384692 | 80.7925 | -0.60645 | 6.191623 | 0.00022 | 0.01014 |
| ENSG00000213585 | VDAC1 | 319.9101964 | 345.7070499 | 304.2922249 | 169.5403555 | 258.3065021 | 214.9186 | 0.597608 | 8.075172 | 0.00022 | 0.01014 |
| ENSG00000131165 | CHMP1A | 40.3325176 | 33.61818961 | 34.75997326 | 21.66348986 | 16.14415638 | 23.7625 | 0.808906 | 4.890333 | 0.00023 | 0.01016 |
| ENSG00000146535 | GNA12 | 48.12402668 | 67.79668239 | 52.99471333 | 38.14657998 | 32.28831276 | 31.15528 | 0.721484 | 5.52879 | 0.00023 | 0.01032 |
| ENSG00000223904 | HMGB1P12 | 0 | 0 | 1.139671254 | 1.883781727 | 4.520363787 | 6.864722 | -3.40546 | 1.755685 | 0.00023 | 0.01032 |
| ENSG00000248522 | SBF1P1 | 2.291620318 | 0.56030316 | 1.139671254 | 4.238508887 | 5.166130042 | 10.56111 | -2.21216 | 2.336662 | 0.00025 | 0.01109 |
| ENSG00000232573 | RPL3P4 | 65.5403411 | 44.26394966 | 67.81043964 | 101.2532678 | 79.42924939 | 107.1953 | -0.69788 | 6.300062 | 0.00025 | 0.01114 |
| ENSG00000198612 | COPS8 | 53.62391545 | 53.22880022 | 45.58685018 | 27.31483505 | 32.28831276 | 34.85167 | 0.698039 | 5.401748 | 0.00025 | 0.01114 |
| ENSG00000215030 | RPL13P12 | 117.3309603 | 88.52789932 | 118.5258105 | 150.2315928 | 154.338135 | 184.8194 | -0.59142 | 7.094568 | 0.00025 | 0.01114 |
| ENSG00000227615 | RP11-864N7.2 | 22.45787912 | 20.17091377 | 20.51408258 | 39.08847084 | 31.64254651 | 36.96389 | -0.77287 | 4.887752 | 0.00025 | 0.01114 |
| ENSG00000214022 | REPIN1 | 9.624805337 | 13.44727585 | 8.547534408 | 5.18039975 | 3.874597531 | 1.584167 | 1.535216 | 3.025473 | 0.00026 | 0.01133 |
| ENSG00000232888 | RPS11P5 | 25.2078235 | 28.57546117 | 33.05046638 | 36.73374368 | 55.53589795 | 64.95083 | -0.85425 | 5.372918 | 0.00026 | 0.01133 |
| ENSG00000249921 | CTC-329H14.1 | 18.33296255 | 18.49000429 | 14.81572631 | 32.02428937 | 29.70524774 | 28.515 | -0.80053 | 4.626916 | 0.00026 | 0.01133 |
| ENSG00000213442 | RPL18AP3 | 18.79128661 | 7.844244243 | 17.66490444 | 22.60538073 | 35.51714404 | 39.60416 | -1.11954 | 4.618958 | 0.00027 | 0.01143 |
| ENSG00000132681 | ATP1A4 | 81.58168333 | 98.6133562 | 89.46419347 | 72.99654194 | 33.57984527 | 42.7725 | 0.835178 | 6.15389 | 0.00027 | 0.01156 |
| ENSG00000151148 | UBE3B | 54.54056357 | 47.06546546 | 41.02816516 | 32.4952348 | 28.41371523 | 23.7625 | 0.7531 | 5.291007 | 0.00027 | 0.01156 |
| ENSG00000152137 | HSPB8 | 334.5765665 | 280.7118833 | 276.3702792 | 216.6348986 | 216.9774618 | 203.3014 | 0.487262 | 8.000114 | 0.00027 | 0.01156 |
| ENSG00000229758 | DYNLT3P2 | 1.374972191 | 2.801515801 | 3.419013763 | 0 | 0 | 0 | 5.252652 | 1.205293 | 0.00027 | 0.01156 |
| ENSG00000151623 | NR3C2 | 21.54123099 | 20.73121693 | 21.08391821 | 44.73981602 | 35.51714404 | 30.62722 | -0.81163 | 4.913941 | 0.00027 | 0.01156 |
| ENSG00000185164 | NOMO2 | 3.666592509 | 9.525153724 | 11.96654817 | 0.941890864 | 1.29153251 | 3.168333 | 2.156942 | 2.597231 | 0.00028 | 0.0116 |
| ENSG00000165312 | OTUD1 | 28.87441601 | 20.73121693 | 18.23474007 | 13.65741752 | 9.686493828 | 8.976944 | 1.062291 | 4.17052 | 0.00028 | 0.0116 |
| ENSG00000121691 | CAT | 137.038895 | 146.2391248 | 226.224744 | 105.9627222 | 108.4887309 | 99.8025 | 0.694788 | 7.110786 | 0.00028 | 0.01179 |
| ENSG00000123131 | PRDX4 | 14.20804597 | 15.68848849 | 12.5363838 | 24.96010789 | 22.60181893 | 32.21139 | -0.91267 | 4.422113 | 0.00029 | 0.01205 |
| ENSG00000197852 | FAM212B | 8.249833146 | 8.404547404 | 5.128520645 | 2.354727159 | 1.937298766 | 2.112222 | 1.734088 | 2.544422 | 0.00029 | 0.01205 |
| ENSG00000196576 | PLXNB2 | 94.41475711 | 93.57062776 | 90.60386472 | 59.81006984 | 63.28509301 | 67.59111 | 0.548487 | 6.309969 | 0.00029 | 0.01205 |
| ENSG00000226144 | RPS27AP3 | 13.74972191 | 7.844244243 | 17.66490444 | 23.54727159 | 23.89335144 | 34.32361 | -1.05533 | 4.405989 | 0.00029 | 0.01212 |
| ENSG00000168283 | BMI1 | 32.54100852 | 32.49758329 | 30.20128824 | 51.33305207 | 49.0782354 | 49.63722 | -0.65463 | 5.38961 | 0.0003 | 0.01229 |
| ENSG00000145244 | CORIN | 57.74883202 | 63.87456027 | 62.11208336 | 37.20468912 | 39.39174157 | 42.7725 | 0.620294 | 5.688938 | 0.0003 | 0.01229 |
| ENSG00000131069 | ACSS2 | 53.62391545 | 59.95243815 | 60.40257648 | 37.20468912 | 36.80867655 | 38.54805 | 0.622779 | 5.609724 | 0.00031 | 0.01245 |
| ENSG00000111678 | C12orf57 | 72.41520206 | 54.9097097 | 62.68191899 | 90.42152291 | 87.17844446 | 123.565 | -0.66141 | 6.374979 | 0.00031 | 0.01245 |
| ENSG00000218175 | AC016739.2 | 161.7883945 | 100.8545688 | 195.4536201 | 239.2402794 | 245.391177 | 274.0608 | -0.72726 | 7.670851 | 0.00031 | 0.01251 |
| ENSG00000159461 | AMFR | 74.70682237 | 86.28668668 | 60.40257648 | 41.91414343 | 41.97480659 | 53.86166 | 0.683061 | 5.931589 | 0.00032 | 0.01288 |
| ENSG00000152661 | GJA1 | 395.5336669 | 417.9861575 | 403.4436241 | 541.1163012 | 519.1960692 | 556.5705 | -0.41062 | 8.886697 | 0.00032 | 0.01295 |
| ENSG00000003137 | CYP26B1 | 7.333185018 | 14.56788217 | 9.117370035 | 2.354727159 | 4.520363787 | 3.168333 | 1.631924 | 2.967617 | 0.00033 | 0.01304 |
| ENSG00000105221 | AKT2 | 186.5378939 | 184.9000429 | 174.9395375 | 130.4518846 | 137.5482124 | 125.6772 | 0.474951 | 7.301898 | 0.00033 | 0.01316 |
| ENSG00000237793 | RPL18AP16 | 3.208268446 | 3.922122122 | 7.407863154 | 10.83174493 | 13.56109136 | 12.14528 | -1.33751 | 3.237892 | 0.00033 | 0.01322 |
| ENSG00000159692 | CTBP1 | 161.3300704 | 143.9979122 | 158.4143044 | 201.0936994 | 224.7266568 | 219.143 | -0.47301 | 7.537011 | 0.00033 | 0.01323 |
| ENSG00000204310 | AGPAT1 | 17.41631442 | 15.12818533 | 17.09506882 | 7.535126909 | 9.686493828 | 6.864722 | 1.059776 | 3.740904 | 0.00034 | 0.01336 |
| ENSG00000236552 | RPL13AP5 | 112.7477197 | 87.96759616 | 107.6989335 | 125.7424303 | 169.1907589 | 180.0669 | -0.61777 | 7.038086 | 0.00034 | 0.01336 |
| ENSG00000184602 | SNN | 6.416536891 | 4.482425282 | 5.128520645 | 0.941890864 | 1.29153251 | 1.584167 | 2.046906 | 2.144823 | 0.00034 | 0.01336 |
| ENSG00000112996 | MRPS30 | 18.79128661 | 19.61061061 | 20.51408258 | 37.20468912 | 34.22561153 | 29.57111 | -0.78113 | 4.790243 | 0.00035 | 0.01354 |
| ENSG00000241656 | UBA52P7 | 71.04022986 | 68.91728871 | 76.35797404 | 96.54381353 | 109.1344971 | 108.2514 | -0.53479 | 6.480318 | 0.00035 | 0.01357 |
| ENSG00000205105 | COX17P1 | 82.95665552 | 74.52032031 | 76.35797404 | 106.4336676 | 113.0090947 | 117.2283 | -0.52167 | 6.5863 | 0.00035 | 0.01357 |
| ENSG00000237528 | RP11-234P3.4 | 78.37341488 | 56.59061918 | 108.8386048 | 116.7944671 | 149.172005 | 143.103 | -0.74512 | 6.775705 | 0.00035 | 0.01363 |
| ENSG00000136560 | TANK | 24.74949944 | 23.53273273 | 23.93309634 | 9.418908637 | 14.20685761 | 14.78555 | 0.927993 | 4.285311 | 0.00036 | 0.01394 |
| ENSG00000101474 | APMAP | 16.04134223 | 22.97242957 | 24.50293197 | 12.24458123 | 10.33226008 | 7.920833 | 1.033683 | 4.060899 | 0.00037 | 0.01404 |
| ENSG00000167720 | SRR | 5.499888764 | 6.723637923 | 6.268191899 | 1.883781727 | 1.29153251 | 2.112222 | 1.732393 | 2.335671 | 0.00037 | 0.01412 |
| ENSG00000238181 | AHCYP2 | 2.749944382 | 4.482425282 | 4.558685018 | 0.470945432 | 1.29153251 | 0 | 2.694595 | 1.714978 | 0.00037 | 0.01421 |
| ENSG00000196628 | TCF4 | 107.706155 | 110.3797226 | 127.6431805 | 169.06941 | 152.4008362 | 166.8655 | -0.50227 | 7.129694 | 0.00037 | 0.01421 |
| ENSG00000109819 | PPARGC1A | 140.2471635 | 142.3170027 | 131.6320299 | 204.8612629 | 170.4822914 | 211.2222 | -0.50439 | 7.391941 | 0.00038 | 0.01429 |

| | | | | | | | | | | | |
|---|---|---|---|---|---|---|---|---|---|---|---|
| ENSG00000166411 | IDH3A | 604.987764 | 697.5774345 | 584.0815179 | 416.7867072 | 514.6757054 | 384.9525 | 0.520608 | 9.062839 | 0.00038 | 0.01445 |
| ENSG00000161016 | RPL8 | 456.0324433 | 387.1694837 | 444.4717892 | 624.4736426 | 526.299498 | 640.0033 | -0.47627 | 9.006377 | 0.00038 | 0.01448 |
| ENSG00000004779 | NDUFAB1 | 514.6979235 | 438.1570713 | 524.8186126 | 594.8040804 | 775.5652725 | 732.9411 | -0.50802 | 9.223084 | 0.00039 | 0.01465 |
| ENSG00000130347 | RTN4IP1 | 60.04045234 | 54.34940654 | 54.70422021 | 37.67563455 | 34.87137778 | 38.02 | 0.611384 | 5.579301 | 0.0004 | 0.01485 |
| ENSG00000145349 | CAMK2D | 114.5810159 | 113.1812384 | 91.74353598 | 147.8768656 | 158.8584988 | 150.4958 | -0.51275 | 7.027323 | 0.0004 | 0.01492 |
| ENSG00000244005 | NFS1 | 46.29073043 | 53.22880022 | 46.72652143 | 29.66956221 | 32.93407902 | 30.09917 | 0.659582 | 5.353189 | 0.00041 | 0.01525 |
| ENSG00000214318 | ATP5G1P6 | 129.70571 | 112.060632 | 197.163127 | 231.234207 | 209.2282667 | 277.2292 | -0.7117 | 7.598138 | 0.00041 | 0.01527 |
| ENSG00000113441 | LNPEP | 53.62391545 | 49.86698126 | 51.85504207 | 78.64788712 | 72.32582059 | 78.68027 | -0.56454 | 6.027771 | 0.00042 | 0.01532 |
| ENSG00000102225 | CDK16 | 170.4965517 | 165.8497354 | 160.1238112 | 114.4397399 | 119.4667572 | 124.6211 | 0.470991 | 7.166341 | 0.00042 | 0.01544 |
| ENSG00000171823 | FBXL14 | 15.58301816 | 14.56788217 | 18.23474007 | 8.947963205 | 5.811896297 | 7.920833 | 1.057328 | 3.699591 | 0.00042 | 0.01547 |
| ENSG00000231500 | RPS18 | 287.827512 | 241.4906621 | 319.6777869 | 363.5698734 | 395.8547145 | 493.2039 | -0.56088 | 8.456284 | 0.00042 | 0.01549 |
| ENSG00000166908 | PIP4K2C | 28.87441601 | 36.98000858 | 36.46948014 | 19.30876271 | 18.08145515 | 21.65028 | 0.778792 | 4.804486 | 0.00043 | 0.01553 |
| ENSG00000240463 | RPS19P3 | 178.2880608 | 104.776691 | 129.3526874 | 200.622754 | 204.7079029 | 250.8264 | -0.6658 | 7.486032 | 0.00043 | 0.01553 |
| ENSG00000226617 | RPL21P110 | 20.62458286 | 13.44727585 | 22.22358946 | 33.90807109 | 31.64254651 | 32.73944 | -0.79928 | 4.745766 | 0.00043 | 0.01553 |
| ENSG00000174572 | RPL10P2 | 15.58301816 | 12.88697269 | 13.67605505 | 26.84388961 | 26.47641646 | 22.70639 | -0.83927 | 4.373972 | 0.00045 | 0.01626 |
| ENSG00000120833 | SOCS2 | 14.66637004 | 18.49000429 | 13.67605505 | 27.78578048 | 29.70524774 | 25.34667 | -0.81954 | 4.496404 | 0.00045 | 0.01633 |
| ENSG00000179010 | MRFAP1 | 79.29006301 | 64.43486343 | 86.61501533 | 104.5498859 | 131.0905498 | 109.8356 | -0.57897 | 6.596859 | 0.00046 | 0.01644 |
| ENSG00000226525 | RPS7P10 | 98.08134962 | 77.32183611 | 73.50879591 | 128.5681029 | 117.5294585 | 119.3406 | -0.54914 | 6.694313 | 0.00047 | 0.01668 |
| ENSG00000204434 | POTEKP | 95.7897293 | 89.64850564 | 104.8497554 | 51.33305207 | 69.74275556 | 71.2875 | 0.600135 | 6.347775 | 0.00047 | 0.01668 |
| ENSG00000175054 | ATR | 33.91598071 | 30.81667381 | 27.35211011 | 44.73981602 | 48.43246914 | 53.33361 | -0.66036 | 5.350165 | 0.00047 | 0.01672 |
| ENSG00000149428 | HYOU1 | 36.66592509 | 42.58304018 | 41.59800079 | 28.25672591 | 20.66452017 | 23.7625 | 0.71276 | 5.062221 | 0.00048 | 0.01685 |
| ENSG00000214121 | PRDX1P1 | 64.16536891 | 86.28668668 | 71.2294534 | 31.55334393 | 58.76472923 | 41.18833 | 0.766593 | 5.898652 | 0.00048 | 0.01687 |
| ENSG00000213569 | RP13-383K5.4 | 1.833296255 | 1.680909481 | 2.279342509 | 6.593236046 | 7.103428807 | 5.808611 | -1.71635 | 2.387865 | 0.00048 | 0.01691 |
| ENSG00000116754 | SRSF11 | 47.66570262 | 39.78152438 | 21.65375383 | 61.69385157 | 60.05626174 | 69.17527 | -0.79565 | 5.677262 | 0.00048 | 0.017 |
| ENSG00000062282 | DGAT2 | 296.5356692 | 270.6264264 | 264.403731 | 206.7450446 | 209.2282667 | 200.6611 | 0.432872 | 7.922167 | 0.00052 | 0.01805 |
| ENSG00000101444 | AHCY | 24.74949944 | 26.33424853 | 43.87734329 | 15.07025382 | 19.37298766 | 13.72944 | 0.979936 | 4.630443 | 0.00052 | 0.01814 |
| ENSG00000137692 | DCUN1D5 | 85.7065999 | 82.92486772 | 60.97241211 | 104.5498859 | 123.3413547 | 115.6442 | -0.57351 | 6.592129 | 0.00053 | 0.01843 |
| ENSG00000140983 | RHOT2 | 48.12402668 | 39.22122122 | 50.14553519 | 28.25672591 | 30.99678025 | 26.40278 | 0.688627 | 5.259362 | 0.00054 | 0.0187 |
| ENSG00000005882 | PDK2 | 303.8688542 | 285.7546117 | 339.6220338 | 182.7268276 | 240.2250469 | 230.7603 | 0.509943 | 8.048735 | 0.00054 | 0.0187 |
| ENSG00000131779 | PEX11B | 25.2078235 | 17.36939797 | 23.36326072 | 9.889854069 | 14.85262387 | 8.448889 | 1.019057 | 4.138353 | 0.00055 | 0.01877 |
| ENSG00000234782 | TPT1P9 | 149.8719688 | 118.2239668 | 151.0064412 | 204.8612629 | 225.3724231 | 177.9547 | -0.53394 | 7.427363 | 0.00055 | 0.01891 |
| ENSG00000241590 | RPL17P37 | 21.08290693 | 10.08545688 | 14.81572631 | 23.54727159 | 32.28831276 | 34.32361 | -0.94155 | 4.566173 | 0.00055 | 0.01895 |
| ENSG00000108669 | CYTH1 | 43.08246198 | 64.43486343 | 46.72652143 | 31.0823985 | 27.76794897 | 33.79555 | 0.723754 | 5.399676 | 0.00056 | 0.01899 |
| ENSG00000188807 | TMEM201 | 21.54123099 | 17.36939797 | 23.93309634 | 9.418908637 | 9.686493828 | 13.20139 | 0.954963 | 4.08602 | 0.00057 | 0.01946 |
| ENSG00000165416 | SUGT1 | 38.04089728 | 34.17849277 | 30.20128824 | 62.63574243 | 48.43246914 | 50.69333 | -0.65995 | 5.499066 | 0.00058 | 0.01974 |
| ENSG00000083845 | RPS5 | 369.8675194 | 282.3927928 | 356.147267 | 488.8413582 | 422.3311309 | 522.2469 | -0.50739 | 8.673022 | 0.00059 | 0.01999 |
| ENSG00000100897 | DCAF11 | 121.9142009 | 113.7415415 | 109.4084404 | 72.99654194 | 92.99034075 | 71.81555 | 0.546243 | 6.617548 | 0.00061 | 0.02046 |
| ENSG00000116016 | EPAS1 | 170.0382276 | 165.8497354 | 196.0234558 | 225.5828619 | 238.2877482 | 305.7442 | -0.53371 | 7.767393 | 0.00062 | 0.02065 |
| ENSG00000118729 | CASQ2 | 225.4954393 | 236.4479336 | 177.7887157 | 164.3599557 | 144.0058749 | 132.0139 | 0.537849 | 7.502139 | 0.00062 | 0.02065 |
| ENSG00000235605 | RP5-827C21.1 | 68.74860955 | 32.49758329 | 43.30750767 | 74.40937823 | 74.26311935 | 124.0931 | -0.90783 | 6.145608 | 0.00062 | 0.0208 |
| ENSG00000109472 | CPE | 249.7866147 | 192.7442871 | 201.1519764 | 145.9930839 | 165.3161613 | 138.3505 | 0.522283 | 7.519153 | 0.00063 | 0.02084 |
| ENSG00000105372 | RPS19 | 267.6612532 | 189.3824682 | 250.1578403 | 297.6375129 | 350.6510766 | 404.4905 | -0.57194 | 8.201172 | 0.00063 | 0.02084 |
| ENSG00000197296 | FITM2 | 43.99911011 | 54.9097097 | 57.55339835 | 32.96618023 | 25.83065021 | 37.49194 | 0.682505 | 5.433949 | 0.00063 | 0.02097 |
| ENSG00000107371 | EXOSC3 | 5.0415647 | 6.723637923 | 2.279342509 | 12.24458123 | 12.26955885 | 10.03306 | -1.27544 | 3.190775 | 0.00064 | 0.02107 |
| ENSG00000217128 | FNIP1 | 49.49899887 | 47.06546546 | 42.73767204 | 69.22897848 | 61.34779425 | 80.26444 | -0.59838 | 5.894559 | 0.00064 | 0.02107 |
| ENSG00000144191 | CNGA3 | 3.666592509 | 3.361818961 | 4.558685018 | 0.941890864 | 0.645766255 | 0.528056 | 2.317368 | 1.750624 | 0.00065 | 0.02114 |
| ENSG00000100403 | ZC3H7B | 73.33185018 | 101.9751752 | 83.7658372 | 46.15265232 | 48.43246914 | 68.64722 | 0.661438 | 6.158185 | 0.00065 | 0.02114 |
| ENSG00000185624 | P4HB | 80.20671114 | 90.2088088 | 73.50879591 | 50.86210664 | 63.93085927 | 46.99694 | 0.600702 | 6.100623 | 0.00065 | 0.02116 |
| ENSG00000107863 | ARHGAP21 | 46.29073043 | 34.73879593 | 42.16783641 | 70.64181478 | 49.72400165 | 78.68027 | -0.6949 | 5.780996 | 0.00065 | 0.02116 |
| ENSG00000070159 | PTPN3 | 49.04067481 | 63.31425711 | 43.87734329 | 33.90807109 | 27.12218272 | 34.85167 | 0.692543 | 5.432847 | 0.00068 | 0.02194 |
| ENSG00000122550 | KLHL7 | 70.12358174 | 71.71880451 | 59.83274086 | 39.08847084 | 44.55787161 | 49.63722 | 0.602246 | 5.831775 | 0.00068 | 0.02203 |
| ENSG00000215895 | RP11-334L9.1 | 75.16514644 | 91.88971828 | 74.64846716 | 45.68170689 | 60.70202799 | 53.86166 | 0.599651 | 6.086457 | 0.00068 | 0.02203 |
| ENSG00000056586 | RC3H2 | 120.0809047 | 118.78427 | 106.5592623 | 171.8950826 | 145.2974074 | 164.2253 | -0.4805 | 7.118594 | 0.00069 | 0.02207 |
| ENSG00000015413 | DPEP1 | 13.29139785 | 7.844244243 | 13.10621943 | 4.709454318 | 5.166130042 | 4.224444 | 1.285035 | 3.200321 | 0.0007 | 0.02241 |
| ENSG00000077312 | SNRPA | 43.08246198 | 22.41212641 | 41.02816516 | 53.2168338 | 61.34779425 | 64.95083 | -0.742 | 5.605419 | 0.0007 | 0.02248 |
| ENSG00000196177 | ACADSB | 22.91620318 | 24.65333905 | 26.78227448 | 12.71552666 | 14.20685761 | 15.31361 | 0.812691 | 4.356392 | 0.00071 | 0.02262 |
| ENSG00000228399 | RP4-575N6.2 | 41.70748979 | 42.02273702 | 33.05046638 | 52.74588837 | 66.51392429 | 61.25444 | -0.61625 | 5.657028 | 0.00071 | 0.02264 |
| ENSG00000177303 | CASKIN2 | 39.87419354 | 38.66091806 | 47.29635706 | 27.78578048 | 24.5391177 | 25.87472 | 0.67603 | 5.13441 | 0.00072 | 0.02271 |
| ENSG00000084754 | HADHA | 525.2393769 | 528.3658801 | 490.0586394 | 397.9488899 | 401.6666107 | 377.5597 | 0.391465 | 8.828486 | 0.00072 | 0.02271 |
| ENSG00000136938 | ANP32B | 33.91598071 | 27.45485485 | 35.32980889 | 49.92021578 | 48.43246914 | 51.74944 | -0.6322 | 5.398491 | 0.00072 | 0.02282 |
| ENSG00000134324 | LPIN1 | 40.3325176 | 61.63334763 | 49.00586394 | 32.4952348 | 27.76794897 | 31.68333 | 0.700813 | 5.376177 | 0.00074 | 0.02307 |
| ENSG00000120053 | GOT1 | 463.3656283 | 427.5113113 | 469.5445568 | 307.0564216 | 382.2936231 | 294.1269 | 0.470549 | 8.613393 | 0.00074 | 0.02307 |
| ENSG00000066933 | MYO9A | 23.83285131 | 17.36939797 | 16.52523319 | 40.03036171 | 31.64254651 | 29.04305 | -0.794 | 4.786275 | 0.00074 | 0.02307 |
| ENSG00000224059 | HSPA8P16 | 9.166481273 | 8.964850564 | 11.96654817 | 18.36687184 | 18.08145515 | 20.06611 | -0.91426 | 3.948878 | 0.00074 | 0.0231 |
| ENSG00000103966 | EHD4 | 216.7872821 | 231.9655083 | 221.0962234 | 152.1153745 | 165.3161613 | 176.3705 | 0.440372 | 7.607405 | 0.00074 | 0.02316 |
| ENSG00000165795 | NDRG2 | 1205.392287 | 987.8144715 | 1000.631361 | 751.1579638 | 854.9945219 | 738.2216 | 0.447207 | 9.852089 | 0.00075 | 0.02319 |
| ENSG00000169116 | PARM1 | 24.29117537 | 20.73121693 | 18.8045757 | 24.48916246 | 50.36976791 | 47.525 | -0.91865 | 4.98976 | 0.00077 | 0.0237 |
| ENSG00000100226 | GTPBP1 | 34.83262884 | 42.58304018 | 41.59800079 | 25.90199875 | 22.60181893 | 25.34667 | 0.673655 | 5.05428 | 0.00077 | 0.02375 |
| ENSG00000213700 | RPL17P50 | 12.37474972 | 9.525153724 | 16.52523319 | 21.19254443 | 25.18488395 | 24.81861 | -0.88351 | 4.263785 | 0.00079 | 0.02416 |
| ENSG00000136238 | RAC1 | 75.16514644 | 85.16608036 | 97.44189225 | 103.607995 | 141.4228099 | 140.4628 | -0.5796 | 6.75383 | 0.00079 | 0.02421 |
| ENSG00000139914 | FITM1 | 85.24827584 | 56.59061918 | 80.91665906 | 106.4336676 | 113.0090947 | 113.0039 | -0.57194 | 6.547945 | 0.0008 | 0.02452 |
| ENSG00000184983 | NDUFA6 | 336.8681868 | 262.221879 | 378.3708565 | 427.6184521 | 525.6537318 | 458.8803 | -0.52944 | 8.640495 | 0.0008 | 0.02452 |
| ENSG00000217239 | RP1-182O16.1 | 22.45787912 | 6.163334763 | 17.66490444 | 27.31483505 | 31.64254651 | 42.7725 | -1.11588 | 4.686643 | 0.00081 | 0.02452 |
| ENSG00000236801 | RPL24P8 | 78.83173895 | 51.54789074 | 79.20715218 | 104.5498859 | 94.92763952 | 126.2053 | -0.63454 | 6.497629 | 0.00081 | 0.02452 |
| ENSG00000159640 | ACE | 13.74972191 | 26.33424853 | 26.21243885 | 11.30269036 | 7.749195063 | 11.61722 | 1.075254 | 4.100836 | 0.00083 | 0.02506 |

| | | | | | | | | | | | |
|---|---|---|---|---|---|---|---|---|---|---|---|
| ENSG00000198258 | UBL5 | 33.91598071 | 21.29152009 | 49.00586394 | 57.92628812 | 52.95283293 | 78.15222 | -0.86195 | 5.641579 | 0.00083 | 0.02507 |
| ENSG00000117153 | KLHL12 | 27.95776788 | 26.89455169 | 27.35211011 | 19.77970814 | 14.85262387 | 12.14528 | 0.798201 | 4.503388 | 0.00084 | 0.02507 |
| ENSG00000112531 | QKI | 617.3625137 | 616.3334763 | 567.5562847 | 803.9038522 | 725.8412709 | 853.3377 | -0.40416 | 9.44815 | 0.00084 | 0.02507 |
| ENSG00000176171 | BNIP3 | 60.4987764 | 59.39213499 | 54.70422021 | 32.96618023 | 45.84940412 | 35.90778 | 0.621277 | 5.621752 | 0.00084 | 0.02507 |
| ENSG00000232472 | EEF1B2P3 | 43.54078605 | 31.37697697 | 23.36326072 | 48.97832491 | 52.30706667 | 64.42277 | -0.73839 | 5.496194 | 0.00087 | 0.02586 |
| ENSG00000132819 | RBM38 | 49.49899887 | 52.1081939 | 60.40257648 | 34.37901652 | 38.74597531 | 32.73944 | 0.614981 | 5.512487 | 0.00088 | 0.02609 |
| ENSG00000239559 | RPL37P2 | 3.208268446 | 3.361818961 | 6.268191899 | 9.418908637 | 10.97802634 | 10.56111 | -1.27342 | 3.047052 | 0.00088 | 0.02609 |
| ENSG00000091428 | RAPGEF4 | 38.04089728 | 41.46243386 | 37.03931577 | 24.01821702 | 27.12218272 | 22.17833 | 0.673823 | 5.031336 | 0.00088 | 0.02609 |
| ENSG00000114853 | ZBTB47 | 77.45676676 | 49.86698126 | 64.9612615 | 39.08847084 | 40.68327408 | 42.7725 | 0.656444 | 5.748075 | 0.00088 | 0.02609 |
| ENSG00000152782 | PANK1 | 44.91575824 | 57.7112255 | 48.43602831 | 32.96618023 | 34.87137778 | 29.04305 | 0.638934 | 5.404045 | 0.00089 | 0.02609 |
| ENSG00000214745 | RP11-54I5.1 | 21.08290693 | 11.76636636 | 18.8045757 | 8.947963205 | 9.686493828 | 2.640278 | 1.291448 | 3.735971 | 0.0009 | 0.02637 |
| ENSG00000204228 | HSD17B8 | 101.7479421 | 75.08062347 | 83.7658372 | 137.9870115 | 124.6328873 | 115.1161 | -0.53155 | 6.748797 | 0.0009 | 0.02638 |
| ENSG00000109171 | SLAIN2 | 208.0791249 | 184.9000429 | 200.0123051 | 281.1544228 | 248.6200083 | 266.14 | -0.4251 | 7.861824 | 0.00091 | 0.02665 |
| ENSG00000164032 | H2AFZ | 127.8724138 | 143.9979122 | 101.4307416 | 182.7268276 | 193.7298766 | 162.6411 | -0.52782 | 7.257496 | 0.00092 | 0.02669 |
| ENSG00000125875 | TBC1D20 | 21.99955505 | 40.9021307 | 27.35211011 | 19.77970814 | 13.56109136 | 12.67333 | 0.94667 | 4.571068 | 0.00093 | 0.02696 |
| ENSG00000233839 | RP11-389O22.4 | 6.416536891 | 3.922122122 | 7.407863154 | 10.3607995 | 21.31028642 | 12.14528 | -1.26243 | 3.468568 | 0.00093 | 0.02696 |
| ENSG00000243859 | RPL5P17 | 16.95799035 | 15.12818533 | 16.52523319 | 22.1344353 | 28.41371523 | 39.07611 | -0.87344 | 4.58517 | 0.00095 | 0.0275 |
| ENSG00000198925 | ATG9A | 78.37341488 | 71.15850135 | 96.87205662 | 64.99046959 | 37.4544428 | 51.22139 | 0.665682 | 6.087217 | 0.00095 | 0.0275 |
| ENSG00000087088 | BAX | 32.08268446 | 27.45485485 | 35.32980889 | 54.62967009 | 40.68327408 | 53.86166 | -0.66146 | 5.386984 | 0.00095 | 0.0275 |
| ENSG00000121898 | CPXM2 | 38.95754541 | 32.49758329 | 37.03931577 | 17.42498098 | 23.89335144 | 24.81861 | 0.728509 | 4.916139 | 0.00096 | 0.02758 |
| ENSG00000100266 | PACSIN2 | 180.5796811 | 162.4879165 | 165.2523319 | 129.9809392 | 109.7802634 | 128.8455 | 0.460834 | 7.204017 | 0.00096 | 0.02761 |
| ENSG00000082641 | NFE2L1 | 776.4009638 | 782.1832117 | 671.2663688 | 562.779791 | 551.484382 | 580.8611 | 0.395654 | 9.356155 | 0.00099 | 0.02809 |
| ENSG00000214199 | EEF1A1P12 | 11.91642565 | 12.32666953 | 10.25704129 | 20.721599 | 23.89335144 | 18.48194 | -0.85479 | 4.107077 | 0.00099 | 0.02809 |
| ENSG00000126882 | FAM78A | 13.29139785 | 16.24879165 | 11.96654817 | 8.006072341 | 5.166130042 | 6.336666 | 1.053199 | 3.500002 | 0.00099 | 0.02809 |
| ENSG00000140479 | PCSK6 | 24.29117537 | 29.13576433 | 25.64260322 | 13.18647209 | 13.56109136 | 18.48194 | 0.797198 | 4.446098 | 0.00099 | 0.02809 |
| ENSG00000144043 | TEX261 | 10.0831294 | 8.404547404 | 15.38556193 | 5.651345182 | 1.937298766 | 5.280555 | 1.325772 | 3.158693 | 0.00099 | 0.02809 |
| ENSG00000127314 | RAP1B | 42.62413792 | 33.61818961 | 25.0727676 | 45.68170689 | 55.53589795 | 68.64722 | -0.73108 | 5.531302 | 0.001 | 0.0282 |
| ENSG00000108861 | DUSP3 | 19.24961067 | 24.09303589 | 19.37441132 | 14.12836296 | 8.394961318 | 10.56111 | 0.887103 | 4.098975 | 0.00102 | 0.02854 |
| ENSG00000160392 | C19orf47 | 29.79106414 | 25.21364221 | 26.21243885 | 14.12836296 | 14.85262387 | 18.48194 | 0.778832 | 4.498943 | 0.00102 | 0.02854 |
| ENSG00000163584 | RPL22L1 | 65.08201704 | 54.34940654 | 82.05633032 | 82.41545057 | 106.5514321 | 130.9578 | -0.66559 | 6.455236 | 0.00102 | 0.02854 |
| ENSG00000143363 | PRUNE1 | 10.0831294 | 11.76636636 | 13.67605505 | 5.18039975 | 5.166130042 | 5.808611 | 1.113779 | 3.272757 | 0.00102 | 0.02854 |
| ENSG00000243181 | RP11-734J24.1 | 5.0415647 | 2.241212641 | 2.849178136 | 8.477017773 | 7.749195063 | 10.56111 | -1.34999 | 2.858845 | 0.00103 | 0.02876 |
| ENSG00000249754 | NDUFB4P9 | 4.583240636 | 1.680909481 | 6.268191899 | 10.83174493 | 12.26955885 | 8.448889 | -1.30526 | 3.060463 | 0.00104 | 0.02892 |
| ENSG00000141756 | FKBP10 | 23.37452725 | 26.33424853 | 21.65375383 | 16.01214468 | 11.62379259 | 12.14528 | 0.82215 | 4.299667 | 0.00107 | 0.02968 |
| ENSG00000132313 | MRPL35 | 2.749944382 | 4.482425282 | 4.558685018 | 11.30269036 | 10.97802634 | 7.392777 | -1.34177 | 2.982116 | 0.00107 | 0.02973 |
| ENSG00000267590 | NDUFA3P1 | 85.24827584 | 83.48517088 | 116.2464679 | 165.3018466 | 113.6548609 | 153.6642 | -0.60906 | 6.916431 | 0.00108 | 0.02976 |
| ENSG00000198034 | RPS4X | 397.3669632 | 292.4782496 | 373.8121714 | 511.446739 | 461.7228725 | 499.5405 | -0.46891 | 8.727552 | 0.00108 | 0.02976 |
| ENSG00000217733 | CCT7P1 | 20.62458286 | 14.56788217 | 13.67605505 | 8.006072341 | 10.33226008 | 4.7525 | 1.110222 | 3.717267 | 0.00109 | 0.02999 |
| ENSG00000109501 | WFS1 | 92.12313679 | 91.32941512 | 97.44189225 | 79.11883255 | 54.89013169 | 53.86166 | 0.569771 | 6.311032 | 0.00111 | 0.03039 |
| ENSG00000277443 | MARCKS | 114.5810159 | 85.16608036 | 99.15139913 | 138.457957 | 125.9244198 | 184.2914 | -0.58454 | 6.974895 | 0.00113 | 0.03079 |
| ENSG00000138796 | HADH | 166.8299592 | 150.1612469 | 141.8890712 | 120.0910851 | 112.3633284 | 95.57805 | 0.485383 | 7.048596 | 0.00113 | 0.03079 |
| ENSG00000196531 | NACA | 237.411865 | 176.4954955 | 240.4706347 | 310.823985 | 291.2405811 | 311.5528 | -0.48086 | 8.035824 | 0.00114 | 0.03092 |
| ENSG00000219470 | RP3-337H4.6 | 154.4552094 | 105.3369941 | 157.2746331 | 209.0997717 | 189.855279 | 208.0539 | -0.5406 | 7.424689 | 0.00115 | 0.03117 |
| ENSG00000174748 | RPL15 | 44.45743417 | 49.86698126 | 53.56454896 | 62.164797 | 74.90888561 | 83.96083 | -0.58005 | 5.962319 | 0.00116 | 0.0312 |
| ENSG00000092964 | DPYSL2 | 51.33229513 | 62.75395395 | 66.10093275 | 76.29315996 | 83.94961318 | 121.9808 | -0.65128 | 6.285017 | 0.00116 | 0.0312 |
| ENSG00000108262 | GIT1 | 64.16536891 | 62.19365079 | 64.9612615 | 44.73981602 | 41.32904033 | 45.41278 | 0.536919 | 5.780263 | 0.00116 | 0.0312 |
| ENSG00000173221 | GLRX | 7.791509082 | 6.163334763 | 8.547534408 | 0.941890864 | 4.520363787 | 2.112222 | 1.643828 | 2.592895 | 0.00117 | 0.0312 |
| ENSG00000179152 | TCAIM | 16.49966629 | 17.36939797 | 17.09506882 | 11.30269036 | 9.040727573 | 4.7525 | 1.005293 | 3.786151 | 0.00117 | 0.0312 |
| ENSG00000259956 | RBM15B | 7.791509082 | 12.88697269 | 14.24589068 | 4.709454318 | 3.874597531 | 5.808611 | 1.233932 | 3.207183 | 0.00117 | 0.0312 |
| ENSG00000166326 | TRIM44 | 36.20760103 | 26.89455169 | 30.77112387 | 51.33305207 | 41.32904033 | 55.44583 | -0.6563 | 5.376127 | 0.00117 | 0.0312 |
| ENSG00000172346 | CSDC2 | 17.41631442 | 19.05030745 | 25.0727676 | 9.418908637 | 12.9153251 | 10.56111 | 0.907968 | 4.062884 | 0.00118 | 0.03137 |
| ENSG00000033627 | ATP6V0A1 | 88.91486835 | 92.45002144 | 92.88320723 | 59.33912441 | 72.32582059 | 61.7825 | 0.509478 | 6.302838 | 0.00119 | 0.03162 |
| ENSG00000227590 | ATP5G1P5 | 114.5810159 | 82.92486772 | 119.0956461 | 152.1153745 | 138.8397449 | 170.0339 | -0.54178 | 7.030397 | 0.00121 | 0.03216 |
| ENSG00000168256 | NKIRAS2 | 32.99933258 | 28.01515801 | 31.91079512 | 20.721599 | 20.66452017 | 14.2575 | 0.743836 | 4.694574 | 0.00122 | 0.03219 |
| ENSG00000129084 | PSMA1 | 19.24961067 | 12.88697269 | 10.25704129 | 19.30876271 | 34.87137778 | 29.57111 | -0.94537 | 4.453247 | 0.00122 | 0.03219 |
| ENSG00000136149 | RPL13AP25 | 60.04045234 | 53.22880022 | 63.25175462 | 71.58370564 | 95.57340577 | 93.99388 | -0.55793 | 6.205241 | 0.00122 | 0.03219 |
| ENSG00000128524 | ATP6V1F | 16.49966629 | 15.12818533 | 22.22358946 | 24.96010789 | 30.99678025 | 38.54805 | -0.8104 | 4.68052 | 0.00123 | 0.03219 |
| ENSG00000170837 | GPR27 | 5.0415647 | 5.042728442 | 7.977698781 | 0.941890864 | 1.937298766 | 2.640278 | 1.690864 | 2.312457 | 0.00127 | 0.03324 |
| ENSG00000145495 | 44626 | 119.6225806 | 151.2818533 | 139.6097287 | 173.3079189 | 190.5010453 | 204.8855 | -0.47202 | 7.358087 | 0.00128 | 0.03336 |
| ENSG00000138029 | HADHB | 2090.874378 | 2188.544144 | 1928.893598 | 1393.998478 | 1778.440267 | 1418.357 | 0.436046 | 10.81444 | 0.00129 | 0.03354 |
| ENSG00000089597 | GANAB | 43.99911011 | 51.54789074 | 47.29635706 | 34.84996196 | 29.70524774 | 29.04305 | 0.59853 | 5.34072 | 0.00129 | 0.03366 |
| ENSG00000011105 | TSPAN9 | 27.04111976 | 47.06546546 | 44.44717892 | 22.60538073 | 15.49839013 | 26.93083 | 0.841679 | 4.983159 | 0.00131 | 0.03386 |
| ENSG00000132155 | RAF1 | 147.5803485 | 131.1109395 | 110.5481117 | 100.7823224 | 82.01231441 | 88.71333 | 0.517923 | 6.800771 | 0.00131 | 0.03397 |
| ENSG00000205544 | TMEM256 | 41.24916573 | 34.17849277 | 74.07863154 | 76.76410539 | 78.13771688 | 104.0269 | -0.79689 | 6.107981 | 0.00132 | 0.03398 |
| ENSG00000115415 | STAT1 | 16.04134223 | 20.73121693 | 17.09506882 | 10.83174493 | 9.040727573 | 8.976944 | 0.879172 | 3.894992 | 0.00132 | 0.03398 |
| ENSG00000113583 | C5orf15 | 13.29139785 | 12.88697269 | 23.36326072 | 31.55334393 | 34.22561153 | 24.29055 | -0.86752 | 4.593633 | 0.00132 | 0.03398 |
| ENSG00000125375 | ATP5S | 10.0831294 | 7.844244243 | 7.977698781 | 16.95403555 | 15.49839013 | 16.36972 | -0.89987 | 3.759201 | 0.00133 | 0.03423 |
| ENSG00000123009 | NME2P1 | 8.249833146 | 5.042728442 | 11.39671254 | 15.07025382 | 22.60181893 | 14.78555 | -1.06509 | 3.778602 | 0.00134 | 0.03427 |
| ENSG00000071553 | ATP6AP1 | 79.29006301 | 62.75395395 | 76.35797404 | 53.2168338 | 48.43246914 | 48.05305 | 0.544266 | 5.967484 | 0.00134 | 0.0343 |
| ENSG00000145362 | ANK2 | 137.9555432 | 158.5657943 | 149.2969343 | 227.4666436 | 169.8365251 | 234.4567 | -0.50762 | 7.498126 | 0.00135 | 0.03432 |
| ENSG00000230779 | RP1-140J1.4 | 29.33274007 | 20.17091377 | 21.08391821 | 39.55941627 | 35.51714404 | 39.07611 | -0.6813 | 4.997745 | 0.00136 | 0.03449 |
| ENSG00000120658 | ENOX1 | 1.374972191 | 1.680909481 | 2.849178136 | 0 | 0 | 0 | 4.908257 | 1.027579 | 0.00138 | 0.03488 |
| ENSG00000243199 | RP11-408P14.1 | 49.49899887 | 46.5051623 | 37.6091514 | 65.93236046 | 58.76472923 | 75.51194 | -0.58132 | 5.82758 | 0.00139 | 0.03511 |
| ENSG00000182899 | RPL35A | 334.5765665 | 289.1164307 | 364.1249658 | 408.7806348 | 432.0176247 | 543.8972 | -0.48716 | 8.63087 | 0.00139 | 0.03515 |
| ENSG00000182700 | IGIP | 33.45765665 | 25.21364221 | 22.79342509 | 40.97225257 | 41.97480659 | 46.46889 | -0.65289 | 5.18002 | 0.0014 | 0.03519 |

| | | | | | | | | | | | |
|---|---|---|---|---|---|---|---|---|---|---|---|
| ENSG00000132824 | SERINC3 | 74.24849831 | 96.37214356 | 68.38027526 | 48.03643405 | 62.63932676 | 46.46889 | 0.611823 | 6.065594 | 0.00141 | 0.03547 |
| ENSG00000087274 | ADD1 | 82.49833146 | 137.2742743 | 114.5369611 | 77.70599625 | 69.74275556 | 71.81555 | 0.601067 | 6.542584 | 0.00142 | 0.03566 |
| ENSG00000113580 | NR3C1 | 64.16536891 | 90.2088088 | 60.97241211 | 108.3174493 | 102.6768346 | 104.0269 | -0.55228 | 6.481745 | 0.00143 | 0.03586 |
| ENSG00000068615 | REEP1 | 3.666592509 | 7.283941083 | 5.698356272 | 14.12836296 | 10.97802634 | 11.08917 | -1.15148 | 3.299077 | 0.00145 | 0.03611 |
| ENSG00000138347 | MYPN | 61.41542453 | 77.88213927 | 98.01172788 | 55.10061553 | 49.72400165 | 49.63722 | 0.609846 | 6.050307 | 0.00145 | 0.03611 |
| ENSG00000111540 | RAB5B | 52.70726732 | 81.80426139 | 50.14553519 | 30.61145307 | 38.74597531 | 43.30055 | 0.713638 | 5.658994 | 0.00146 | 0.03627 |
| ENSG00000128591 | FLNC | 618.7374859 | 761.4519948 | 827.9711663 | 583.50139 | 386.8139869 | 536.5044 | 0.549136 | 9.277034 | 0.00146 | 0.03627 |
| ENSG00000119711 | ALDH6A1 | 69.20693361 | 134.4727585 | 79.20715218 | 67.34519675 | 51.01553416 | 46.46889 | 0.77094 | 6.241401 | 0.00147 | 0.0365 |
| ENSG00000214975 | PPIAP29 | 21.08290693 | 18.49000429 | 25.64260322 | 28.25672591 | 39.39174157 | 42.7725 | -0.75197 | 4.913556 | 0.00151 | 0.03711 |
| ENSG00000139644 | TMBIM6 | 32.08268446 | 30.81667381 | 29.63145261 | 35.79185282 | 57.47319671 | 58.08611 | -0.69661 | 5.373325 | 0.00151 | 0.03711 |
| ENSG00000147573 | TRIM55 | 24.29117537 | 28.01515801 | 30.20128824 | 14.59930839 | 18.7272214 | 15.84167 | 0.751296 | 4.519078 | 0.00151 | 0.03711 |
| ENSG00000100324 | TAB1 | 11.91642565 | 10.08545688 | 10.82687692 | 7.064181478 | 2.583065021 | 4.7525 | 1.127863 | 3.1831 | 0.00151 | 0.03716 |
| ENSG00000140718 | FTO | 20.62458286 | 17.36939797 | 16.52523319 | 11.7736358 | 4.520363787 | 10.56111 | 0.979923 | 3.893508 | 0.00153 | 0.03741 |
| ENSG00000130821 | SLC6A8 | 119.1642565 | 112.6209352 | 107.6989335 | 91.36341378 | 69.74275556 | 80.26444 | 0.485784 | 6.61642 | 0.00153 | 0.03743 |
| ENSG00000110958 | PTGES3 | 142.9971079 | 115.9827542 | 157.8444687 | 179.901155 | 196.3129416 | 205.4136 | -0.47921 | 7.386833 | 0.00154 | 0.03754 |
| ENSG00000117592 | PRDX6 | 133.8306266 | 128.8697269 | 141.8890712 | 74.88032366 | 100.7395358 | 107.7233 | 0.518493 | 6.853717 | 0.00154 | 0.03754 |
| ENSG00000171735 | CAMTA1 | 140.7054875 | 118.2239668 | 107.6989335 | 189.791009 | 142.7143424 | 198.0208 | -0.53337 | 7.237021 | 0.00155 | 0.03767 |
| ENSG00000186940 | CHCHD2P9 | 5.499888764 | 5.042728442 | 1.709506882 | 17.89592641 | 7.749195063 | 8.976944 | -1.48385 | 3.180262 | 0.00156 | 0.03788 |
| ENSG00000238193 | RP11-555H23.1 | 591.6963662 | 571.5092234 | 580.0926685 | 471.8873227 | 457.848275 | 425.6128 | 0.363161 | 9.015664 | 0.00157 | 0.03797 |
| ENSG00000173692 | PSMD1 | 186.0795698 | 182.0985271 | 166.9618388 | 118.6782488 | 140.1312774 | 136.7664 | 0.44015 | 7.287288 | 0.00157 | 0.03797 |
| ENSG00000089818 | NECAP1 | 5.0415647 | 8.404547404 | 5.128520645 | 2.354727159 | 1.937298766 | 1.584167 | 1.603712 | 2.361689 | 0.00158 | 0.03812 |
| ENSG00000197063 | MAFG | 19.24961067 | 16.24879165 | 17.66490444 | 29.66956221 | 27.12218272 | 29.04305 | -0.68733 | 4.599942 | 0.0016 | 0.03849 |
| ENSG00000126561 | STAT5A | 17.87463848 | 20.73121693 | 20.51408258 | 12.71552666 | 9.040727573 | 10.56111 | 0.841377 | 4.032504 | 0.00161 | 0.03865 |
| ENSG00000168906 | MAT2A | 180.121357 | 192.183984 | 139.039893 | 133.2775572 | 117.5294585 | 106.1392 | 0.517782 | 7.189536 | 0.00165 | 0.03916 |
| ENSG00000181856 | SLC2A4 | 330.909974 | 322.1743171 | 340.1918694 | 212.8673352 | 282.8456198 | 237.625 | 0.44113 | 8.173508 | 0.00165 | 0.03916 |
| ENSG00000174429 | ABRA | 47.20737856 | 39.22122122 | 41.02816516 | 26.84388961 | 32.28831276 | 22.17833 | 0.663264 | 5.16604 | 0.00165 | 0.03916 |
| ENSG00000149547 | EI24 | 47.66570262 | 50.98758758 | 48.43602831 | 33.43712566 | 30.351014 | 34.85167 | 0.569007 | 5.395474 | 0.00165 | 0.03916 |
| ENSG00000159792 | PSKH1 | 9.624805337 | 18.49000429 | 17.66490444 | 5.651345182 | 7.103428807 | 7.920833 | 1.129575 | 3.587254 | 0.00166 | 0.03917 |
| ENSG00000143365 | RORC | 31.16603633 | 39.78152438 | 22.79342509 | 16.01214468 | 21.31028642 | 15.84167 | 0.830496 | 4.674254 | 0.00166 | 0.03917 |
| ENSG00000141458 | NPC1 | 10.0831294 | 15.12818533 | 11.39671254 | 7.064181478 | 4.520363787 | 4.7525 | 1.117226 | 3.309949 | 0.00166 | 0.03917 |
| ENSG00000142173 | COL6A2 | 45.3740823 | 69.47759187 | 57.55339835 | 35.32090739 | 30.99678025 | 42.7725 | 0.647008 | 5.583544 | 0.00166 | 0.03917 |
| ENSG00000096746 | HNRNPH3 | 79.29006301 | 67.79668239 | 75.21830279 | 97.48570439 | 100.0937696 | 112.4758 | -0.47699 | 6.488208 | 0.00167 | 0.03921 |
| ENSG00000229037 | RP11-272G22.3 | 7.791509082 | 10.64576004 | 10.25704129 | 2.354727159 | 4.520363787 | 4.7525 | 1.308081 | 2.949713 | 0.00167 | 0.03928 |
| ENSG00000116667 | C1orf21 | 22.45787912 | 15.12818533 | 15.95539756 | 31.0823985 | 34.87137778 | 25.34667 | -0.7455 | 4.653359 | 0.00169 | 0.03957 |
| ENSG00000168273 | SMIM4 | 46.29073043 | 46.5051623 | 51.85504207 | 59.81006984 | 76.84618437 | 75.51194 | -0.54762 | 5.914405 | 0.00169 | 0.03957 |
| ENSG00000163171 | CDC42EP3 | 20.1662588 | 20.73121693 | 22.79342509 | 9.889854069 | 16.14415638 | 8.976944 | 0.888488 | 4.122616 | 0.0017 | 0.03959 |
| ENSG00000058272 | PPP1R12A | 80.6650352 | 90.76911196 | 71.79928903 | 104.5498859 | 114.9463934 | 120.9247 | -0.48284 | 6.618099 | 0.00171 | 0.03959 |
| ENSG00000197958 | RPL12 | 100.3729699 | 91.88971828 | 87.75468659 | 112.0850128 | 136.9024461 | 143.103 | -0.47976 | 6.819561 | 0.00171 | 0.03959 |
| ENSG00000173457 | PPP1R14B | 22.45787912 | 19.05030745 | 17.66490444 | 26.37294418 | 41.97480659 | 33.2675 | -0.75414 | 4.788795 | 0.00171 | 0.03959 |
| ENSG00000137449 | CPEB2 | 66.91531329 | 68.91728871 | 52.99471333 | 82.41545057 | 98.80223705 | 90.2975 | -0.516 | 6.278876 | 0.00174 | 0.04014 |
| ENSG00000232362 | ATP5LP2 | 120.9975528 | 115.422451 | 174.3697019 | 177.0754824 | 203.4163704 | 222.8394 | -0.55564 | 7.407983 | 0.00176 | 0.04058 |
| ENSG00000164258 | NDUFS4 | 246.5783462 | 219.0785357 | 267.8227448 | 318.3591119 | 304.1559062 | 369.6389 | -0.43646 | 8.173266 | 0.00177 | 0.04066 |
| ENSG00000107223 | EDF1 | 119.6225806 | 113.7415415 | 116.8163036 | 153.9991562 | 169.1907589 | 147.8555 | -0.42441 | 7.106486 | 0.00178 | 0.04095 |
| ENSG00000220960 | RP1-72A23.1 | 3.666592509 | 1.680909481 | 2.279342509 | 5.18039975 | 7.749195063 | 8.448889 | -1.40766 | 2.549525 | 0.00178 | 0.04095 |
| ENSG00000091073 | DTX2 | 1.374972191 | 1.680909481 | 0.569835627 | 4.238508887 | 4.520363787 | 5.280555 | -1.87783 | 1.996002 | 0.00183 | 0.04181 |
| ENSG00000241932 | RP11-14K2.1 | 17.87463848 | 13.44727585 | 16.52523319 | 25.90199875 | 21.95605268 | 35.37972 | -0.79724 | 4.522529 | 0.00183 | 0.04181 |
| ENSG00000175048 | ZDHHC14 | 10.54145346 | 8.964850564 | 7.407863154 | 2.825672591 | 5.166130042 | 3.168333 | 1.312137 | 2.894302 | 0.00183 | 0.04182 |
| ENSG00000131236 | CAP1 | 42.62413792 | 45.94485914 | 47.29635706 | 24.96010789 | 32.28831276 | 32.21139 | 0.6091 | 5.268768 | 0.00185 | 0.04212 |
| ENSG00000164068 | RNF123 | 87.08157209 | 104.776691 | 128.7828517 | 79.58977798 | 59.41049548 | 75.51194 | 0.568532 | 6.496545 | 0.00187 | 0.04252 |
| ENSG00000142208 | AKT1 | 164.0800148 | 149.0406406 | 151.0064412 | 109.2593402 | 112.3633284 | 123.565 | 0.428613 | 7.088117 | 0.00189 | 0.0428 |
| ENSG00000251306 | NDUFB4P2 | 81.58168333 | 95.25153724 | 113.3972898 | 163.4180648 | 109.1344971 | 161.0569 | -0.58717 | 6.928743 | 0.00191 | 0.04311 |
| ENSG00000019144 | PHLDB1 | 130.6223581 | 131.6712427 | 124.2241667 | 104.5498859 | 83.94961318 | 93.46583 | 0.449276 | 6.815895 | 0.00191 | 0.04311 |
| ENSG00000144476 | ACKR3 | 118.7059325 | 115.422451 | 107.6989335 | 66.87425132 | 96.21917203 | 77.62416 | 0.516313 | 6.615761 | 0.00194 | 0.04363 |
| ENSG00000214263 | RPSAP53 | 32.54100852 | 28.57546117 | 30.77112387 | 54.62967009 | 36.16291029 | 53.33361 | -0.65778 | 5.343036 | 0.00195 | 0.04383 |
| ENSG00000147471 | PROSC | 10.0831294 | 15.12818533 | 10.82687692 | 1.412836296 | 7.103428807 | 5.808611 | 1.373025 | 3.223763 | 0.00195 | 0.04383 |
| ENSG00000121644 | DESI2 | 12.37474972 | 7.844244243 | 4.558685018 | 20.721599 | 12.9153251 | 19.01 | -1.06521 | 3.824317 | 0.00196 | 0.04395 |
| ENSG00000134240 | HMGCS2 | 12.83307378 | 10.08545688 | 68.38027526 | 10.83174493 | 9.686493828 | 3.696389 | 1.901704 | 4.321845 | 0.00198 | 0.04433 |
| ENSG00000113328 | CCNG1 | 345.1180199 | 459.4485914 | 352.1584176 | 277.8578048 | 264.1183984 | 299.9355 | 0.456739 | 8.384519 | 0.002 | 0.04468 |
| ENSG00000166848 | TERF2IP | 9.166481273 | 10.08545688 | 10.25704129 | 2.354727159 | 5.166130042 | 5.280555 | 1.226932 | 3.013511 | 0.00201 | 0.04483 |
| ENSG00000266472 | MRPS21 | 26.12447163 | 29.13576433 | 32.48063075 | 37.20468912 | 55.53589795 | 45.94083 | -0.65249 | 5.266151 | 0.00202 | 0.04486 |
| ENSG00000179820 | MYADM | 8.249833146 | 10.08545688 | 10.25704129 | 5.18039975 | 2.583065021 | 4.224444 | 1.186617 | 2.977375 | 0.00204 | 0.04532 |
| ENSG00000112339 | HBS1L | 57.29050796 | 56.03031602 | 60.97241211 | 41.91414343 | 37.4544428 | 41.18833 | 0.525063 | 5.652228 | 0.00205 | 0.04541 |
| ENSG00000137824 | RMDN3 | 9.166481273 | 10.64576004 | 12.5363838 | 2.825672591 | 7.103428807 | 4.224444 | 1.228295 | 3.123218 | 0.00206 | 0.04545 |
| ENSG00000125037 | EMC3 | 60.4987764 | 55.47001286 | 41.59800079 | 32.96618023 | 38.74597531 | 31.15528 | 0.627859 | 5.478101 | 0.00206 | 0.04545 |
| ENSG00000144579 | CTDSP1 | 85.7065999 | 95.8118404 | 109.978276 | 69.22897848 | 69.09698931 | 69.17527 | 0.485867 | 6.395458 | 0.00207 | 0.04545 |
| ENSG00000122406 | RPL5 | 398.2836113 | 305.9255255 | 347.5997326 | 491.6670308 | 461.7228725 | 451.4875 | -0.41641 | 8.681551 | 0.00207 | 0.04546 |
| ENSG00000176485 | PLA2G16 | 16.95799035 | 14.56788217 | 22.79342509 | 31.0823985 | 34.87137778 | 25.34667 | -0.74568 | 4.654307 | 0.00207 | 0.04546 |
| ENSG00000183963 | SMTN | 152.6219132 | 124.9476047 | 132.7717011 | 185.0815547 | 175.6484214 | 191.6842 | -0.4268 | 7.3362 | 0.00208 | 0.04546 |
| ENSG00000177889 | UBE2N | 70.12358174 | 76.20122979 | 79.20715218 | 44.73981602 | 56.1816642 | 56.50194 | 0.522624 | 6.018048 | 0.00208 | 0.04548 |
| ENSG00000204396 | VWA7 | 7.333185018 | 6.163334763 | 17.66490444 | 4.709454318 | 1.937298766 | 3.168333 | 1.595133 | 2.980087 | 0.00209 | 0.04553 |
| ENSG00000198881 | ASB12 | 5.499888764 | 3.922122122 | 2.279342509 | 1.412836296 | 0 | 0.528056 | 2.376277 | 1.783639 | 0.00211 | 0.04579 |
| ENSG00000116260 | QSOX1 | 8.249833146 | 7.283941083 | 7.977698781 | 2.825672591 | 1.29153251 | 4.7525 | 1.348474 | 2.709482 | 0.00211 | 0.04579 |
| ENSG00000115307 | AUP1 | 108.6228031 | 104.2163878 | 102.5704129 | 135.1613389 | 131.0905498 | 170.5619 | -0.47002 | 6.982464 | 0.00211 | 0.04579 |
| ENSG00000139197 | PEX5 | 36.20760103 | 24.09303589 | 38.74882265 | 16.48309011 | 25.18488395 | 14.78555 | 0.832112 | 4.752234 | 0.00213 | 0.046 |
| ENSG00000081189 | MEF2C | 84.33162771 | 80.68365507 | 79.77698781 | 109.2593402 | 100.0937696 | 133.598 | -0.48774 | 6.630424 | 0.00213 | 0.046 |

| | | | | | | | | | | | |
|---|---|---|---|---|---|---|---|---|---|---|---|
| ENSG00000146677 | AC004453.8 | 216.328958 | 133.3521521 | 190.3250995 | 231.234207 | 261.5353334 | 306.8003 | -0.56347 | 7.809407 | 0.00214 | 0.04614 |
| ENSG00000126432 | PRDX5 | 188.3711902 | 145.6788217 | 147.5874274 | 236.4146068 | 211.8113317 | 211.7503 | -0.45207 | 7.580959 | 0.00215 | 0.04625 |
| ENSG00000170791 | CHCHD7 | 15.1246941 | 12.88697269 | 19.37441132 | 22.1344353 | 32.93407902 | 26.93083 | -0.77703 | 4.485252 | 0.00216 | 0.04636 |
| ENSG00000120333 | MRPS14 | 28.87441601 | 28.01515801 | 32.48063075 | 41.91414343 | 51.66130042 | 42.24444 | -0.59609 | 5.263194 | 0.00217 | 0.04651 |
| ENSG00000049283 | EPN3 | 12.37474972 | 10.08545688 | 11.39671254 | 5.18039975 | 5.811896297 | 5.808611 | 1.016922 | 3.25609 | 0.00218 | 0.04666 |
| ENSG00000100644 | HIF1A | 37.12424916 | 36.41970542 | 41.02816516 | 53.2168338 | 57.47319671 | 54.91778 | -0.52966 | 5.574337 | 0.00221 | 0.04709 |
| ENSG00000157184 | CPT2 | 60.04045234 | 58.27152866 | 55.27405584 | 30.61145307 | 43.91210536 | 42.7725 | 0.578669 | 5.629583 | 0.00221 | 0.04709 |
| ENSG00000254502 | FNTAL1 | 17.41631442 | 14.00757901 | 14.24589068 | 9.889854069 | 7.103428807 | 6.864722 | 0.921887 | 3.676712 | 0.00222 | 0.0471 |
| ENSG00000204899 | MZT1 | 10.54145346 | 10.08545688 | 8.547534408 | 4.238508887 | 5.166130042 | 4.224444 | 1.109273 | 3.04099 | 0.00222 | 0.0471 |
| ENSG00000171992 | SYNPO | 42.62413792 | 39.78152438 | 68.95011089 | 34.37901652 | 20.66452017 | 32.21139 | 0.774835 | 5.355175 | 0.00222 | 0.0471 |
| ENSG00000122679 | RAMP3 | 2.749944382 | 2.801515801 | 3.98884939 | 0 | 1.29153251 | 0.528056 | 2.405516 | 1.549346 | 0.00223 | 0.04718 |
| ENSG00000197358 | BNIP3P1 | 43.08246198 | 57.15092234 | 63.82159025 | 26.84388961 | 46.49517038 | 22.70639 | 0.784877 | 5.462181 | 0.00224 | 0.04718 |
| ENSG00000196154 | S100A4 | 11.91642565 | 8.404547404 | 11.96654817 | 17.42498098 | 20.01875391 | 20.06611 | -0.81516 | 3.997487 | 0.00225 | 0.04732 |
| ENSG00000112972 | HMGCS1 | 23.83285131 | 39.78152438 | 27.92194573 | 17.89592641 | 20.66452017 | 13.20139 | 0.821212 | 4.633917 | 0.00227 | 0.04777 |
| ENSG00000116750 | UCHL5 | 25.2078235 | 17.92970113 | 21.08391821 | 32.4952348 | 33.57984527 | 35.90778 | -0.65355 | 4.846604 | 0.00229 | 0.04809 |
| ENSG00000234152 | ELOBP1 | 3.208268446 | 4.482425282 | 2.849178136 | 8.006072341 | 10.33226008 | 7.392777 | -1.26011 | 2.811037 | 0.00231 | 0.04829 |
| ENSG00000131828 | PDHA1 | 395.0753429 | 446.0013155 | 352.7282532 | 297.6375129 | 323.5288939 | 266.14 | 0.429188 | 8.442572 | 0.00232 | 0.04859 |
| ENSG00000168334 | XIRP1 | 27.04111976 | 67.79668239 | 43.30750767 | 31.0823985 | 12.9153251 | 23.23444 | 1.013546 | 5.142104 | 0.00234 | 0.0488 |
| ENSG00000196735 | HLA-DQA1 | 19.70793474 | 19.61061061 | 15.95539756 | 32.02428937 | 27.76794897 | 28.515 | -0.67471 | 4.645685 | 0.00234 | 0.04881 |
| ENSG00000248746 | ACTN3 | 25.66614756 | 28.57546117 | 27.92194573 | 16.48309011 | 20.01875391 | 13.20139 | 0.733267 | 4.522845 | 0.00235 | 0.04888 |
| ENSG00000188612 | SUMO2 | 176.9130886 | 172.5733734 | 120.2353173 | 185.5525001 | 248.6200083 | 257.163 | -0.55358 | 7.602867 | 0.00236 | 0.04896 |
| ENSG00000106299 | WASL | 21.99955505 | 21.85182325 | 33.05046638 | 44.73981602 | 39.39174157 | 37.49194 | -0.67249 | 5.091107 | 0.00237 | 0.04903 |
| ENSG00000111196 | MAGOHB | 10.0831294 | 5.603031602 | 5.698356272 | 13.65741752 | 21.31028642 | 11.61722 | -1.06851 | 3.616405 | 0.00239 | 0.04946 |
| ENSG00000141562 | NARF | 11.45810159 | 15.68848849 | 8.547534408 | 3.296618023 | 8.394961318 | 3.696389 | 1.259524 | 3.248421 | 0.0024 | 0.04951 |
| ENSG00000108829 | LRRC59 | 16.49966629 | 15.12818533 | 21.08391821 | 9.418908637 | 12.26955885 | 5.280555 | 0.979562 | 3.836825 | 0.00243 | 0.04998 |

### Supple Table 6 Cardiac DEs for HFD_Spp_3wks vs HFD_CTL

| Ensembl ID | Gene symbol | HFD_Spp_3wk_1 | HFD_Spp_3wk_2 | HFD_Spp_3wk_3 | HFD-CTL-1 | HFD-CTL-2 | HFD-CTL-3 | logFC | logCPM | PValue | FDR |
|---|---|---|---|---|---|---|---|---|---|---|---|
| ENSG00000133112 | TPT1 | 407.0493785 | 374.4958403 | 350.8322453 | 688.7116624 | 660.745259 | 688.788136 | -0.84802723 | 9.047563735 | 1.74E-10 | 1.98E-06 |
| ENSG00000154518 | ATP5G3 | 4607.03606 | 4273.922551 | 4471.563339 | 6898.869383 | 7460.4378 | 7476.17761 | -0.7095417 | 12.51799352 | 9.72E-09 | 5.35E-05 |
| ENSG00000110717 | NDUFS8 | 143.0444795 | 140.5481991 | 147.5559149 | 251.9791475 | 241.254772 | 250.182925 | -0.78664641 | 7.618527255 | 1.41E-08 | 5.35E-05 |
| ENSG00000112695 | COX7A2 | 17.06495545 | 14.81818073 | 14.44603363 | 35.72838658 | 33.7374747 | 46.5821763 | -1.32550506 | 4.800494362 | 2.99E-08 | 7.59E-05 |
| ENSG00000231739 | GAPDHP59 | 144.0483004 | 163.8980596 | 164.0656677 | 258.5606924 | 267.990129 | 299.905472 | -0.8075116 | 7.762478344 | 3.34E-08 | 7.59E-05 |
| ENSG00000172115 | CYCS | 78.79994133 | 76.33608255 | 94.41514838 | 155.1364154 | 145.134797 | 150.214434 | -0.85461396 | 6.875567645 | 6.83E-08 | 0.000111 |
| ENSG00000217027 | TPT1P4 | 64.74644861 | 61.06886604 | 76.35760634 | 108.5954908 | 150.227246 | 141.84011 | -0.98412765 | 6.657578455 | 6.84E-08 | 0.000111 |
| ENSG00000248448 | COX5BP1 | 531.5231711 | 500.2258586 | 459.1774976 | 859.3617194 | 770.232913 | 816.496574 | -0.71451691 | 9.359733142 | 9.44E-08 | 0.000132 |
| ENSG00000214318 | ATP5G1P6 | 125.4776136 | 140.9972348 | 126.4027943 | 230.8241817 | 206.244185 | 274.782501 | -0.85779314 | 7.531112592 | 1.05E-07 | 0.000132 |
| ENSG00000276293 | PIP4K2B | 21.58214954 | 23.34986055 | 21.66905045 | 8.461986296 | 6.36556127 | 9.42111432 | 1.43487301 | 4.036802129 | 1.53E-07 | 0.000174 |
| ENSG00000179094 | PER1 | 57.71970225 | 68.25343852 | 61.91157271 | 34.78816588 | 28.6450257 | 33.4972953 | 0.9472974 | 5.609493779 | 4.51E-07 | 0.000466 |
| ENSG00000136859 | ANGPTL2 | 66.25217997 | 83.96969081 | 63.45936202 | 36.66860728 | 32.4643625 | 39.254643 | 0.97632847 | 5.783438559 | 6.07E-07 | 0.000574 |
| ENSG00000162413 | KLHL21 | 40.15283635 | 41.31129173 | 34.05136499 | 19.74463469 | 15.9139032 | 18.3188334 | 1.08700754 | 4.885974395 | 8.20E-07 | 0.000717 |
| ENSG00000182606 | TRAK1 | 178.1782113 | 193.9834568 | 196.5692433 | 123.1689116 | 107.577985 | 111.483186 | 0.72994804 | 7.25890199 | 9.20E-07 | 0.000746 |
| ENSG00000135821 | GLUL | 221.8444208 | 258.6446091 | 235.7799061 | 135.3917807 | 146.407909 | 154.401596 | 0.71681261 | 7.595248337 | 1.24E-06 | 0.000867 |
| ENSG00000185641 | CTD-2287O16.1 | 182.1934949 | 212.8429596 | 175.4161227 | 298.0499618 | 300.454492 | 396.733592 | -0.80218209 | 8.032036463 | 1.43E-06 | 0.000867 |
| ENSG00000102858 | MGRN1 | 84.82286678 | 99.23690732 | 105.2496736 | 55.94313162 | 49.6513779 | 56.5266859 | 0.83093545 | 6.258430109 | 1.44E-06 | 0.000867 |
| ENSG00000168079 | SCARA5 | 6.524835906 | 8.980715594 | 10.31859545 | 1.410331049 | 1.90966838 | 1.57018572 | 2.37623531 | 2.646095812 | 1.44E-06 | 0.000867 |
| ENSG00000109906 | ZBTB16 | 54.70823952 | 135.6088055 | 148.0718447 | 46.54092463 | 40.7395921 | 37.161062 | 1.44078281 | 6.297463684 | 1.45E-06 | 0.000867 |
| ENSG00000115593 | SMYD1 | 275.5488394 | 254.6032871 | 289.9525322 | 179.1120433 | 174.416379 | 149.691039 | 0.70398992 | 7.793063233 | 1.76E-06 | 0.000998 |
| ENSG00000173915 | USMG5 | 202.7718236 | 205.2093513 | 182.6391395 | 283.9466513 | 336.101635 | 333.926163 | -0.68978938 | 8.011821542 | 2.13E-06 | 0.001143 |
| ENSG00000213862 | CTD-2270N23.1 | 148.5654945 | 145.9366284 | 130.5302325 | 217.1909816 | 227.250537 | 239.191625 | -0.68448242 | 7.535616954 | 2.21E-06 | 0.001143 |
| ENSG00000131389 | SLC6A6 | 59.72734407 | 64.2121165 | 68.61865975 | 40.42949008 | 28.6450257 | 32.9739001 | 0.90212965 | 5.656717124 | 2.50E-06 | 0.001235 |
| ENSG00000266563 | EIF1P5 | 43.66620953 | 44.45454219 | 43.33810089 | 64.40511792 | 86.5716332 | 108.342815 | -0.9760091 | 6.039314834 | 3.04E-06 | 0.001438 |
| ENSG00000229822 | RP5-1174J21.2 | 47.68149316 | 49.39393577 | 53.65669635 | 80.38886981 | 85.298521 | 99.4450956 | -0.81432933 | 6.1307776 | 3.22E-06 | 0.001463 |
| ENSG00000146535 | GNA12 | 65.24835906 | 54.33332935 | 72.23016816 | 38.07893833 | 31.8278063 | 30.8803191 | 0.91819853 | 5.64404465 | 3.50E-06 | 0.001528 |
| ENSG00000232734 | ATP5G1P7 | 77.29420997 | 96.54269264 | 82.03283384 | 138.2124428 | 133.04023 | 167.486477 | -0.77668542 | 6.865621444 | 4.32E-06 | 0.001818 |
| ENSG00000237528 | RP11-234P3.4 | 73.78083679 | 88.90908439 | 73.77795747 | 116.5873667 | 147.044465 | 141.84011 | -0.77213011 | 6.750294831 | 4.86E-06 | 0.001973 |
| ENSG00000224004 | ATP5C1P1 | 433.6506326 | 462.5068531 | 458.145638 | 674.1382416 | 656.289366 | 687.741345 | -0.57550785 | 9.136730829 | 5.86E-06 | 0.002297 |
| ENSG00000232362 | ATP5LP2 | 137.5234645 | 111.3608734 | 91.31956974 | 176.7614915 | 200.51518 | 220.872791 | -0.81271595 | 7.29533354 | 6.21E-06 | 0.002323 |
| ENSG00000227008 | RP3-417G15.1 | 42.16047816 | 32.33057614 | 34.05136499 | 62.05456617 | 72.5673984 | 62.8074288 | -0.85974622 | 5.691337877 | 6.34E-06 | 0.002323 |
| ENSG00000242058 | RPS4XP19 | 47.17958271 | 39.51514862 | 39.21066271 | 70.98666282 | 70.65773 | 79.5560764 | -0.8138397 | 5.874045549 | 7.05E-06 | 0.002501 |
| ENSG00000240371 | RPS4XP13 | 96.36680723 | 90.7052275 | 94.41514838 | 166.4190638 | 131.130562 | 241.808601 | -0.94030134 | 7.105516854 | 8.04E-06 | 0.002768 |
| ENSG00000074416 | MGLL | 16.06113454 | 13.47107339 | 14.44603363 | 5.171213848 | 5.72900514 | 5.2339524 | 1.43984616 | 3.4809878 | 8.53E-06 | 0.002851 |
| ENSG00000161011 | SQSTM1 | 235.3960031 | 219.1294605 | 198.1170327 | 130.6906772 | 150.227246 | 128.231834 | 0.67635387 | 7.477133733 | 9.45E-06 | 0.003066 |
| ENSG00000261679 | ATP5HP1 | 26.60125408 | 22.45178899 | 22.18498022 | 47.01103498 | 45.195485 | 40.8248287 | -0.90109083 | 5.123499398 | 9.76E-06 | 0.003078 |
| ENSG00000104812 | GYS1 | 93.85725496 | 79.92836879 | 89.77178042 | 57.82357302 | 47.7417095 | 48.1523621 | 0.77009168 | 6.146029232 | 1.03E-05 | 0.003173 |
| ENSG00000147123 | NDUFB11 | 115.4394045 | 115.4021954 | 102.6700247 | 180.9924847 | 159.775588 | 190.515867 | -0.67268629 | 7.180170086 | 1.17E-05 | 0.003417 |
| ENSG00000251306 | NDUFB4P2 | 85.82668769 | 84.86776237 | 67.5868002 | 163.1282914 | 107.577985 | 159.635548 | -0.8556126 | 6.813450743 | 1.19E-05 | 0.003417 |
| ENSG00000242262 | RP11-100N21.1 | 269.0240035 | 246.5206431 | 249.7100099 | 361.514859 | 382.570232 | 427.613911 | -0.61450605 | 8.338002005 | 1.20E-05 | 0.003417 |
| ENSG00000218682 | AC010150.1 | 121.46233 | 97.88979998 | 119.1797775 | 168.7696156 | 183.328164 | 191.562658 | -0.68415747 | 7.206843215 | 1.30E-05 | 0.003595 |
| ENSG00000114023 | FAM162A | 48.68531407 | 49.39393577 | 50.04518794 | 80.85898016 | 78.2964036 | 86.8836098 | -0.73268559 | 6.055842668 | 1.44E-05 | 0.003884 |
| ENSG00000129084 | PSMA1 | 10.54011954 | 9.878787184 | 13.93010386 | 19.27452434 | 34.3740308 | 29.3101334 | -1.25822672 | 4.332016987 | 1.49E-05 | 0.003886 |
| ENSG00000133639 | BTG1 | 71.77319497 | 57.92561558 | 63.45936202 | 95.90251136 | 119.672552 | 109.389605 | -0.74766362 | 6.442686494 | 1.51E-05 | 0.003886 |
| ENSG00000233055 | RP11-490D19.8 | 274.5450185 | 291.8732568 | 263.124184 | 434.3819632 | 402.303472 | 401.444149 | -0.5777041 | 8.432436453 | 1.74E-05 | 0.004395 |
| ENSG00000100994 | PYGB | 599.7829929 | 732.8263925 | 578.357275 | 441.4336185 | 396.574467 | 395.686801 | 0.63110859 | 9.037560442 | 1.98E-05 | 0.004888 |
| ENSG00000242706 | RPS27AP9 | 53.70441861 | 53.43525779 | 42.82217112 | 74.74754562 | 87.2081893 | 96.8281194 | -0.78212113 | 6.105922506 | 2.05E-05 | 0.004949 |
| ENSG00000234750 | CTD-2666L21.2 | 218.8329581 | 253.7052155 | 227.0090999 | 417.4579906 | 301.091048 | 440.698792 | -0.72980053 | 8.279923626 | 2.11E-05 | 0.004994 |
| ENSG00000112964 | GHR | 14.05349272 | 11.22589449 | 11.86638477 | 18.8041399 | 38.1933676 | 33.4972953 | -1.26776047 | 4.447545785 | 2.37E-05 | 0.005493 |
| ENSG00000146729 | GBAS | 270.0278244 | 289.1790421 | 287.3728833 | 158.4271879 | 211.97319 | 127.708439 | 0.76893623 | 7.815447368 | 2.88E-05 | 0.006553 |
| ENSG00000178464 | RPL10P16 | 31.62035862 | 32.33057614 | 37.14694362 | 55.00291093 | 56.6534953 | 61.2372431 | -0.77456743 | 5.538108798 | 3.21E-05 | 0.007139 |
| ENSG00000136842 | TMOD1 | 165.1285395 | 147.7327715 | 167.1612463 | 101.0737252 | 111.397322 | 96.3047241 | 0.63792417 | 7.051148733 | 3.31E-05 | 0.00724 |
| ENSG00000234031 | RPS3AP44 | 6.524835906 | 8.082644035 | 9.802665678 | 16.45386224 | 27.3719134 | 17.7954382 | -1.31152415 | 3.906293509 | 3.47E-05 | 0.007352 |
| ENSG00000236929 | RP11-699A7.1 | 104.3973745 | 87.11294127 | 75.84167656 | 114.7069253 | 185.874389 | 194.703029 | -0.88605732 | 6.995257611 | 3.50E-05 | 0.007352 |
| ENSG00000251939 | RNU6-1278P | 1.505731363 | 1.796143119 | 3.611508408 | 6.581544897 | 8.91178577 | 10.9913 | -1.91579077 | 2.682083974 | 3.57E-05 | 0.007367 |
| ENSG00000277443 | MARCKS | 99.88018041 | 72.29476054 | 84.09655293 | 138.2124428 | 124.128445 | 182.664939 | -0.79915869 | 6.879045777 | 3.69E-05 | 0.007435 |
| ENSG00000140945 | CDH13 | 97.8725386 | 78.13222567 | 108.861182 | 58.76379372 | 50.9244901 | 58.6202669 | 0.75177242 | 6.262050441 | 3.73E-05 | 0.007435 |
| ENSG00000196776 | CD47 | 10.54011954 | 13.02203761 | 12.38231454 | 21.15496574 | 29.9181379 | 25.6463667 | -1.0735101 | 4.28589566 | 4.07E-05 | 0.007975 |
| ENSG00000138326 | RPS24 | 327.2456162 | 316.1211889 | 297.175549 | 442.3738392 | 444.316176 | 488.327759 | -0.54778267 | 8.595261121 | 4.48E-05 | 0.008627 |
| ENSG00000143420 | ENSA | 33.12608999 | 61.9669376 | 51.07704748 | 29.61695204 | 20.369796 | 18.3188334 | 1.08836938 | 5.216494181 | 4.89E-05 | 0.009263 |
| ENSG00000243859 | RPL5P17 | 16.06113454 | 13.02203761 | 10.83452522 | 22.09518644 | 28.0084696 | 38.7312477 | -1.14672561 | 4.474718415 | 5.03E-05 | 0.009363 |
| ENSG00000101558 | VAPA | 184.7030472 | 142.793378 | 144.9762661 | 253.3894785 | 225.340869 | 258.033853 | -0.64261954 | 7.661002166 | 5.15E-05 | 0.009429 |
| ENSG00000143933 | CALM2 | 230.3768985 | 251.0110009 | 225.4613106 | 325.7864724 | 364.746661 | 350.151415 | -0.55639969 | 8.189873406 | 5.43E-05 | 0.009782 |
| ENSG00000237550 | RPL9P9 | 72.77701588 | 107.3195514 | 98.54258656 | 129.2803462 | 183.328164 | 164.869501 | -0.7720064 | 6.984097012 | 6.03E-05 | 0.010667 |
| ENSG00000243256 | RPL30P14 | 42.16047816 | 41.76032751 | 35.08322453 | 60.17412477 | 63.0190565 | 81.6496574 | -0.78120529 | 5.776284204 | 6.10E-05 | 0.010667 |
| ENSG00000149577 | SIDT2 | 42.66238862 | 46.69972189 | 55.20448566 | 20.21474504 | 29.2815818 | 29.3101334 | 0.88437471 | 5.261350453 | 6.21E-05 | 0.010692 |
| ENSG00000161533 | ACOX1 | 55.21014998 | 65.55922384 | 72.23016816 | 40.89960043 | 39.4664798 | 31.4037144 | 0.78710185 | 5.700944652 | 6.38E-05 | 0.010817 |
| ENSG00000169047 | IRS1 | 24.59361226 | 26.493111 | 17.02568249 | 45.60070393 | 40.103036 | 39.7780382 | -0.87654749 | 5.052207267 | 6.65E-05 | 0.010953 |
| ENSG00000231006 | RPL7P32 | 75.28656815 | 77.68318989 | 63.9752918 | 104.834608 | 113.306991 | 125.614858 | -0.66102636 | 6.558526855 | 6.65E-05 | 0.010953 |
| ENSG00000225224 | RPS27AP12 | 1.003820909 | 1.347107339 | 1.031859545 | 4.701103498 | 5.09244901 | 6.28074288 | -2.17339532 | 2.062907876 | 7.38E-05 | 0.01194 |
| ENSG00000116688 | MFN2 | 512.9524843 | 533.9035421 | 557.7200841 | 415.5775492 | 287.086813 | 326.075234 | 0.63900279 | 8.782390518 | 7.46E-05 | 0.01194 |
| ENSG00000229758 | DYNLT3P2 | 4.015283635 | 1.796143119 | 2.579648863 | 0 | 0 | 0 | 5.47425341 | 1.299288607 | 7.87E-05 | 0.012422 |
| ENSG00000063660 | GPC1 | 260.9934363 | 337.6749064 | 249.7100099 | 157.0168568 | 166.777705 | 204.647539 | 0.68373968 | 7.850775098 | 8.19E-05 | 0.012685 |
| ENSG00000119715 | ESRRB | 17.5668659 | 19.75757431 | 13.41417409 | 7.051655247 | 3.81933676 | 9.42111432 | 1.29004168 | 3.713971981 | 8.26E-05 | 0.012685 |
| ENSG00000140479 | PCSK6 | 28.6088959 | 34.57575504 | 25.79648863 | 13.16308979 | 13.3676787 | 18.3188334 | 0.98797607 | 4.558794824 | 8.87E-05 | 0.013431 |
| ENSG00000234782 | TPT1P9 | 138.0253749 | 131.1184477 | 128.4665134 | 204.4980022 | 222.158088 | 176.384196 | -0.59912843 | 7.387963896 | 9.18E-05 | 0.013714 |
| ENSG00000235089 | RP3-445O10.1 | 47.17958271 | 54.33332935 | 49.52925816 | 98.25306311 | 68.7480617 | 83.7432384 | -0.73600425 | 6.087101639 | 9.66E-05 | 0.014021 |
| ENSG00000232901 | CYCSP10 | 132.5043599 | 125.7300183 | 147.0399852 | 188.9843606 | 205.607629 | 209.881491 | -0.57675378 | 7.401159508 | 9.93E-05 | 0.014021 |
| ENSG00000117479 | SLC19A2 | 2.509552272 | 4.041322018 | 1.547789318 | 0 | 0 | 0 | 5.46047406 | 1.296182314 | 1.00E-04 | 0.014021 |

| | | | | | | | | | | | |
|---|---|---|---|---|---|---|---|---|---|---|---|
| ENSG00000187446 | CHP1 | 27.60507499 | 30.08539724 | 33.01950544 | 19.74463469 | 14.0042348 | 12.5614858 | 0.95136329 | 4.590295049 | 1.00E-04 | 0.014021 |
| ENSG00000084234 | APLP2 | 418.5933189 | 428.3801339 | 407.0685905 | 308.3923895 | 270.536354 | 293.62473 | 0.52195673 | 8.474650141 | 0.0001 | 0.014021 |
| ENSG00000004779 | NDUFAB1 | 512.9524843 | 414.0109889 | 446.2792533 | 593.7493718 | 764.503908 | 726.472593 | -0.60171198 | 9.172212492 | 0.000101 | 0.014021 |
| ENSG00000090372 | STRN4 | 60.73116498 | 64.66115228 | 69.13458952 | 45.60070393 | 33.7374747 | 37.161062 | 0.72812618 | 5.731709101 | 0.000104 | 0.01424 |
| ENSG00000126561 | STAT5A | 23.0878809 | 21.10468165 | 21.15312067 | 12.69297944 | 8.91178577 | 10.4679048 | 1.002484 | 4.125847287 | 0.000107 | 0.014499 |
| ENSG00000267590 | NDUFA3P1 | 79.80376224 | 102.8291936 | 66.03901089 | 165.0087328 | 112.033878 | 152.308015 | -0.78829894 | 6.833929176 | 0.000112 | 0.014947 |
| ENSG00000240583 | AQP1 | 5.019104543 | 8.082644035 | 6.191157271 | 0.9402207 | 1.27311225 | 2.09358096 | 2.13343955 | 2.343243516 | 0.000115 | 0.014947 |
| ENSG00000115541 | HSPE1 | 13.55158227 | 17.51239541 | 15.47789318 | 25.38595889 | 38.1933676 | 28.7867382 | -0.96742646 | 4.57514106 | 0.000116 | 0.014947 |
| ENSG00000198053 | SIRPA | 11.54394045 | 11.22589449 | 9.286735906 | 23.97562784 | 21.0063522 | 19.8890191 | -1.01451233 | 4.093003042 | 0.000117 | 0.014947 |
| ENSG00000078804 | TP53INP2 | 282.0736753 | 278.8512192 | 294.5959001 | 387.8410386 | 401.666916 | 433.371259 | -0.51542891 | 8.43962748 | 0.000117 | 0.014947 |
| ENSG00000115268 | RPS15 | 94.86107587 | 143.6914495 | 100.0903759 | 157.4869672 | 197.332399 | 213.545258 | -0.74292171 | 7.246911639 | 0.00013 | 0.016383 |
| ENSG00000231500 | RPS18 | 260.4915258 | 294.1184357 | 263.6401138 | 362.92519 | 390.208906 | 488.851154 | -0.60145245 | 8.427030594 | 0.000136 | 0.016946 |
| ENSG00000138442 | WDR12 | 25.09552272 | 16.16528807 | 25.28055885 | 35.25827623 | 40.7395921 | 48.1523621 | -0.90380112 | 5.023001712 | 0.000141 | 0.017451 |
| ENSG00000162889 | MAPKAPK2 | 211.3043013 | 205.2093513 | 223.9135213 | 139.1526635 | 160.412144 | 133.989181 | 0.56507369 | 7.492843882 | 0.000144 | 0.01757 |
| ENSG00000213700 | RPL17P50 | 10.03820909 | 14.81818073 | 10.31859545 | 21.15496574 | 24.8256889 | 24.5995763 | -0.98399189 | 4.20738691 | 0.000146 | 0.017652 |
| ENSG00000213468 | FIRRE | 18.57068681 | 18.41046697 | 17.02568249 | 41.36971078 | 25.4622451 | 36.1142715 | -0.94161924 | 4.764951118 | 0.000153 | 0.017904 |
| ENSG00000244734 | HBB | 334.2723626 | 46.69972109 | 116.6001286 | 498.3169708 | 658.835591 | 532.292959 | -1.76369915 | 8.509841644 | 0.000153 | 0.017904 |
| ENSG00000254373 | RP11-34P1.2 | 120.9604195 | 132.465555 | 119.1797775 | 187.5740296 | 162.321812 | 243.378787 | -0.67198698 | 7.339109715 | 0.000153 | 0.017904 |
| ENSG00000271423 | CYCSP1 | 76.79229951 | 83.52065503 | 82.54876361 | 109.0656012 | 122.218776 | 149.691039 | -0.64807559 | 6.71091896 | 0.000158 | 0.018111 |
| ENSG00000039560 | RAI14 | 0.501910454 | 0.898071559 | 0.515929773 | 5.171213848 | 3.81933676 | 3.66376668 | -2.60443022 | 1.767590822 | 0.000159 | 0.018111 |
| ENSG00000184752 | NDUFA12 | 90.84579224 | 114.9531596 | 89.25585065 | 131.1607876 | 168.050817 | 168.009872 | -0.65799483 | 6.997252326 | 0.000159 | 0.018111 |
| ENSG00000163947 | ARHGEF3 | 11.04203 | 11.67493027 | 14.44603363 | 6.111434547 | 3.18278063 | 5.2339524 | 1.30289398 | 3.292612205 | 0.000162 | 0.018246 |
| ENSG00000167283 | ATP5L | 538.5499175 | 302.2010798 | 576.8094857 | 706.5758557 | 922.369827 | 886.108141 | -0.82741647 | 9.357173433 | 0.000171 | 0.018816 |
| ENSG00000233406 | RP11-296A18.5 | 57.71970225 | 74.09090365 | 57.78413453 | 98.72317346 | 92.9371945 | 106.249234 | -0.64950641 | 6.360385194 | 0.000172 | 0.018816 |
| ENSG00000224631 | RPS27AP16 | 279.5641231 | 280.6473623 | 245.5825717 | 404.7650112 | 350.742426 | 417.669401 | -0.54252724 | 8.369680357 | 0.000172 | 0.018816 |
| ENSG00000168028 | RPSA | 38.14519453 | 47.59779265 | 27.34427795 | 66.75566967 | 65.565281 | 61.7606383 | -0.7714416 | 5.702532174 | 0.00018 | 0.019451 |
| ENSG00000127824 | TUBA4A | 1075.594104 | 1255.953076 | 1381.144001 | 1642.565562 | 1914.12427 | 1864.85724 | -0.54603503 | 10.57275575 | 0.000184 | 0.01972 |
| ENSG00000154582 | ELOC | 104.3973745 | 75.88704677 | 100.6063056 | 152.3157533 | 125.401557 | 177.430986 | -0.70075495 | 6.948947367 | 0.000188 | 0.01982 |
| ENSG00000113583 | C5orf15 | 17.5668659 | 13.47107339 | 15.99382295 | 31.49739344 | 33.7374747 | 24.076181 | -0.92262822 | 4.554658274 | 0.000189 | 0.01982 |
| ENSG00000123131 | PRDX4 | 15.05731363 | 14.36914495 | 10.83452522 | 24.91584854 | 22.2794644 | 31.9271096 | -0.97315744 | 4.379923336 | 0.000191 | 0.01982 |
| ENSG00000099795 | NDUFB7 | 135.5158227 | 183.2065981 | 127.9505836 | 236.9356163 | 217.702195 | 238.144834 | -0.63158504 | 7.576182466 | 0.000192 | 0.01982 |
| ENSG00000091436 | MAP3K20 | 171.1514649 | 148.6308431 | 180.5754204 | 251.0389268 | 220.884976 | 272.68892 | -0.57598844 | 7.702983585 | 0.000197 | 0.020158 |
| ENSG00000008988 | RPS20 | 70.26746361 | 92.05233484 | 63.45936202 | 103.424277 | 125.401557 | 142.363505 | -0.71185317 | 6.647722383 | 0.0002 | 0.020321 |
| ENSG00000215030 | RPL13P12 | 91.84961314 | 123.4848394 | 94.41514838 | 149.9652016 | 152.136914 | 183.188334 | -0.64535763 | 7.059403966 | 0.000205 | 0.020575 |
| ENSG00000072422 | RHOBTB1 | 13.04967181 | 17.51239541 | 28.89206726 | 7.991875946 | 8.27522965 | 7.8509286 | 1.29225966 | 3.913228447 | 0.000208 | 0.020575 |
| ENSG00000198959 | TGM2 | 45.67385135 | 56.57850825 | 46.43367953 | 29.14684169 | 32.4643625 | 26.6931572 | 0.75921797 | 5.347461814 | 0.000208 | 0.020575 |
| ENSG00000198952 | SMG5 | 70.76937406 | 70.04958164 | 77.90539566 | 51.71213848 | 41.3761482 | 44.4885954 | 0.65989944 | 5.922288906 | 0.000212 | 0.020758 |
| ENSG00000183873 | SCN5A | 377.9385721 | 366.4131963 | 465.8845846 | 282.5363202 | 195.422731 | 284.72701 | 0.66306745 | 8.367223324 | 0.000215 | 0.020901 |
| ENSG00000182899 | RPL35A | 304.6596458 | 334.5316559 | 268.2834817 | 408.0557836 | 425.856049 | 539.097097 | -0.59681407 | 8.573358633 | 0.000218 | 0.020998 |
| ENSG00000166592 | RRAD | 80.30567269 | 94.29751374 | 63.9752918 | 99.19328381 | 173.143266 | 147.597458 | -0.8086401 | 6.784406683 | 0.00022 | 0.02102 |
| ENSG00000106633 | GCK | 28.10698544 | 48.49586421 | 62.42750248 | 20.68485539 | 20.369796 | 26.169762 | 1.04392603 | 5.15513342 | 0.000224 | 0.021226 |
| ENSG00000166123 | GPT2 | 18.57068681 | 6.735536696 | 16.50975272 | 3.760882798 | 3.81933676 | 5.75734764 | 1.62111841 | 3.360718683 | 0.000229 | 0.0215 |
| ENSG00000173457 | PPP1R14B | 17.5668659 | 19.75757431 | 14.9619634 | 26.32617959 | 41.3761482 | 32.9739001 | -0.92394277 | 4.709883496 | 0.000233 | 0.021712 |
| ENSG00000236391 | AC092573.2 | 461.757618 | 429.7272412 | 444.2155342 | 689.6518831 | 553.167274 | 679.890416 | -0.52661586 | 9.087544687 | 0.000238 | 0.02199 |
| ENSG00000249360 | CTD-2233C11.3 | 9.536298632 | 9.429751374 | 10.31859545 | 19.74463469 | 20.369796 | 18.3188334 | -0.99164696 | 3.952147566 | 0.000247 | 0.02264 |
| ENSG00000243199 | RP11-408P14.1 | 38.14519453 | 48.49586421 | 31.47171613 | 65.81544897 | 57.9266075 | 74.8455193 | -0.7463333 | 5.747958527 | 0.00025 | 0.022755 |
| ENSG00000033627 | ATP6V0A1 | 89.34006087 | 97.4407642 | 107.3133927 | 59.23390407 | 71.2942862 | 61.2372431 | 0.62157821 | 6.359356066 | 0.000277 | 0.024876 |
| ENSG00000182359 | KBTBD3 | 42.16047816 | 26.94214678 | 37.6628734 | 80.38886981 | 52.8341585 | 56.0032907 | -0.83646776 | 5.652092777 | 0.000278 | 0.024876 |
| ENSG00000160058 | BSDC1 | 42.16047816 | 40.86225595 | 38.69473294 | 20.21474504 | 15.9139032 | 30.3569239 | 0.85964712 | 5.02923145 | 0.000292 | 0.025952 |
| ENSG00000250479 | CHCHD10 | 102.3897327 | 115.4021954 | 86.16027202 | 130.2205669 | 176.326047 | 171.150243 | -0.64706278 | 7.032871823 | 0.000302 | 0.026549 |
| ENSG00000150401 | DCUN1D2 | 16.06113454 | 13.92010917 | 13.41417409 | 7.521765597 | 7.00211739 | 5.2339524 | 1.12506207 | 3.544878251 | 0.000333 | 0.028943 |
| ENSG00000225200 | AB019441.29 | 95.36298632 | 124.382911 | 101.6381652 | 144.3238774 | 166.141149 | 176.384196 | -0.59550253 | 7.082136759 | 0.000334 | 0.028943 |
| ENSG00000114737 | CISH | 20.57832863 | 45.35261375 | 39.72659249 | 10.81253805 | 21.0063522 | 15.7018572 | 1.17180977 | 4.740369183 | 0.000339 | 0.028981 |
| ENSG00000121671 | CRY2 | 23.58979135 | 24.69696788 | 24.76462908 | 13.63320014 | 14.0042348 | 12.0380905 | 0.88061066 | 4.318138577 | 0.000339 | 0.028981 |
| ENSG00000163584 | RPL22L1 | 64.74644861 | 68.7024743 | 63.45936202 | 82.26931121 | 105.031761 | 129.802019 | -0.68402726 | 6.433004101 | 0.000356 | 0.030135 |
| ENSG00000197444 | OGDHL | 279.5641231 | 286.4848275 | 317.2968101 | 211.0795471 | 221.521532 | 179.524567 | 0.52984344 | 7.968097519 | 0.000358 | 0.030135 |
| ENSG00000198034 | RPS4X | 355.8545121 | 357.4324807 | 341.5455094 | 510.5398399 | 455.13763 | 495.131897 | -0.470337 | 8.71491967 | 0.000363 | 0.030282 |
| ENSG00000235907 | RP4-580O19.2 | 21.08023908 | 14.36914495 | 17.54161227 | 28.20662099 | 32.4643625 | 34.5440858 | -0.84534697 | 4.67156949 | 0.000373 | 0.030959 |
| ENSG00000198795 | ZNF521 | 26.60125408 | 21.55371743 | 22.18498022 | 39.01915903 | 38.8299237 | 38.7312477 | -0.72972546 | 5.000052906 | 0.000383 | 0.031495 |
| ENSG00000073578 | SDHA | 9.536298632 | 22.00275321 | 10.83452522 | 7.521765597 | 1.90966838 | 3.66376668 | 1.63801052 | 3.410150726 | 0.000392 | 0.032071 |
| ENSG00000132341 | RAN | 95.36298632 | 78.13222567 | 85.64434224 | 113.7667046 | 135.586455 | 139.746529 | -0.58510484 | 6.764916281 | 0.000397 | 0.032181 |
| ENSG00000214975 | PPIAP29 | 22.08405999 | 22.00275321 | 15.47789318 | 28.20662099 | 38.8299237 | 42.3950144 | -0.86403409 | 4.854423407 | 0.000413 | 0.033308 |
| ENSG00000100209 | HSCB | 0.501910454 | 0 | 0.515929773 | 2.350551749 | 4.45589289 | 3.14037144 | -3.08034934 | 1.449877934 | 0.000423 | 0.033812 |
| ENSG00000164258 | NDUFS4 | 245.9361226 | 197.5757431 | 232.6843274 | 317.7945965 | 299.817936 | 366.376668 | -0.54263901 | 8.116525877 | 0.000426 | 0.033812 |
| ENSG00000225178 | RPSAP58 | 63.24071725 | 66.00825962 | 75.32574679 | 118.9379185 | 91.0275261 | 102.062072 | -0.61438973 | 6.443968672 | 0.000429 | 0.033812 |
| ENSG00000214753 | HNRNPUL2 | 17.5668659 | 25.14600366 | 19.08940158 | 36.19849693 | 31.1912502 | 40.8248287 | -0.80488881 | 4.872713043 | 0.000438 | 0.034284 |
| ENSG00000236801 | RPL24P8 | 55.71206043 | 82.17354769 | 65.00715134 | 104.3644977 | 93.5737506 | 125.091462 | -0.66957173 | 6.469331091 | 0.000458 | 0.035262 |
| ENSG00000161791 | FMNL3 | 68.25982179 | 72.74379632 | 67.07087043 | 55.47302128 | 33.1009186 | 35.5908763 | 0.73111557 | 5.827135574 | 0.000458 | 0.035262 |
| ENSG00000224948 | ATP5G1P8 | 81.81140406 | 80.37740457 | 87.70806133 | 105.7748287 | 132.403674 | 135.559367 | -0.57793675 | 6.709343277 | 0.000462 | 0.035262 |
| ENSG00000117298 | ECE1 | 61.73498588 | 75.88704677 | 68.61865975 | 47.95125568 | 43.2858166 | 41.3482239 | 0.63560569 | 5.851345416 | 0.000464 | 0.035262 |
| ENSG00000136238 | RAC1 | 89.84197133 | 65.55922384 | 86.67620179 | 103.424277 | 139.405792 | 139.223134 | -0.65717837 | 6.708985848 | 0.000466 | 0.035262 |
| ENSG00000112996 | MRPS30 | 18.57068681 | 18.41046697 | 21.15312067 | 37.13871763 | 33.7374747 | 29.3101334 | -0.78815408 | 4.768727087 | 0.00047 | 0.035389 |
| ENSG00000206418 | RAB12 | 111.4241209 | 107.3195514 | 109.3771118 | 139.6227739 | 162.958368 | 177.954382 | -0.54881312 | 7.082487194 | 0.000479 | 0.035429 |
| ENSG00000178741 | COX5A | 1177.481926 | 1044.008188 | 1132.981781 | 1437.59745 | 1689.41996 | 1513.65903 | -0.46800645 | 10.38068276 | 0.000479 | 0.035429 |
| ENSG00000228224 | NACAP1 | 55.21014998 | 43.10743485 | 44.36996044 | 65.81544897 | 75.1136229 | 82.6964479 | -0.64706478 | 5.950000176 | 0.00048 | 0.035429 |
| ENSG00000263232 | ATP5A1P3 | 59.72734407 | 84.86776237 | 79.45318497 | 94.49218031 | 130.494006 | 136.082762 | -0.68337515 | 6.617519896 | 0.000489 | 0.035827 |
| ENSG00000147010 | SH3KBP1 | 83.31713542 | 86.21486971 | 89.77178042 | 130.2205669 | 137.496123 | 113.053372 | -0.55245135 | 6.747583949 | 0.000494 | 0.035947 |
| ENSG00000181904 | C5orf24 | 27.10316453 | 23.34986055 | 27.86020772 | 36.19849693 | 45.195485 | 50.7693383 | -0.7521816 | 5.163107195 | 0.000499 | 0.036128 |
| ENSG00000142534 | RPS11 | 195.7450772 | 220.0275321 | 196.0533136 | 256.2101406 | 295.362043 | 338.113325 | -0.53873597 | 7.971762097 | 0.000515 | 0.036981 |
| ENSG00000115211 | EIF2B4 | 48.68531407 | 52.08815045 | 49.52925816 | 78.97853876 | 74.4770668 | 72.7519383 | -0.58947305 | 5.991747235 | 0.00052 | 0.036981 |
| ENSG00000225496 | AC104651.2 | 8.532477724 | 6.735536696 | 13.41417409 | 15.51364154 | 24.8256889 | 21.4592048 | -1.10091473 | 3.978505303 | 0.000521 | 0.036981 |

| | | | | | | | | | | | |
|---|---|---|---|---|---|---|---|---|---|---|---|
| ENSG00000213619 | NDUFS3 | 506.9295589 | 475.5288907 | 471.0438823 | 662.3854829 | 641.648576 | 667.328931 | -0.43985658 | 9.159050649 | 0.000538 | 0.037803 |
| ENSG00000216938 | RPL7P58 | 3.011462726 | 1.796143119 | 5.159297725 | 12.22286909 | 7.63867352 | 8.37432384 | -1.52970093 | 2.873285988 | 0.000539 | 0.037803 |
| ENSG00000125730 | C3 | 22.58597045 | 24.24793211 | 19.60533136 | 15.04353119 | 10.8214542 | 7.8509286 | 0.9635675 | 4.164919699 | 0.000552 | 0.038261 |
| ENSG00000107262 | BAG1 | 59.72734407 | 59.7217587 | 58.81599407 | 91.20140786 | 81.4791842 | 91.594167 | -0.570351 | 6.222491058 | 0.000552 | 0.038261 |
| ENSG00000175920 | DOK7 | 8.532477724 | 13.02203761 | 10.83452522 | 3.290772449 | 5.09244901 | 4.71055716 | 1.32547065 | 3.123184757 | 0.000571 | 0.039117 |
| ENSG00000218175 | AC016739.2 | 142.0406586 | 197.1267073 | 171.8046143 | 238.8160577 | 241.891328 | 271.642129 | -0.55690822 | 7.723983145 | 0.000575 | 0.039117 |
| ENSG00000117707 | PROX1 | 76.79229951 | 61.9669376 | 59.33192384 | 93.08184926 | 98.6661996 | 106.772629 | -0.59119145 | 6.385000058 | 0.000575 | 0.039117 |
| ENSG00000197879 | MYO1C | 84.82286678 | 105.5234082 | 95.96293769 | 64.87522827 | 57.9266075 | 67.5179859 | 0.58658373 | 6.334999884 | 0.000582 | 0.039376 |
| ENSG00000189186 | DCAF8L2 | 9.536298632 | 10.77685871 | 4.12743818 | 1.880441399 | 2.54622451 | 2.6169762 | 1.79364923 | 2.68224042 | 0.000589 | 0.039388 |
| ENSG00000205981 | DNAJC19 | 73.78083679 | 69.60054586 | 66.03901089 | 92.61173891 | 103.758649 | 116.193743 | -0.57573334 | 6.45609217 | 0.000592 | 0.039388 |
| ENSG00000204564 | C6orf136 | 24.09170181 | 22.90082477 | 24.76462908 | 36.19849693 | 34.3740308 | 54.9565002 | -0.81012027 | 5.077646108 | 0.000593 | 0.039388 |
| ENSG00000117643 | MAN1C1 | 19.07259726 | 17.96143119 | 21.66905045 | 13.16308979 | 6.36556127 | 9.42111432 | 0.9772114 | 3.986244112 | 0.000612 | 0.04026 |
| ENSG00000215464 | AP000354.2 | 16.56304499 | 22.00275321 | 13.93010386 | 27.73651064 | 35.6471431 | 30.8803191 | -0.82408064 | 4.659069341 | 0.000621 | 0.04026 |
| ENSG00000179988 | PSTK | 8.030567269 | 7.184572476 | 8.770806133 | 15.98375189 | 14.0042348 | 17.7954382 | -1.00188922 | 3.687611638 | 0.000622 | 0.04026 |
| ENSG00000075415 | SLC25A3 | 2527.621048 | 2417.608638 | 2598.738264 | 3280.900131 | 3509.33393 | 3301.57717 | -0.41974265 | 11.521633 | 0.000625 | 0.04026 |
| ENSG00000185088 | RPS27L | 3.011462726 | 7.633608255 | 8.254876361 | 11.75275874 | 14.6407909 | 19.3656239 | -1.25867786 | 3.540222934 | 0.000625 | 0.04026 |
| ENSG00000172586 | CHCHD1 | 31.11844817 | 36.82093394 | 27.34427795 | 52.65235918 | 44.5589289 | 56.5266859 | -0.68947006 | 5.408636354 | 0.000627 | 0.04026 |
| ENSG00000070087 | PFN2 | 72.77701588 | 63.76308072 | 62.94343225 | 96.84273206 | 108.851098 | 91.0707717 | -0.57001583 | 6.382966584 | 0.000635 | 0.040527 |
| ENSG00000219470 | RP3-337H4.6 | 128.4890763 | 162.5509523 | 108.861182 | 208.7289953 | 187.147501 | 206.217724 | -0.58888104 | 7.391950521 | 0.000641 | 0.04067 |
| ENSG00000108262 | GIT1 | 64.74644861 | 74.09090365 | 61.39564293 | 44.66048323 | 40.7395921 | 45.0119906 | 0.6167973 | 5.817058334 | 0.000654 | 0.041031 |
| ENSG00000159592 | GPBP1L1 | 51.6967768 | 35.4738266 | 61.91157271 | 76.15787667 | 77.6598474 | 88.4537955 | -0.70576038 | 6.043648306 | 0.000656 | 0.041031 |
| ENSG00000204396 | VWA7 | 8.030567269 | 8.082644035 | 11.350455 | 4.701103498 | 1.90966838 | 3.14037144 | 1.41907234 | 2.879189326 | 0.000657 | 0.041031 |
| ENSG00000254112 | KB-1205A7.1 | 40.15283635 | 38.61707706 | 40.75845203 | 47.48114533 | 68.7480617 | 85.8368193 | -0.75315283 | 5.762440475 | 0.00067 | 0.041565 |
| ENSG00000149428 | HYOU1 | 37.14137362 | 43.55647063 | 39.21066271 | 28.20662099 | 20.369796 | 23.5527858 | 0.72040604 | 5.057357629 | 0.000673 | 0.041565 |
| ENSG00000182446 | NPLOC4 | 38.64710498 | 30.9834688 | 59.33192384 | 27.26640029 | 21.0063522 | 19.3656239 | 0.91716635 | 5.08446979 | 0.000686 | 0.042115 |
| ENSG00000139112 | GABARAPL1 | 60.73116498 | 66.4572954 | 71.71423838 | 41.83982113 | 41.3761482 | 46.5821763 | 0.61354409 | 5.806823229 | 0.00069 | 0.042115 |
| ENSG00000168273 | SMIM4 | 41.15665726 | 53.88429357 | 34.05136499 | 59.70401442 | 75.7501791 | 74.8455193 | -0.69341236 | 5.841643152 | 0.000724 | 0.043951 |
| ENSG00000105953 | OGDH | 1653.794947 | 1258.198255 | 1815.55687 | 1209.59393 | 863.170108 | 1041.55653 | 0.60140969 | 10.35346778 | 0.000752 | 0.045416 |
| ENSG00000137692 | DCUN1D5 | 81.81140406 | 73.19283209 | 79.45318497 | 104.3644977 | 121.58222 | 114.623558 | -0.53646259 | 6.593624491 | 0.000756 | 0.045416 |
| ENSG00000134531 | EMP1 | 16.06113454 | 22.45178899 | 18.05754204 | 39.95937973 | 29.9181379 | 29.3101334 | -0.81067416 | 4.75237932 | 0.000764 | 0.045652 |
| ENSG00000163382 | NAXE | 53.20250816 | 43.10743485 | 55.20448566 | 78.03831807 | 77.0232913 | 73.2753336 | -0.59486411 | 6.002594267 | 0.000778 | 0.046291 |
| ENSG00000075886 | TUBA3D | 1.505731363 | 8.082644035 | 4.12743818 | 9.402206996 | 14.0042348 | 15.178462 | -1.44422322 | 3.259256868 | 0.000787 | 0.046566 |
| ENSG00000180957 | PITPNB | 31.62035862 | 35.4738266 | 39.72659249 | 19.74463469 | 22.2794644 | 22.5059953 | 0.72943383 | 4.892287901 | 0.000796 | 0.046807 |
| ENSG00000082641 | NFE2L1 | 750.3561292 | 738.2148219 | 758.4167656 | 561.781868 | 543.618932 | 575.734764 | 0.41819655 | 9.357307316 | 0.000799 | 0.046807 |
| ENSG00000143393 | PI4KB | 52.70059771 | 46.25068531 | 57.78413453 | 33.37783484 | 38.1933676 | 23.0293906 | 0.7326639 | 5.425524463 | 0.000809 | 0.047152 |
| ENSG00000141562 | NARF | 10.03820909 | 11.67493027 | 16.50975272 | 3.290772449 | 8.27522965 | 3.66376668 | 1.36823765 | 3.313212792 | 0.000824 | 0.047539 |
| ENSG00000197852 | FAM212B | 5.019104543 | 7.184572476 | 8.254876361 | 2.350551749 | 1.90966838 | 2.09358096 | 1.64472908 | 2.482246996 | 0.000824 | 0.047539 |
| ENSG00000163735 | CXCL5 | 4.517194089 | 4.041322018 | 3.611508408 | 6.581544897 | 14.0042348 | 11.5146953 | -1.35631957 | 3.027057639 | 0.000839 | 0.048154 |
| ENSG00000185834 | RPL12P4 | 11.04203 | 20.65564587 | 11.86638477 | 27.73651064 | 28.6450257 | 25.6463667 | -0.89413473 | 4.448623019 | 0.000846 | 0.048304 |
| ENSG00000087302 | C14orf166 | 127.9871659 | 126.1790541 | 120.211637 | 173.9408294 | 169.32393 | 178.477777 | -0.47891567 | 7.231035144 | 0.000856 | 0.048485 |
| ENSG00000165629 | ATP5C1 | 695.1459793 | 636.2836999 | 673.8042829 | 886.6281197 | 1043.95205 | 841.09615 | -0.46646015 | 9.63811376 | 0.000858 | 0.048485 |
| ENSG00000230807 | AC099535.4 | 3.51337318 | 2.245178899 | 3.611508408 | 7.991875946 | 7.00211739 | 9.94450956 | -1.41888377 | 2.734025581 | 0.000877 | 0.049319 |
| ENSG00000121898 | CPXM2 | 36.63946317 | 36.82093394 | 35.08322453 | 17.39408294 | 23.5525767 | 24.5995763 | 0.7377722 | 4.913529666 | 0.000883 | 0.049411 |
| ENSG00000204220 | PFDN6 | 13.55158227 | 19.30853853 | 11.350455 | 30.08706239 | 30.5546941 | 21.9826001 | -0.88566101 | 4.460860419 | 0.000897 | 0.049941 |
| ENSG00000159388 | BTG2 | 19.57450772 | 18.85950275 | 12.89824431 | 8.461986296 | 9.5483419 | 7.32753336 | 1.02674819 | 3.801256386 | 0.000905 | 0.049948 |
| ENSG00000142733 | MAP3K6 | 6.022925452 | 11.22589449 | 11.86638477 | 1.880441399 | 3.18278063 | 5.2339524 | 1.49852777 | 2.944952804 | 0.000913 | 0.049948 |
| ENSG00000175390 | EIF3F | 24.59361226 | 30.53443302 | 26.82834817 | 44.66048323 | 42.0127044 | 42.9184097 | -0.65814678 | 5.176790818 | 0.000914 | 0.049948 |
| ENSG00000170035 | UBE2E3 | 109.9183895 | 89.80715594 | 100.6063056 | 148.5548705 | 135.586455 | 144.980481 | -0.51764844 | 6.936010385 | 0.000918 | 0.049948 |
| ENSG00000150527 | CTAGE5 | 18.57068681 | 20.65564587 | 22.18498022 | 31.96750379 | 35.6471431 | 33.4972953 | -0.71359477 | 4.802861986 | 0.000919 | 0.049948 |

**Supplementary Table 7 Prediction of Stat5-regulated targeted genes in this study from the existing Cistromes data [46-49]**

| Target gene | Biosamples | MASC2 binding score | MASC2 Q value | DOI (Signaling Pathways Project Datasets)* | Ref |
|---|---|---|---|---|---|
| ATP5C1 | B lymphocytes, GM12878 cells | 185 | 1< E-05 | 10.1621/WdnDiLYKWM | [46] |
| ATP5G3 | Fibroblasts, MEFs | 142 | 1< E-05 | 10.1621/xXhMIGXpPo | [47] |
| | Mammary gland | 159 | 1< E-05 | 10.1621/w9eiy79IrX | [48] |
| | myeloblasts, K562 cells | 176 | 1< E-05 | 10.1621/WdnDiLYKWM | [46] |
| ATP5L | Mammary gland | 85, 92 | 1< E-05 | 10.1621/w9eiy79IrX | [48] |
| ATP5MK (USMG5) | B lymphocytes, GM12878 cells | 162 | 1< E-05 | 10.1621/WdnDiLYKWM | [46] |
| NDUFA2 | B lymphocytes, GM12878 cells | 121 | 1< E-05 | 10.1621/WdnDiLYKWM | [46] |
| | myeloblasts, K562 cells | 1319 | 1< E-05 | 10.1621/WdnDiLYKWM | [46] |
| | T lymphocytes, CD4+ | 95, 119 | 1< E-05 | 10.1621/lSZJ58eZyg | [49] |
| | Mammary gland | 106, 164, 176, | 1< E-05 | 10.1621/w9eiy79IrX | [48] |
| | Fibroblasts, MEFs | 374 | 1< E-05 | 10.1621/xXhMIGXpPo | [47] |
| NDUFAB1 | myeloblasts, K562 cells | 139 | 1< E-05 | 10.1621/WdnDiLYKWM | [46] |
| | B lymphocytes, GM12878 cells | 174 | 1< E-05 | 10.1621/WdnDiLYKWM | [46] |
| | Fibroblasts, MEFs | 145 | 1< E-05 | 10.1621/xXhMIGXpPo | [47] |
| | T lymphocytes, CD4+ | 77, 108 | 1< E-05 | 10.1621/lSZJ58eZyg | [49] |
| NDUFA1 | B lymphocytes, GM12878 cells | 73 | 1< E-05 | 10.1621/WdnDiLYKWM | [46] |
| NDUFS3 | Mammary gland | 111 | 1< E-05 | 10.1621/w9eiy79IrX | [48] |
| PYGB | Mammary gland | 158 | 1< E-05 | 10.1621/w9eiy79IrX | [48] |
| OGDH | Fibroblasts, MEFs | 101 | 1< E-05 | 10.1621/xXhMIGXpPo | [47] |
| MFN2 | Fibroblasts, MEFs | 103 | 1< E-05 | 10.1621/xXhMIGXpPo | [47] |
| | Mammary gland | 135 | 1< E-05 | 10.1621/w9eiy79IrX | [48] |
| | T lymphocytes, CD4+ | 149 | 1< E-05 | 10.1621/lSZJ58eZyg | [49] |
| GNA12 | B lymphocytes, GM12878 cells | 150 | 1< E-05 | 10.1621/WdnDiLYKWM | [46] |
| | myeloblasts, K562 cells | 446 | 1< E-05 | 10.1621/WdnDiLYKWM | [46] |
| GYS1 | Fibroblasts, MEFs | 138 | 1< E-05 | 10.1621/xXhMIGXpPo | [47] |
| SCN5A | Mammary gland | 224 | 1< E-05 | 10.1621/w9eiy79IrX | [48] |
| PER1 | Mammary gland | 102, 208, 252, 277,329,468 | 1< E-05 | 10.1621/w9eiy79IrX | [48] |
| | Fibroblasts, MEFs | 98 | 1< E-05 | 10.1621/xXhMIGXpPo | [47] |
| SLC6A6 | myeloblasts, K562 cells | 121 | 1< E-05 | 10.1621/WdnDiLYKWM | [46] |
| | Fibroblasts, MEFs | 93, 316 | 1< E-05 | 10.1621/xXhMIGXpPo | [47] |

***Signaling Pathways Project Datasets**: http://www.signalingpathways.org/index.jsf
